# Model-guided design of mammalian genetic programs

**DOI:** 10.1101/2020.09.30.320853

**Authors:** Joseph J. Muldoon, Viswajit Kandula, Mihe Hong, Patrick S. Donahue, Jonathan D. Boucher, Neda Bagheri, Joshua N. Leonard

## Abstract

Genetically engineering cells to perform customizable functions is an emerging frontier with numerous technological and translational applications. However, it remains challenging to systematically engineer mammalian cells to execute complex functions. To address this need, we developed a method enabling accurate genetic program design using high-performing genetic parts and predictive computational models. We built multi-functional proteins integrating both transcriptional and post-translational control, validated models for describing these mechanisms, implemented digital and analog processing, and effectively linked genetic circuits with sensors for multi-input evaluations. The functional modularity and compositional versatility of these parts enable one to satisfy a given design objective via multiple synonymous programs. Our approach empowers bioengineers to predictively design mammalian cellular functions that perform as expected even at high levels of biological complexity.

---

Early demonstrations of genetically engineering customized functions in mammalian cells indicate a vast potential to benefit applications including directed stem cell differentiation (*1, 2*) and cancer immunotherapy (*3*). In general, most applications require precise control of gene expression and the capability to sense and respond to external cues (*4-8*). Despite the growing availability of biological *parts* (such as libraries of promoters and regulatory proteins) that could be used to control cell states, assembling parts to compose customized genetic programs that function as intended remains a challenge, and it often requires iterative experimental tuning or down-selection to identify functional configurations. This highly empirical process limits both the scope of programs that one can feasibly compose and fine-tune and likely the performance of functional programs identified in this manner. Thus, the need for systematic and precise design processes represents a grand challenge in the field of mammalian synthetic biology.

Model-guided predictive design has been demonstrated in the composition of some cellular functions, including transcriptional logic in bacteria (*9*) as well as logical (*10*) and analog behaviors in yeast (*11*); however, this type of approach is less developed in mammalian systems. To date, transcription factors (TFs) based on zinc fingers (ZFs) (*12, 13*), transcription activator-like effectors (TALEs) (*14-17*), dCas9 (*18, 19*), and other proteins (*20*) have been used to implement transcriptional logic in mammalian cells. Some of these studies make use of protein splicing (*12, 14, 18*). Other studies have used RNA-binding proteins (*21*), proteases (*22, 23*), and synthetic protein-binding domains (*17*). Yet, none of these approaches currently enable the customized design of sophisticated mammalian cellular functions and prediction of circuit performance based only upon descriptions of the component parts. Associated challenges include the availability of appropriate parts (*24*), suitably descriptive models that support predictions using these parts (*25*), and computational and conceptual tools that facilitate the identification of designs that function robustly despite biological variability and crosstalk (*26-28*). In this study, we sought to address these challenges by developing a model-driven process that enables one to propose a tractable set of candidate circuits for construction and testing without needing empirical trial-and-error tuning. We validated this framework by employing it to implement a variety of functions including digital and analog information processing, and sense-and-respond behaviors.

## Biological parts for integrating transcriptional and post-translational control of gene expression

The strategy that we pursued for genetic program design was uniquely enabled by the COmposable Mammalian Elements of Transcription (COMET): a toolkit of TFs and promoters with tunable properties enabling precise and orthogonal control of gene expression (*13*). These TFs comprise a ZF DNA-binding domain and a functional domain, e.g., VP16 and VP64 are activation domains (AD) that with a ZF form an activator (ZFa). A protein including a ZF domain but lacking an AD functions as a competitive inhibitor of the cognate ZFa. Promoters in this library contain ZF binding sites arranged in different configurations (e.g., ZF1×6-C has six compactly arranged ZF1 sites). Each combination of a promoter and a ZFa (and potentially an inhibitor) confers a characteristic level of transcriptional activity (**Fig. S1A–D**), and as part of this prior work, we developed mathematical models to characterize these relationships (*13*). Here, we investigate whether these biological parts and descriptive computational tools can be adapted and applied to achieve predictive genetic program design.

Although COMET includes many parts for implementing transcriptional regulation, we hypothesized that complex genetic program design would be facilitated by introducing a mechanism for regulation at the post-translational level (**Fig. 1A**,**B**). To investigate this strategy, we evaluated new parts based on split inteins: complementary domains that fold and *trans*-splice to covalently ligate flanking domains (exteins) (*29*). We selected the split intein gp41-1 for its rapid splicing kinetics (*30*). To test an application of this mechanism, we appended an AD to the gp41-1 N-terminal fragment (intN) and a ZF to the C-terminal fragment (intC). These parts were used to construct an AND gate in which a reporter gene was induced only when both fragments were present (**Fig. 1C, Fig. S1E**), demonstrating that COMET-mediated gene expression can be adapted with splicing. We next incorporated this mechanism into our modeling framework by modifying ordinary differential equations from the original study (*13*), which concisely represent transcriptional regulation (**Materials and Methods**), and fitting newly introduced parameters to the data (**Fig. S1F**,**G**). We extended the model to describe parts in which split inteins were fused onto two types of inhibitors (**Fig. S1H**): ZF, which competes with ZFa for binding site occupancy in the promoter; and ZF fused to DsRed-Express2 (abbreviated as DsRed-ZF), which also reduces the cooperativity of ZFa-mediated RNA polymerase II (RNAPII) recruitment at multi-site promoters (*13*). Additionally, we introduced an R95K mutation to ablate the DsRed chromophore (*31*), yielding a non-fluorescent inhibitor we termed DsDed-ZF (**Fig. S1I**). The extended model accurately recapitulated the component dose-dependent performance of the AND gate (**Fig. 1C**), providing verification that this extension can describe split intein-based circuits.

**Fig. 1.**
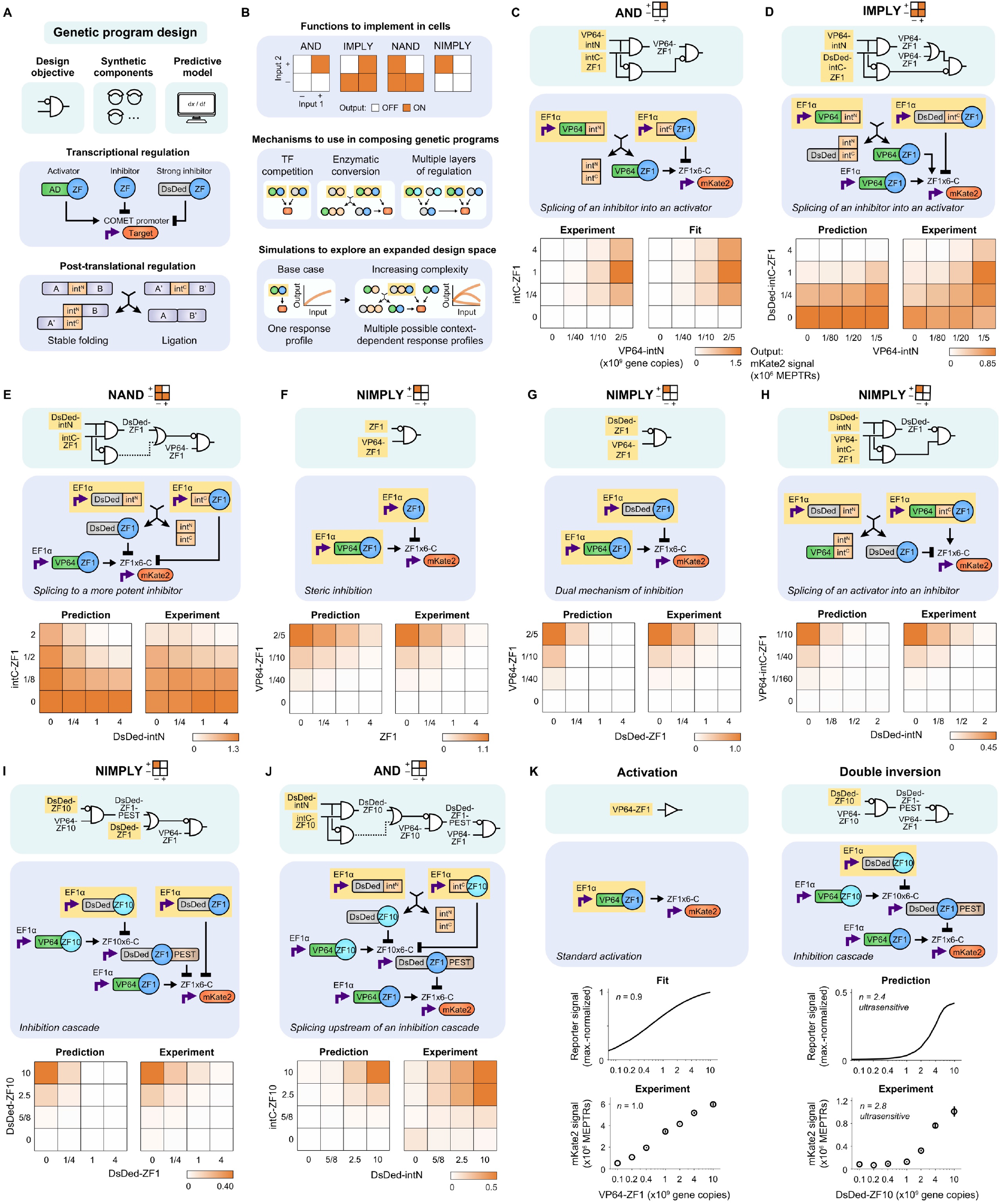
Logical evaluation is enabled by transcriptional and post-translational regulation. (**A**,**B**) Cartoons depict (**A**) the genetic components and (**B**) their arrangement and use in simulations to produce intended functions. Transcription is mediated by COMET TFs, which here are modified with split inteins to incorporate post-translational regulation via splicing. Genetic parts that carry out specified activities and that can be described mathematically should enable the predictive customization of cellular functions. In the schematics, circles are protein domains, arrows indicate splicing or regulation, yellow highlighting denotes the inputs, and the red node is the output. (**C–J**) A panel of logic gates was designed, simulated, and experimentally evaluated. Synthetic digital logic in cells is inherently analog, and component doses were selected to examine this behavior and underscore particular features (e.g., in **C**, reporter signal decreases at a high intC-ZF1 dose because intC-ZF1 inhibits ZFa-mediated transcription). In the electronic diagrams (teal background), lines denote splicing or regulation. Processes that have a modest effect within the dose range examined, and that because of fundamentally analog behavior do not carry out a fully digital function, are denoted by dotted lines. In the mechanistic diagrams (blue background), purple bent arrows are promoters, and black arrows indicate splicing and regulation. Yellow highlighting denotes the components for which dose is varied (in gene copies). Simulation and experimental results are presented in heatmaps that indicate how the two inputs affect reporter output (mKate2 signal in MEPTRs); color-coding denotes the mean reporter signal from three biological replicates (bar graphs in **Fig. S1L**, histograms in **Fig. S1M**), scaled by the maximum value in each heatmap. Simulations in **C** are from a fit to the data, and subsequent panels (**D–J**) are predictions. (**K**) Some of the motifs that were used in the gate designs confer sharp transitions in reporter output. For example, a standard activation dose response was not ultrasensitive, but layering two inhibitors in a cascade did produce ultrasensitivity (Hill coefficient *n* > 1). The downstream inhibitor is tagged with a PEST degron.

### Model-guided design of genetic programs

As a first test of the predictive capacity of the revised model, we simulated a panel of circuits that we hypothesized could carry out various logic operations (**Fig. S1J**). Our objective was to identify promising designs for specific functions, so we opted not to include additional model complexity that might be required to predict all aspects of circuit behavior (e.g., potential cell burden effects). Throughout, simulations employed a statistical model for gene expression variation, which we have previously shown to be important in accounting for the effect of cellular variation on how an engineered function is carried out across a cell population (*13, 32*) (**Materials and Methods, Fig. S1G**). From the panel, we selected several designs to test. First, to make an IMPLY gate, the AND gate was modified by appending DsDed to intC-ZF1 and co-expressing a VP64-ZF1 activator. Experimental outcomes (i.e., reporter readout across component doses) were consistent with the prediction that readout would be low only with DsDed-intC-ZF1 present in sufficient excess over its VP64-intN splicing partner to function as an inhibitor (**Fig. 1D**). To make a NAND gate, a DsDed-ZF1 inhibitor was split into DsDed-intN and intC-ZF1 and co-expressed with an activator. Outcomes were consistent with the prediction that readout would be low only with sufficient reconstitution of the inhibitor (**Fig. 1E**). These initial test cases demonstrate that model-guided design can identify effective topologies, as well as the precise relationship between input component levels and circuit output.

A versatile design framework would enable one to achieve a given performance objective via multiple circuits. We speculated that the combined properties of COMET and splicing-based extensions developed here might provide a sufficient basis for this capability. To investigate this possibility, we compared four designs for a NIMPLY gate, each of which utilizes a different mechanism (i.e., topology and/or choice of parts). The first two designs used inhibition mediated by ZF1 (**Fig. 1F**) or DsDed-ZF1 (**Fig. 1G**). The third design used splicing of an VP64-intC-ZF1 activator to a DsDed-ZF1 inhibitor, such that the readout would be high only with VP64-intC-ZF1 in sufficient excess of its splicing partner DsDed-intN (**Fig. 1H**). The fourth design used a double inversion cascade, in which an upstream inhibitor prevented a downstream inhibitor from acting on the reporter (**Fig. 1I**); this scenario represents a variation on a topology that was previously examined in bacteria (*33*) and later in mammalian cells with dCas9-TFs (*34*). All four designs produced NIMPLY as predicted. We next tested whether splicing could be combined with a cascade, and indeed we were able to build an AND gate by splitting the cascade’s upstream inhibitor into DsDed-intN and intC-ZF10 (**Fig. 1J**). Unlike standard ZFa-mediated activation, this activation via double inversion exhibited ultrasensitivity (Hill coefficient *n* = 2.8)—a signal transformation in which a small change in input yields a large change in output, and high output is produced only with sufficient input (**Fig. 1K, Fig. S1K**). Ultrasensitivity buffered the circuit against low inputs, such that the output remained low for input levels that in the standard activation case would have produced half-maximal activation.

Across the panel, five of the eight gates exhibited a goodness of prediction metric (comparing all simulated and observed outcomes, Q^2^) of at least 90%, indicating a high capacity for predicting dose response landscapes that had not been used in model training (**Fig. S1N**,**O**). Even for the gate with the lowest Q^2^ (IMPLY, **Fig. 1D, Fig. S1N**), the model correctly predicted the trend across most input dose combinations. Altogether, these results demonstrate the feasibility of model-guided design of logic gates in mammalian cells, and that the choice of parts and mechanism yields predictable performance characteristics.

### Compression of circuit design using functional modularity

A putative advantage of orthogonal parts like COMET TFs and promoters is that these parts may be used together without disrupting their functions. However, simply appending modules can lead to inefficient and cumbersome designs, and thus, one focus of our approach was achieving genetic compactness as well as performance. Enhancing compactness could eliminate potential failure modes and reduce cargo size for gene delivery vehicles. Genetic compression—reducing the number of components for a given specification—has been investigated by using recombinase-mediated DNA rearrangement (*35*) and by borrowing from a software engineering strategy to eliminate redundancy (*36*). Here, we sought to implement a previously unexplored form of *topological compaction* based on protein multi-tasking (**Fig. 2A**). We hypothesized that because our genetic parts operate through direct interactions without relying on long-range mechanisms such as chromatin modification, they might exhibit functional modularity, i.e., domains could be concatenated and retain their functions. This property would be of great utility by enabling the use of multi-tasking proteins to act at multiple promoters or in both transcriptional and post-translational roles, to execute multiple functions in an efficient fashion.

**Fig. 2.**
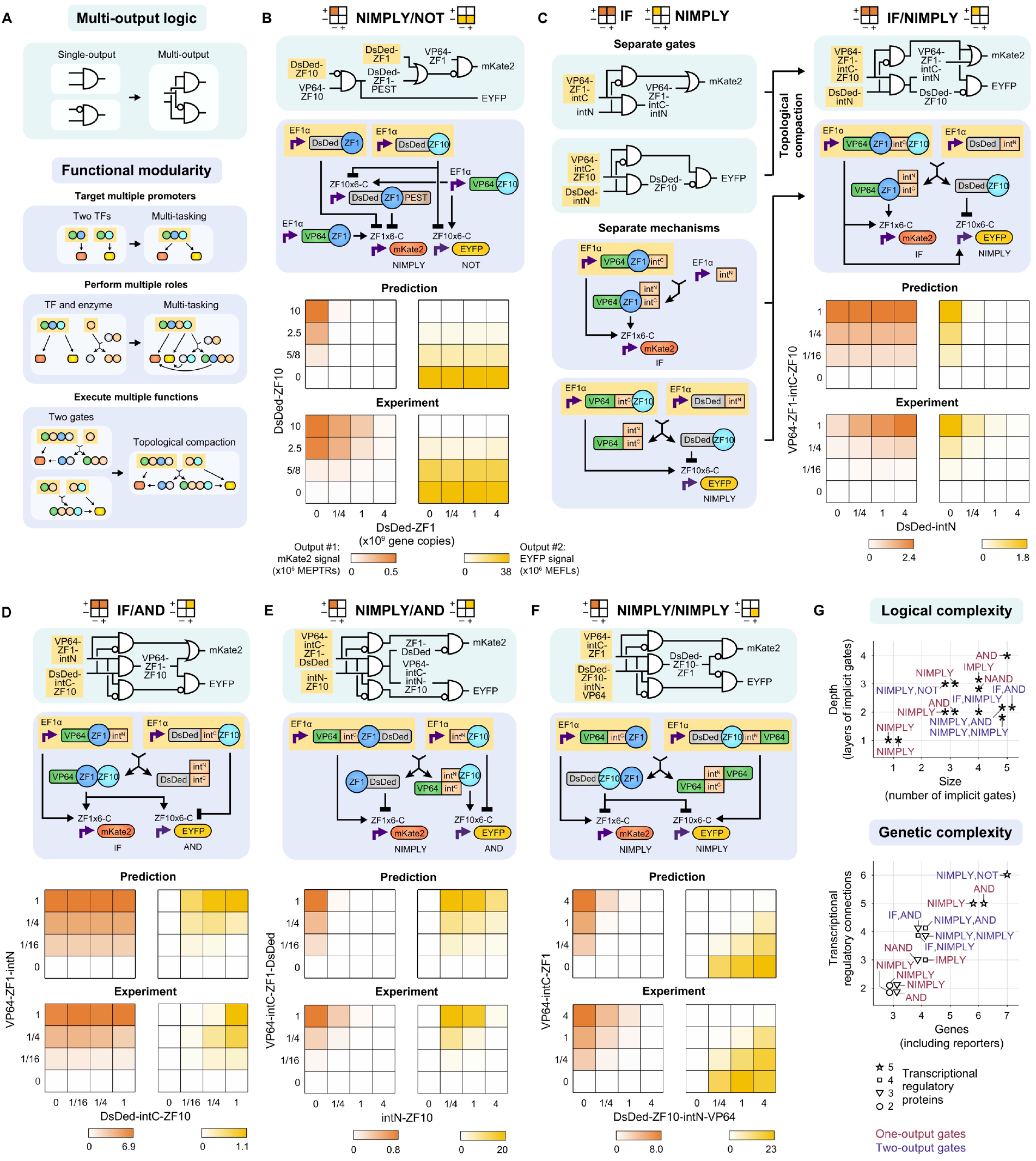
Compact multi-output logic is attained through functional modularity. (**A**) A strategy for multi-output logic is proposed by using multi-tasking proteins that retain the functions of their constituent domains. The cartoons depict the use of multiple DNA-binding domains on a TF to regulate multiple genes, the embedding of a split intein fragment within a functioning TF to enzymatically alter its activity, and the merging of features from multiple genetic programs to enable their compact simultaneous implementation. (**B–F**) A panel of multi-input-multi-output gates was designed, simulated, and experimentally evaluated. As an example, **C** is deconstructed to show how separate topologies containing proteins that have some domains in common and are amenable to the appending of additional domains can be compressed. In the plots, color-coding denotes the mean mKate2 and EYFP reporter signal from three biological replicates (bar graphs in **Fig. S2F**), scaled by the maximum value in each heatmap. (**G**) These plots summarize the complexity of the gates that were designed and validated in **Fig. 1** (red) and **Fig. 2** (purple), with complexity defined based on the size and depth of the circuits in the electronic diagrams (upper) or based on the numbers of genes, regulatory connections, and regulatory proteins employed (lower). The expanded toolkit of genetic parts and model-guided approach were successful for building circuits spanning a range of attributes, which suggests that this design process could be executed reliably for many future objectives.

We investigated whether functional modularity could enable the design of compact multi-input multi-output (MIMO) systems. Ultimately, this capability could support the encoding of sophisticated decision-making strategies in which cells take different actions in different situations. As a base case, we simply appended a NIMPLY gate and a NOT gate in a non-compact manner, and the combination functioned as expected (**Fig. 2B, Fig. S2A**). This success demonstrates the potential for composite functions, but it brings no efficiency relative to the individual gates. To test topological compaction, first, an IF/NIMPLY gate was proposed in which VP64-ZF1-intC-ZF10 would act as a bispecific activator (on two promoters) and interact with an inert DsDed-intN to produce a VP64-ZF1-intC/intN activator and a DsDed-ZF10 inhibitor (**Fig. 2C, Fig. S2B**). The second gate, IF/AND, used an activator and an inhibitor to produce a bispecific activator and an inert protein, through essentially the inverse mechanism of that in the IF/NIMPLY gate (**Fig. 2D, Fig. S2C**). Third, a NIMPLY/AND gate used a VP64-intC-ZF1-DsDed activator and an intN-ZF10 inhibitor to invert their respective activities (**Fig. 2E, Fig. S2D**). We hypothesized that the former protein would act as an activator, in that DsDed would not preclude VP64 from conferring activation. Lastly, a NIMPLY/NIMPLY gate used two activators to produce a bifunctional inhibitor and an inert protein (**Fig. 2F, Fig. S2E**). We note that if this circuit had used the same readout for both reporters it would be a XOR gate. Overall, the model predictions explained most of the variance in experimental outcomes, and several cases were in close agreement (≥90% Q^2^) (**Fig. S2F**,**G**). Minor deviations are potentially attributable to effects such as differences in stability for different proteins; however, we chose not to incorporate such effects into the model because increasing model complexity could lead to overfitting. Moreover, the choice to simplify the description of protein stability did not preclude model-guided identification of high-performing designs.

Notably, when we examined performance at the single-cell level, some population-level outcomes were driven by subpopulations of cells. In some circuits, subpopulations induced one reporter or the other, but not both, and thus population outcomes were driven by shifts in subpopulation frequencies (**Fig. S2A**,**D**,**E**). In other circuits, this task distribution was not apparent (**Fig. S2B**,**C**). Although neither behavior was an explicitly designed feature, both types of behavior were predicted by simulations. Altogether, the gates described in **Figs. 1**,**2** span a wide range of logical complexity (the number and the layers of implicit gates depicted in the electronic diagrams) and genetic complexity (the number of genes, regulatory connections, and regulatory proteins) (**Fig. 2G**). The successful development of these circuits without the need for additional tuning demonstrates that this framework may be well-suited to overcoming complexity-associated barriers with mammalian genetic program design.

### Implementation of analog signal processing

Although digital logic has many uses, biology also processes analog signals for many purposes, and we next examined whether our tools could be employed in this way. The first property that we sought to implement was ultrasensitivity, which is desirable in engineering sharp activation (*37, 38*) and is observed in the natural control of processes including cell growth, division, and apoptosis (*39*). The second property was bandpass concentration filtering, in which an output is produced only when the input falls within a certain range of magnitudes (*22, 40*). Bandpass concentration filtering is salient for both natural and synthetic spatial patterning (*41*). To develop a strategy for implementing these properties, we made use of existing mechanistic insights. Previously, we determined that ZFa-mediated activation is cooperative at the level of transcription initiation, and in comparing promoter architectures, maximal transcription increased with the number and compactness of binding sites (*13*). This COMET promoter feature confers high inducibility as well as a high sensitivity to inhibition by proteins that compete for DNA binding. We also deduced that TF *binding* to promoter is generally non-cooperative, and transcriptional output from such promoters is not inherently ultrasensitive to ZFa dose (*n* = 1). To construct systems that do exhibit ultrasensitivity (*n* > 1), we examined several strategies in which the output is inhibited only at low activator doses (**Fig. S3A–C**). The first design made use of the inhibition conferred by intC-ZF1 prior to splicing with a VP16-intN input (**Fig. S3B**). We reasoned that at low VP16-intN doses, intC-ZF-mediated inhibition would dominate, and at high doses, transactivation by reconstituted VP16-ZF would dominate. We also tested this concept with the addition of a DsDed-ZF to threshold the response by promoting relatively more inhibition at low input doses (**Fig. S3C**). However, the increase in ultrasensitivity was modest for these cases, apparently from insufficient inhibition at low activator doses due to decreased protein stability caused by appending the intC domain to the inhibitory ZF (**Fig. S1F**).

Compared to a ZFa base case (*n* = 1.0) (**Fig. 3A, Fig. S3D**), however, DsDed-ZF thresholding of ZFa-mediated activation did lead to an increase in the Hill coefficient (*n* = 1.9) (**Fig. 3B, Fig. S3E**). This outcome led us to consider a vehicular analogy: the circuits with DsDed-ZF are akin to applying the brake (inhibition) while applying the accelerator (activation), but a more effective approach might be to release the brake as the accelerator is applied. To realize this concept and circumvent choices that modulate protein stability, we used a chemically responsive COMET TF (RaZFa) in which rapamycin-induced heterodimerization domains FRB and FKBP are fused to an AD and a ZF, respectively. In the presence of rapamycin (which in this scenario is not an input, but rather an environmental species), heterodimerization of VP16-FRB and FKBP-ZF converts FKBP-ZF (brake) into RaZFa (accelerator), which induces the reporter. With rapamycin, the response of this circuit to VP16-FRB input indeed exhibited greater ultrasensitivity (*n* = 3.3), consistent with the prediction (**Fig. 3C, Fig. S3F**). Thus, in this system, ultrasensitivity can arise through cascades (**Fig. 1**) or reconstitution (**Fig. 3**), and neither mechanism requires the cooperativity in TF-DNA binding that is often associated with ultrasensitive responses.

**Fig. 3.**
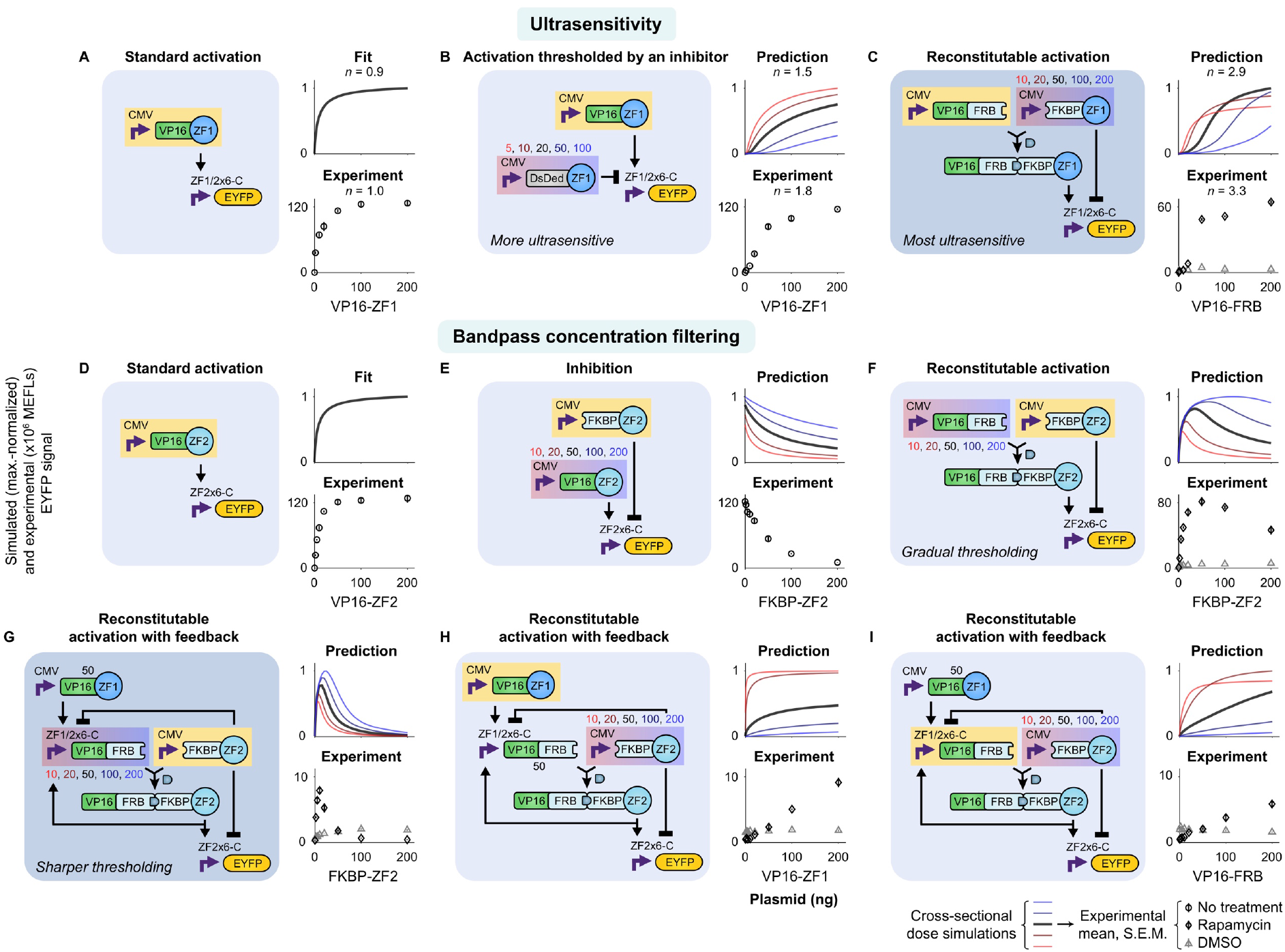
Analog behaviors are constructed by using TFs that play multiple roles. Reconstitutable TFs have dose response properties that are conducive to analog signal processing. Simulated and experimentally observed responses are shown relating to (**A–C**) ultrasensitivity and (**D–I**) bandpass concentration filtering. Several designs were evaluated for the ability to meet these objectives. To implement ultrasensitivity, the Hill coefficient (*n*) was most effectively increased through a strategy of removing an inhibitor in the process of producing an activator (**C**). To implement bandpass concentration filtering, a tighter upper threshold was best achieved through a similar strategy that also included additional regulation: moderate levels of FKBP-ZF act primarily to reconstitute RaZFa, and high levels of FKBP-ZF act to inhibit the reporter and VP16-FRB (**G**). Simulations in **A** and **D** are fitted to data, and the other panels are predictions. The prediction plots present simulations for how output gene expression varies with dose of the component highlighted in yellow; each plot includes a set of responses varying the component highlighted in red-to-blue gradation. Doses for the *x*-axes and above the varied component in the diagrams are in plasmid ng. Each experimental plot corresponds to the simulated condition with the dark line (for the middle dose of the varied component). The ZF1/2×6-C promoter has six partially overlapping ZF1 and ZF2 sites. DMSO is the vehicle for rapamycin, which is used here as an environmental species (not an input). The simulations with RaZFa correspond to conditions with rapamycin treatment. Experiment plots represent the mean and S.E.M. of EYFP reporter signal from three biological replicates (bar graphs in **Fig. S3D–L**).

We next investigated circuits to implement bandpass concentration filtering. Our strategy was to use mechanisms that inhibit reporter output only at high doses of activator input, and the predictions were based on a fitted ZFa base case (**Fig. 3D, Fig. S3G**). We hypothesized that although FKBP-ZF is necessary for RaZFa-mediated activation, excess FKBP-ZF would be inhibitory. We confirmed that FKBP-ZF acted as an inhibitor (**Fig. 3E, Fig. S3H**), and we implemented an RaZFa test circuit; the response to FKBP-ZF input showed a peak in output, but no sharp upper threshold, as predicted (**Fig. 3F, Fig. S3I**). Based on these results, we designed a new topology to achieve a sharper bandpass. Of the regulation within this design, the two paths of negative regulation from FKBP-ZF (and not the positive feedback from RaZFa) appeared to be most important for sharpening the bandpass (**Fig. S3M**). For the primary input to the bandpass, FKBP-ZF, we expected that at zero dose, there would be no activation; at moderate doses, there would be activation; and at high doses, excess FKBP-ZF would both decrease reconstitution (by inhibiting induction of VP16-FRB) and inhibit the reporter. The experimental outcomes closely matched the prediction of a bandpass with a sharp upper threshold (**Fig. 3G, Fig. S3J**). Furthermore, when VP16-ZF or VP16-FRB doses were varied, the responses were activating as predicted (**Fig. 3H–I, Fig. S3K–L**), demonstrating a predictive capacity across multiple inputs for the system. These results demonstrate that our parts and approach are suitable for designing analog behaviors, as well as digital logic gates.

### Integration of genetic circuits with sensors to build sense-and-respond functions

While the predictive design of genetic programs is a substantial technical advance, employing this capability to enable many potential applications will require integrating genetic circuits with native or synthetic parts that sense and modulate the state of the cell or its environment. A recurring challenge associated with this goal is level-matching the output of a sensor to the input requirements of a downstream circuit (*32, 42*). We investigated whether our designed circuits could overcome this challenge and be effectively linked to sensors without requiring laborious trial-and-error tuning. Simulations suggested that adding an upstream layer of signal processing (i.e., for sensing) should be feasible, since in the model, ZFa can be arranged in series without prohibitively driving up background or dampening induced signal (**Fig. S4A**).

We considered two classes of synthetic sensors (intracellular and transmembrane) for which we hypothesized that signaling (i.e., sensor output) could be coupled to COMET-based circuits. For the intracellular sensor, we built a new TF—ABA-ZFa, which is analogous to RaZFa—by fusing the abscisic acid (ABA)-binding domains PYL1 and ABI1 (*43*) to an AD and a ZF, respectively. For transmembrane sensing, we selected the modular extracellular sensor architecture (MESA)—a self-contained receptor and signal transduction system that transduces ligand binding into orthogonal regulation of target genes (*44, 45*). In this mechanism, ligand-mediated dimerization of two transmembrane proteins called the target chain (TC) and protease chain (PC) promotes PC-mediated proteolytic *trans*-cleavage of a TC-bound TF. We explored several strategies for building COMET-compatible MESA based on a recently reported improved MESA design (*46*) and the parts developed in the current study (**Fig. S4B–G**). The best performance was observed using rapalog-inducible COMET-MESA that release either ZFa for activating signaling or DsDed-ZF for inhibitory signaling (the latter represents a new function for MESA receptors) (**Fig. S4G**); the ZFa-releasing COMET-MESA receptor was carried forward. We observed that both sensors displayed excellent performance in terms of reporter induction upon ligand treatment (**Fig. 4A**,**B**). For ABA-ZF2a (ZF2a was selected for its potency stemming from cooperative transcriptional activation (*13*)), ligand-independent signal was unobservable, and induced signal was high, yielding perfect performance (**Fig. 4A**). For Rapa-MESA-ZF6a (ZF6a was also selected for its potency), the ligand-inducible fold difference in signal was ∼200x (**Fig. 4B**), which is several fold higher than was observed for recently reported receptors based on tTA (*46*), and also higher than the fold difference observed for a high-performing MESA that employs a distinct mechanism (*47*). Thus, Rapa-MESA-ZF6a is the highest performing MESA reported to date. Both sensors have a low off state and a high on state, apparently benefitting from the advantageous property of COMET promoter-based cooperativity.

**Fig. 4.**
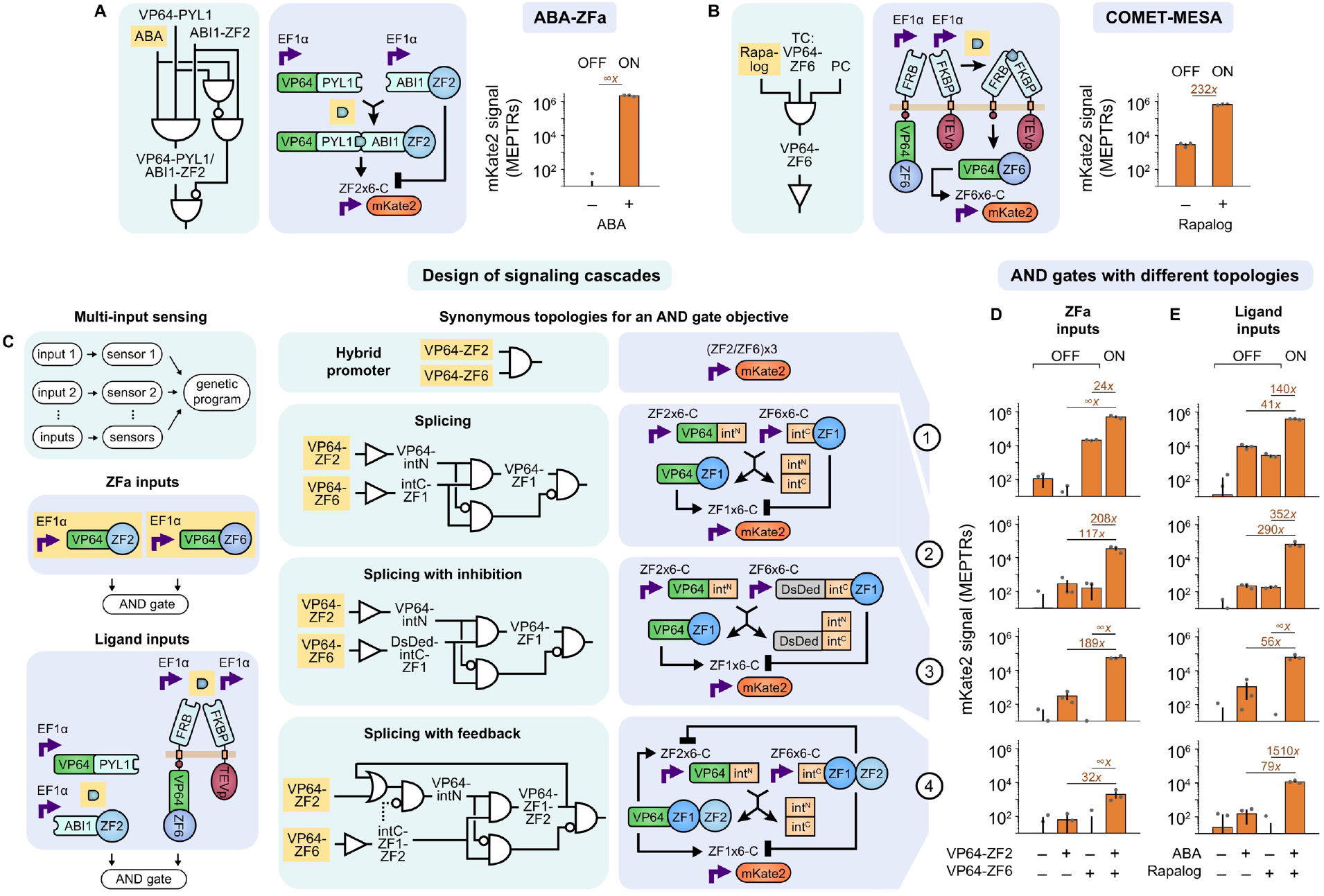
Sensors can be linked to genetic programs to make signaling cascades. MESA and COMET technologies can be combined to construct functional biosensors, and upstream biosensor output is well-matched to the requirements for downstream promoter input. (**A**,**B**) ABA-ZF2a and Rapa-MESA-ZF6a each exhibit ligand-inducible signaling (*p* = 2×10^−3^ and *p* = 1×10^−3^, respectively, one-tailed Welch’s unpaired *t*-test). EtOH is the vehicle for both ligands. For MESA, the TC contains an FRB ectodomain and intracellular COMET TF, and the PC contains an FKBP ectodomain and intracellular TEV protease (TEVp). Each receptor chain contains an FGFR4 transmembrane domain. (**C–E**) Validated sensors were applied to implement multi-input sensing. AND logic was selected as a design goal, and four synonymous topologies—those that are intended to achieve the same goal through different mechanisms—were proposed and evaluated. For each input type (two columns for upstream ZFa or ligand sensing) and topology (four rows), reporter signal with two inputs differed from that with either or no input (*p* < 2×10^−16^ in each case, three-factor ANOVA and Tukey’s HSD test), indicating successful AND gate outcomes. Topologies 2–4 displayed negligible background signal (comparable to the signal with only the reporter present, ∼10^1^–10^2^ MEPTRs, **Fig. S4H**), despite involving multi-layer signaling which can be a potential source of leak. The (ZF2/ZF6)x3 promoter has three pairs of alternating ZF2 and ZF6 sites. Bar graphs represent the mean, S.E.M., and values of mKate2 reporter signal from three biological replicates (depicted as dots; near-zero values are below the log-scaled *y*-axis lower limit). The numbers above bar pairs are the fold difference, and a fold difference of ∞ indicates that the denominator signal is less than or equal to zero.

We carried forward the two validated sensors and examined whether downstream circuits comprising genetic parts and designed topologies from this study could be seamlessly linked with the new input layer. To this end, we designed a panel of four synonymous topologies that implement AND logic through different mechanisms (**Fig. 4C**): 1) a hybrid promoter with alternating TF sites (based on a similar architecture from the original COMET study (*13*)), 2) splicing (as in **Fig. 1C**), 3) splicing with DsDed (as in **Figs. 1D, 2D** for tighter inhibition), and 4) and splicing with feedback (as in **Fig. 3G–I**). All four topologies exhibited AND behavior when tested using ZFa as inputs (**Fig. 4D**), demonstrating the versatility for attaining a given objective in multiple ways. Moreover, when coupled to ligand-activated sensors, these circuits still conferred AND behavior, and performance was maintained (i.e., fold induction with two ligands remained much greater than with each ligand individually) in carrying out this more complex sensing function (**Fig. 4E**). A comparison across the designs provides some insights. The hybrid promoter in topology 1 was high-performing, and the splicing topologies in 2–4 generally yielded improvement over 1, despite the additional regulatory layer, by reducing the output generated from either single input alone to near reporter-only background (**Fig. S4H**, shown with linear scaling). Of the topologies examined, 2 and 3 were the most effective at producing a high output when both inputs were present and low output when either input was present alone. These results demonstrate that genetic programs can be designed by a predictive model-driven process, and then these programs can be readily linked to different classes of sensors to implement high-performing sensing and processing functions.

## Discussion

We developed an approach for accurate genetic program design by engineering new parts that combine transcriptional and post-translational control and validating a computational modeling framework. The experimental observations closely matched simulations, even in scenarios employing new proteins (including those with many domains) and new topologies (including those with many interacting parts), demonstrating a high predictive capacity across a range of complexity (**Fig. 2G**). Since the mechanisms employed for binding, splicing, activation, and inhibition can be described by concise formalisms (**Materials and Methods**), no fundamental revamping (i.e., changing the underlying representation or granularity) of our original descriptive model was needed to enable predictions. Furthermore, no trial-and-error (e.g., empirical tuning of designs or substitution of parts) was needed to arrive at the specified design goals, which streamlined the design-build-test-learn cycle. We understand this to be possible because once the base case parts were characterized, no additional parameterization was needed to simulate how the parts would function when combined in new designs. Lastly, even though a relatively small set of protein domains was utilized, we were able to combine the domains in many ways; a concise library was sufficient to produce the wide variety of behaviors observed.

This study benefited from insights that could facilitate future genetic circuit design efforts. Key strategies that enabled sophisticated design included the use of antagonistic bifunctionality (*48*), in which a component can exert opposing effects on a target gene depending on the other components in the circuit (**Fig. S4I**), and functional modularity, which enabled multiple activities to be combined in individual proteins (**Fig. S4J**). Sophisticated design was also enabled through the use of split genetic parts, including those that splice or dimerize. Split parts are conducive to encoding both digital (**Figs. 1**,**2**) and analog (**Fig. 3**) functions. Split parts also shift some of the regulation from the transcriptional level to the post-translational level (i.e., protein-protein interactions), which could increase the speed of signal processing. Another benefit of split parts relates to circumventing cargo limitations of gene delivery vehicles, in that a large program that does not fit in one vector could be distributed across multiple vectors (*49*), in such a way that the parts interact to reconstitute the program only in cells receiving all of the vectors. Finally, we found that seamless level-matching could be achieved with multi-layer circuits due to the potency of COMET TFs at cognate promoters, and in particular, the fusion of such a TF onto MESA resulted in the highest-performing version of this receptor to date (**Fig. 4B**,**E**).

Altogether, these attributes and insights, in combination with the many ways in which components can be arranged to regulate each other, greatly expand the mammalian genetic program design space. In our current system, one can propose and formulate models for candidate designs based on principles for how the functionally modular parts operate (**Materials and Methods**) and then evaluate in silico outcomes. In the future, it should be possible to further automate this process by using software to sweep large combinatorial spaces and identify candidates that satisfy specified performance objectives. Such advances could further speed up the design process and broaden the scope of possible circuits and behaviors beyond those accessible solely by intuition. The new components and quantitative approaches developed here should enable bioengineers to build customized cellular functions for applications ranging from fundamental research to biotechnology and medicine.

## Supporting information

Plasmid Maps

## ACKNOWLEDGEMENTS

We acknowledge the services of the Northwestern University (NU) Flow Cytometry Core Facility and NUSeq Core Facility. We thank Taylor Dolberg for assistance with experiments; Katelyn Dray, Tae-Eun Kim, Joseph Draut, and Siyuan Feng for some of the cloning in this study; Katelyn Dray, Everett Allchin, and Christopher Coleman for collaboration on the computational models; and members of the Leonard Lab and Bagheri Lab for helpful discussions.

## Author contributions

Conceptualization, J.J.M., V.K., M.H., P.S.D., J.N.L.; Methodology, J.J.M., V.K., M.H., P.S.D., J.D.B.; Software, J.J.M., M.H., J.D.B.; Validation, J.J.M., V.K., P.S.D., J.D.B.; Formal Analysis, J.J.M., M.H.; Investigation, J.J.M., V.K., M.H., P.S.D., J.D.B.; Resources, J.J.M., V.K., M.H., P.S.D., J.D.B., J.N.L.; Data Curation, J.J.M.; Writing – Original Draft, J.J.M., J.N.L.; Writing – Review & Editing, J.J.M., V.K., M.H., P.S.D., J.D.B., N.B., J.N.L.; Visualization, J.J.M.; Supervision, N.B., J.N.L.; Project Administration, J.J.M., J.N.L.; Funding Acquisition, J.J.M., V.K., M.H. P.S.D., N.B., J.N.L.

## Data and materials availability

Plasmid maps are provided as supplementary material, and plasmids will be made available through Addgene. Code is provided as supplementary material and at https://github.com/leonardlab/GeneticPrograms under an open source license (*50*). Source data are available as supplementary material and from the corresponding author upon reasonable request.

## SUPPLEMENTARY MATERIALS

Materials and Methods, Figs. S1 to S4, Tables S1 to S2, References, Source_data.xlsx, Source_code.zip, Plasmid_maps.zip

