## Supplementary material for "Model-guided design of mammalian genetic programs": Source Code: Code_files.docx

| **File name** | **Description** |
| --- | --- |
| Z.mat | Population heterogeneity matrix |
| Params.m | Parameters values |
| ExampleRun.m | Example for running the AND gate model with the population model |

| **Figure and panel** | **File name** |
| --- | --- |
| **Figure 1** | |
| C | model_AND.m |
| D | model_IMPLY.m |
| E | model_NAND.m |
| F | model_NIMPLY_steric.m |
| G | model_NIMPLY_dual.m |
| H | model_NIMPLY_splicing.m |
| I | model_NIMPLY_cascade.m |
| J | model_AND_cascade.m |
| K | model_Activation.m |
| K | model_DoubleInversion.m |
| **Figure 2** | |
| B | model_NIMPLY_NOT.m |
| C | model_IF_NIMPLY.m |
| D | model_IF_AND.m |
| E | model_NIMPLY_AND.m |
| F | model_NIMPLY_NIMPLY.m |
| **Figure 3** | |
| C, F | model_RaZFa.m |
| E | model_FKBP.m |
| G, H, I | model_RaZFaFB.m |
| **Figure 4** | |
| S4A | model_OneLayer.m |
| S4A | model_TwoLayer.m |
| S4A | model_ThreeLayer.m |
