## Supplementary material for "Model-guided design of mammalian genetic programs": Source Code: ExampleRun.m

```
function matsim = ExampleRun(Z)
%Example code for running the AND gate model with the population model


%Plasmid ng
dose_intCZF1 = 10;
dose_ADintN  = 10;

%Number of cells (number of rows in Z)
nc = size(Z, 1);

%Initialize
matsimc = zeros(nc, 1);


%For each cell
for c = 1:nc
    
    %Simulate AND model
    sim = model_AND(dose_intCZF1, dose_ADintN, Z(c, :));
    
    %Store the last time point for the last variable (Reporter protein)
    matsimc(c) = sim(end, end);
end

%Average across the population
matsim = mean(matsimc);


end
```
