## Supplementary material for "Model-guided design of mammalian genetic programs": Source Code: model_Activation.m

```
function [sim, T] = model_Activation(dose_ZF1a_1, z)
%Fig. 1K


%Retrieve parameters
[p, T] = Params;

%Inducible promoter
d.Rep1 = 21.1 / 200; %~4e9 gene copies

%Initial values
IV = zeros(4, 1);


%Model
[~, sim] = ode15s(@(t, y, options)[ 
    
    % Y1 ZF1a RNA (INPUT)
      z(1)      * p.ktxEF1a * dose_ZF1a_1 ...
    - p.kdegR   * y(1)
    
    % Y2 ZF1a protein
      p.ktl     * y(1) ...
    - p.kdegZFP * y(2)
    
    % Y3 Reporter RNA
      z(2)      * p.ktxZF * d.Rep1^0.5 * (p.b1 + p.m1E64 * p.w1E64 * y(2)) ./ (1 + p.w1E64 * y(2)) ...
    - p.kdegR   * y(3)
    
    % Y4 Reporter protein
      p.ktl     * y(3) ...
    - p.kdegRep * y(4)
    
    ], T, IV);


end
```
