## Supplementary material for "Model-guided design of mammalian genetic programs": Source Code: model_AND.m

```
function [sim, T] = model_AND(dose_intCZF1, dose_ADintN, z)
%Fig. 1C


%Retrieve parameters
[p, T] = Params;

%Inducible promoter
d.Rep1 = 21.1 / 200; %~4e9 gene copies

%Function for ZF-mediated inhibition
function out = fI1(A, I)
    out = (p.b1 + p.m1E64 * p.w1E64 * A) ...
        ./ (1 + p.w1E64 * A + p.w1E64 * I);
end

%Initial values
IV = zeros(7, 1);


%Model
[~, sim] = ode15s(@(t, y, options)[ 
    
    % Y1 AD-intN RNA (INPUT)
      z(1)       * p.ktxEF1a * dose_ADintN ...
    - p.kdegR    * y(1)
    
    % Y2 AD-intN protein
      p.ktl      * y(1)         ...
    - p.rec      * y(2) .* y(4) ...
    - p.kdegZFP  * y(2)
    
    % Y3 intC-ZF1 RNA (INPUT)
      z(2)       * p.ktxEF1a * dose_intCZF1  ...
    - p.kdegR    * y(3)
    
    % Y4 intC-ZF1 protein
      p.ktl      * y(3)         ...
    - p.rec      * y(2) .* y(4) ...
    - p.kdegintC * y(4)
   
    % Y5 AD-ZF1 reconstituted protein
      p.rec      * y(2) .* y(4) ...
    - p.kdegZFP  * y(5)
    
    % Y6 Reporter RNA
      z(3)       * p.ktxZF * d.Rep1^0.5 * fI1(y(5), y(4)) ...
    - p.kdegR    * y(6)
    
    % Y7 Reporter protein
      p.ktl      * y(6) ...
    - p.kdegRep  * y(7)
    
    ], T, IV);


end
```
