## Supplementary material for "Model-guided design of mammalian genetic programs": Source Code: model_AND_cascade.m

```
function [sim, T] = model_AND_cascade(dose_intCZF10, dose_DsDedintN, z)
%Fig. 1J


%Retrieve parameters
[p, T] = Params;

%Plasmid ng-equivalents of constitutive genes
dose_ZF10a = 21.1;
dose_ZF1a  = 2.11;

%Inducible promoters
d.DsDedZF1PEST = 21.1 / 200; %~4e9 gene copies
d.Rep1         = 21.1 / 200; %~4e9 gene copies

%Functions for DsDed-ZF-mediated inhibition
function out = fIDI10(A, DI, I)
    out = (p.b10 + p.w10E64 * A .* max(min(((((p.wr10E64 * DI) ...
        ./ (p.w10E64 * A + eps)) - p.l) * (1 - p.m10E64)) ...
        ./ (p.u - p.l) + p.m10E64, p.m10E64), 1)) ...
        ./ (1 + p.w10E64 * A + p.wr10E64 * DI + p.w10E64 * I);
end
function out = fIDI1(A, R)
    out = (p.b1 + p.w1E64 * A .* max(min(((((p.wr1E64 * R) ...
        ./ (p.w1E64 * A + eps)) - p.l) * (1 - p.m1E64)) ...
        ./ (p.u - p.l) + p.m1E64, p.m1E64), 1)) ...
        ./ (1 + p.w1E64 * A + p.wr1E64 * R);
end

%Initial values
IV = zeros(14, 1);


%Model
[~, sim] = ode15s(@(t, y, options)[ 
    
    % Y1 DsDed-intN RNA (INPUT)
      z(1)       * p.ktxEF1a * dose_DsDedintN ...
    - p.kdegR    * y(1)
    
    % Y2 DsDed-intN protein
      p.ktl      * y(1)         ...
    - p.rec      * y(2) .* y(4) ...
    - p.kdegZFP  * y(2)
    
    % Y3 intC-ZF10 RNA (INPUT)
      z(2)       * p.ktxEF1a * dose_intCZF10 ...
    - p.kdegR    * y(3)
    
    % Y4 intC-ZF10 protein
      p.ktl      * y(3)         ...
    - p.rec      * y(2) .* y(4) ...
    - p.kdegintC * y(4)
    
    % Y5 DsDed-ZF10 protein (reconstituted)
      p.rec      * y(2) .* y(4) ...
    - p.kdegZFP  * y(5)
    
    % Y6 intC/intN
      p.rec      * y(2) .* y(4) ...
    - p.kdegintC * y(6)

    % Y7 ZF10a RNA 
      z(3)       * p.ktxEF1a * dose_ZF10a ...
    - p.kdegR    * y(7)
    
    % Y8 ZF10a protein
      p.ktl      * y(7) ...
    - p.kdegZFP  * y(8)
    
    % Y9 ZF1a RNA
      z(4)       * p.ktxEF1a * dose_ZF1a ...     
    - p.kdegR    * y(9)
    
    % Y10 ZF1a protein
      p.ktl      * y(9) ...
    - p.kdegZFP  * y(10)
    
    % Y11 DsDed-ZF1-PEST RNA
      z(5)       * p.ktxZF * d.DsDedZF1PEST^0.5 * fIDI10(y(8), y(5), y(4)) ...
    - p.kdegR    * y(11)
    
    % Y12 DsDed-ZF1-PEST protein
      p.ktl          * y(11) ...
    - p.kdegZFP_PEST * y(12)
    
    % Y13 Reporter RNA
      z(6)      * p.ktxZF * d.Rep1^0.5 * fIDI1(y(10), y(12)) ...
    - p.kdegR   * y(13)
    
    % Y14 Reporter protein
      p.ktl     * y(13) ...
    - p.kdegRep * y(14)
    
    ], T, IV);


end
```
