## Supplementary material for "Model-guided design of mammalian genetic programs": Source Code: model_DoubleInversion.m

```
function [sim, T] = model_DoubleInversion(dose_DsDedZF10, z)
%Fig. 1K


%Retrieve parameters
[p, T] = Params;

%Plasmid ng-equivalents for constitutive promoters
d.ZF10a     = 21.1; %~4e9 gene copies
d.DsDedZF10 = 21.1; %~4e9 gene copies
d.ZF1a      = 2.11; %~4e8 gene copies

%Inducible promoters
d.Rep1         = 21.1 / 200; %~4e9 gene copies
d.DsDedZF1PEST = 21.1 / 200; %~4e9 gene copies

%Functions for DsDed-ZF-mediated inhibition
function out = fDI10(A, DI)
    out = (p.b10 + p.w10E64 * A .* max(min(((((p.wr10E64 * DI) ...
        ./ (p.w10E64 * A + eps)) - p.l) * (1 - p.m10E64)) ...
        ./ (p.u - p.l) + p.m10E64, p.m10E64), 1)) ...
        ./ (1 + p.w10E64 * A + p.wr10E64 * DI);
end
function out = fDI1(A, DI)
    out = (p.b1 + p.w1E64 * A .* max(min(((((p.wr1E64 * DI) ...
        ./ (p.w1E64 * A + eps)) - p.l) * (1 - p.m1E64)) ...
        ./ (p.u - p.l) + p.m1E64, p.m1E64), 1)) ...
        ./ (1 + p.w1E64 * A + p.wr1E64 * DI);
end

%Initial values
IV = zeros(10, 1);


%Model
[~, sim] = ode15s(@(t, y, options)[ 
    
    % Y1 ZF10a RNA 
      z(1)       * p.ktxEF1a * d.ZF10a ...
    - p.kdegR    * y(1)
    
    % Y2 ZF10a protein
      p.ktl      * y(1) ...
    - p.kdegZFP  * y(2)
    
    % Y3 DsDed-ZF10 RNA (INPUT)
      z(2)       * p.ktxEF1a * dose_DsDedZF10 ...     
    - p.kdegR    * y(3)
    
    % Y4 DsDed-ZF10 protein
      p.ktl      * y(3) ...
    - p.kdegZFP  * y(4)
    
    % Y5 ZF1a RNA
      z(3)       * p.ktxEF1a * d.ZF1a ...     
    - p.kdegR    * y(5)
    
    % Y6 ZF1a protein
      p.ktl      * y(5) ...
    - p.kdegZFP  * y(6)
    
    % Y7 DsDed-ZF1-PEST RNA
      z(4)       * p.ktxZF * d.DsDedZF1PEST^0.5 * fDI10(y(2), y(4)) ...
    - p.kdegR    * y(7)
    
    % Y8 DsDed-ZF1-PEST protein
      p.ktl          * y(7) ...
    - p.kdegZFP_PEST * y(8)
    
    % Y9 Reporter RNA
      z(5)      * p.ktxZF * d.Rep1^0.5 * fDI1(y(6), y(8)) ...
    - p.kdegR   * y(9)
    
    % Y10 Reporter protein
      p.ktl     * y(9) ...
    - p.kdegRep * y(10)
    
    ], T, IV);


end
```
