## Supplementary material for "Model-guided design of mammalian genetic programs": Source Code: model_IMPLY.m

```
function [sim, T] = model_IMPLY(dose_DsDedintCZF1, dose_ADintN, z)
%Fig. 1D


%Retrieve parameters
[p, T] = params_general_200116;

%Inducible promoter
d.Rep1 = 21.1 / 200; %~4e9 gene copies

%Constitutive promoter
d.ZF1a = 1.06; %~2e8 gene copies

%Function for DsDed-ZF-mediated inhibition
function out = fDI1(A, DI)
    out = (p.b1 + p.w1E64 * A .* max(min(((((p.wr1E64 * DI) ...
        ./ (p.w1E64 * A + eps)) - p.l) * (1 - p.m1E64)) ...
        ./ (p.u - p.l) + p.m1E64, p.m1E64), 1)) ...
        ./ (1 + p.w1E64 * A + p.wr1E64 * DI);
end

%Initial values
IV = zeros(9, 1);


%Model
[~, sim] = ode15s(@(t, y, options)[ 
    
    % Y1 ZF1a RNA
      z(1)       * p.ktxEF1a * d.ZF1a ...
    - p.kdegR    * y(1)
    
    % Y2 ZF1a protein
      p.ktl      * y(1) ...
    - p.kdegZFP  * y(2)
    
    % Y3 AD-intN RNA (INPUT)
      z(2)       * p.ktxEF1a * dose_ADintN ...
    - p.kdegR    * y(3)
    
    % Y4 AD-intN protein
      p.ktl      * y(3) ...
    - p.rec      * y(4) .* y(6) ...
    - p.kdegZFP  * y(4)
    
    % Y5 DsDed-intC-ZF1 RNA (INPUT)
      z(3)       * p.ktxEF1a * dose_DsDedintCZF1 ...
    - p.kdegR    * y(5)
    
    % Y6 DsDed-intC-ZF1 protein
      p.ktl      * y(5) ...
    - p.rec      * y(4) .* y(6) ...
    - p.kdegintC * y(6)
    
    % Y7 AD-ZF1 reconstituted protein
      p.rec      * y(4) .* y(6) ...
    - p.kdegZFP  * y(7)
    
    % Y8 Reporter RNA
      z(4)      * p.ktxZF * d.Rep1^0.5 * fDI1(y(2) + y(7), y(6)) ...
    - p.kdegR    * y(8)
    
    % Y9 Reporter protein
      p.ktl      * y(8) ...
    - p.kdegRep  * y(9)
    
    ], T, IV);


end
```
