## Supplementary material for "Model-guided design of mammalian genetic programs": Source Code: model_NAND.m

```
function [sim, T] = model_NAND(dose_intCZF1, dose_DsDedintN, z)
%Fig. 1E


%Retrieve parameters
[p, T] = Params;

%Plasmid ng-equivalents of constitutive gene
dose_VP64ZF1 = 2.11;

%Inducible promoter
d.Rep1 = 21.1 / 200; %~4e9 gene copies

%Function for DsDed-ZF inhibition
function out = fRI1(A, DI, I)
    out = (p.b1 + p.w1E64 * A .* max(min(((((p.wr1E64 * DI) ...
        ./ (p.w1E64 * A + eps)) - p.l) * (1 - p.m1E64)) ...
        ./ (p.u - p.l) + p.m1E64, p.m1E64), 1)) ...
        ./ (1 + p.w1E64 * A + p.wr1E64 * DI + p.w1E64 * I);
end

%Initial values
IV = zeros(10, 1);


%Model
[~, sim] = ode15s(@(t, y, options)[ 
    
    % Y1 DsDed-intN RNA
      z(1)       * p.ktxEF1a * dose_DsDedintN ...
    - p.kdegR    * y(1)
    
    % Y2 DsDed-intN protein
      p.ktl      * y(1)         ...
    - p.rec      * y(2) .* y(4) ...
    - p.kdegZFP  * y(2)
    
    % Y3 intC-ZF1 RNA
      z(2)       * p.ktxEF1a * dose_intCZF1 ...
    - p.kdegR    * y(3)
    
    % Y4 intC-ZF1 protein
      p.ktl      * y(3)         ...
    - p.rec      * y(2) .* y(4) ...
    - p.kdegintC * y(4)
    
    % Y5 DsDed-ZF1 protein (reconstituted)
      p.rec      * y(2) .* y(4) ...
    - p.kdegZFP  * y(5)
    
    % Y6 intC/intN protein (reconstituted)
      p.rec      * y(2) .* y(4) ...
    - p.kdegintC * y(6)
    
    % Y7 VP64-ZF1 RNA
      z(3)       * p.ktxEF1a * dose_VP64ZF1 ...
    - p.kdegR    * y(7)
    
    % Y8 VP64-ZF1 protein
      p.ktl      * y(7) ...
    - p.kdegZFP  * y(8)
    
    % Y9 Reporter RNA
      z(4)       * p.ktxZF * d.Rep1^0.5 * fRI1(y(8), y(5), y(4)) ...
    - p.kdegR    * y(9)
    
    % Y10 Reporter protein
      p.ktl      * y(9) ...
    - p.kdegRep  * y(10)
    
    ], T, IV);


end
```
