## Supplementary material for "Model-guided design of mammalian genetic programs": Source Code: model_NIMPLY_dual.m

```
function [sim, T] = model_NIMPLY_dual(dose_ZF1a, dose_DsDedZF1, z)
%Fig. 1G


%Retrieve parameters
[p, T] = Params;

%Inducible promoter
d.Rep1 = 21.1 / 200; %~4e9 gene copies

%Function for DsDed-ZF-mediated inhibition
function out = fDI1(A, DI)
    out = (p.b1 + p.w1E64 * A .* max(min(((((p.wr1E64 * DI) ...
        ./ (p.w1E64 * A + eps)) - p.l) * (1 - p.m1E64)) ...
        ./ (p.u - p.l) + p.m1E64, p.m1E64), 1)) ...
        ./ (1 + p.w1E64 * A + p.wr1E64 * DI);
end

%Initial values
IV = zeros(6, 1);


%Model
[~, sim] = ode15s(@(t, y, options)[ 
    
    % Y1 ZF1a RNA (INPUT)
      z(1)       * p.ktxEF1a * dose_ZF1a ...
    - p.kdegR    * y(1)
    
    % Y2 ZF1a protein
      p.ktl      * y(1) ...
    - p.kdegZFP  * y(2)
    
    % Y3 DsDed-ZF1 RNA (INPUT)
      z(2)       * p.ktxEF1a * dose_DsDedZF1 ...   
    - p.kdegR    * y(3)
    
    % Y4 DsDed-ZF1 protein
      p.ktl      * y(3) ...
    - p.kdegZFP  * y(4)
    
    % Y5 Reporter RNA
      z(3)       * p.ktxZF * d.Rep1^0.5 * fDI1(y(2), y(4)) ...
    - p.kdegR    * y(5)
    
    % Y6 Reporter protein
      p.ktl      * y(5) ...
    - p.kdegRep  * y(6)
    
    ], T, IV);


end
```
