## Supplementary material for "Model-guided design of mammalian genetic programs": Source Code: model_IF_AND.m

```
function [sim, T] = model_IF_AND(dose_VP64ZF1intN, dose_DsDedintCZF10, z)
%Fig. 2D


%Retrieve parameters
[p, T] = Params;

%Inducible promoters
d.Rep1  = 21.1 / 200; %~4e9 gene copies
d.Rep10 = 21.1 / 200; %~4e9 gene copies

%Initial values
IV = zeros(10, 1);


%Model
[~, sim] = ode15s(@(t, y, options)[ 
    
    % Y1 VP64-ZF1-intN RNA
      z(1)       * p.ktxEF1a * dose_VP64ZF1intN  ...
    - p.kdegR    * y(1)
    
    % Y2 VP64-ZF1-intN protein
      p.ktl      * y(1) ...
    - p.rec      * y(2) .* y(4) ...
    - p.kdegZFP  * y(2)
    
    % Y3 DsDed-intC-ZF10 RNA
      z(2)       * p.ktxEF1a * dose_DsDedintCZF10  ...
    - p.kdegR    * y(3)
    
    % Y4 DsDed-intC-ZF10 protein 
      p.ktl      * y(3) ...
    - p.rec      * y(2) .* y(4) ...
    - p.kdegintC * y(4)
    
    % Y5 DsDed-intC/intN protein (reconstituted)
      p.rec      * y(2) .* y(4) ...
    - p.kdegintC * y(5)
    
    % Y6 VP64-ZF1-ZF10 protein (reconstituted)
      p.rec      * y(2) .* y(4) ...
    - p.kdegZFP  * y(6)
    
    % Y7 Reporter1 RNA
      z(3) * p.ktxZF * d.Rep1^0.5 ...
          * (p.b1 + p.m1E64 * p.w1E64 * (y(2) + y(6))) ...
          ./ (1 + p.w1E64 * (y(2) + y(6))) ...
    - p.kdegR   * y(7)
    
    % Y8 Reporter1 protein
      p.ktl     * y(7)  ...
    - p.kdegRep * y(8)
    
    % Y9 Reporter10 RNA
      z(4) * p.ktxZF * d.Rep10^0.5 ...
          * ((p.b10 + p.w10E64 * y(6) .* max(min(((((p.wr10E64 * y(4)) ...
          ./ (eps + p.w10E64 * y(6))) - p.l) * (1 - p.m10E64)) ...
          ./ (p.u - p.l) + p.m10E64, p.m10E64), 1)) ...
          ./ (1 + p.w10E64 * y(6) + p.wr10E64 * y(4))) ...
    - p.kdegR   * y(9)
    
    % Y10 Reporter10 protein
      p.ktl     * y(9)  ...
    - p.kdegRep * y(10)
    
    ], T, IV);


end
```
