## Supplementary material for "Model-guided design of mammalian genetic programs": Source Code: model_NIMPLY_NOT.m

```
function [sim, T] = model_NIMPLY_NOT(dose_DsDedZF10, dose_DsDedZF1, z)
%Fig. 2B


%Retrieve parameters
[p, T] = Params;

%Plasmid ng-equivalents of constitutive genes
dose_VP64ZF10 = 21.1; %~4e9 gene copies
dose_VP64ZF1  = 2.11; %~4e8 gene copies

%Inducible promoters
d.Rep1         = 21.1 / 200; %~4e9 gene copies
d.Rep10        = 21.1 / 200; %~4e9 gene copies
d.DsDedZF1PEST = 21.1 / 200; %~4e9 gene copies

%Initial values
IV = zeros(14, 1);


%Model
[~, sim] = ode15s(@(t, y, options)[ 
    
    % Y1 DsDed-ZF10 RNA
      z(1)       * p.ktxEF1a * dose_DsDedZF10 ...
    - p.kdegR    * y(1)
    
    % Y2 DsDed-ZF10 protein
      p.ktl      * y(1)  ...
    - p.kdegZFP  * y(2)
    
    % Y3 DsDed-ZF1 RNA
      z(2)       * p.ktxEF1a * dose_DsDedZF1 ...
    - p.kdegR    * y(3)
    
    % Y4 DsDed-ZF1 protein
      p.ktl      * y(3)  ...
    - p.kdegZFP  * y(4)
    
    % Y5 VP64-ZF10 RNA
      z(3)       * p.ktxEF1a * dose_VP64ZF10 ...
    - p.kdegR    * y(5)
    
    % Y6 VP64-ZF10 protein
      p.ktl      * y(5)  ...
    - p.kdegZFP  * y(6)
    
    % Y7 VP64-ZF1 RNA
      z(4)       * p.ktxEF1a * dose_VP64ZF1 ...
    - p.kdegR    * y(7)
    
    % Y8 VP64-ZF1 protein
      p.ktl      * y(7)  ...
    - p.kdegZFP  * y(8)
    
    % Y9 DsDed-ZF1-PEST RNA
      z(5) * p.ktxZF * d.DsDedZF1PEST^0.5 ...
          * ((p.b10 + p.w10E64 * y(6) .* max(min(((((p.wr10E64 * y(2)) ...
          ./ (eps + p.w10E64 * y(6))) - p.l) * (1 - p.m10E64)) ...
          ./ (p.u - p.l) + p.m10E64, p.m10E64), 1)) ...
          ./ (1 + p.w10E64 * y(6) + p.wr10E64 * y(2))) ...
    - p.kdegR   * y(9)
    
    % Y10 DsDed-ZF1-PEST protein
      p.ktl          * y(9)  ...
    - p.kdegZFP_PEST * y(10)
    
    % Y11 Reporter1 RNA
      z(6) * p.ktxZF * d.Rep1^0.5 ...
          * ((p.b1 + p.w1E64 * y(8) .* max(min(((((p.wr1E64 * (y(4) + y(10))) ...
          ./ (eps + p.w1E64 * y(8))) - p.l) * (1 - p.m1E64)) ...
          ./ (p.u - p.l) + p.m1E64, p.m1E64), 1)) ...
          ./ (1 + p.w1E64 * y(8) + p.wr1E64 * (y(4) + y(10)))) ...
    - p.kdegR   * y(11)
    
    % Y12 Reporter1 protein
      p.ktl     * y(11)  ...
    - p.kdegRep * y(12)
    
    % Y13 Reporter10 RNA
    z(7) * p.ktxZF * d.Rep10^0.5 ...
        * ((p.b10 + p.w10E64 * y(6) .* max(min(((((p.wr10E64 * y(2)) ...
        ./ (eps + p.w10E64 * y(6))) - p.l) * (1 - p.m10E64)) ...
        ./ (p.u - p.l) + p.m10E64, p.m10E64), 1)) ...
        ./ (1 + p.w10E64 * y(6) + p.wr10E64 * y(2))) ...
    - p.kdegR   * y(13)
    
    % Y14 Reporter10 protein
      p.ktl     * y(13)  ...
    - p.kdegRep * y(14)
    
    ], T, IV);


end
```
