## Supplementary material for "Model-guided design of mammalian genetic programs": Source Code: model_FKBP.m

```
function sim = model_FKBP(dose_FKBPZF2, dose_VP16ZF2, z)
%Fig. 3E


%Genetic program
%  Promoter  Gene
%  ********  ****
%  CMV       VP16-ZF2
%  CMV       FKBP-ZF2
%  ZF2x6-C   EYFP


%Retrieve parameters
[p, T] = Params;

%Inducible promoter
d.Rep = 200 / 200;  %200 ng

%Initial values
IV = zeros(6, 1);

%Function for FKBP-mediated inhibition
    function f = fAI(b, m, wA, wI, A, I)
        f = (b + m*wA*A) ./ (1 + wA*A + wI*I);
    end


%Model
[~, sim] = ode15s(@(t, y, options)[ 
    
    %Y1 VP16-ZF2 RNA
      z(1)      * p.ktxCMV * dose_VP16ZF2 ...
    - p.kdegR   * y(1)
    
    %Y2 VP16-ZF2 protein
      p.ktl     * y(1) ...
    - p.kdegZFP * y(2)    
    
    %Y3 FKBP-ZF2 RNA
      z(2)      * p.ktxCMV * dose_FKBPZF2 ...
    - p.kdegR   * y(3)
    
    %Y4 FKBP-ZF2 protein
      p.ktl     * y(3) ...
    - p.kdegZFP * y(4)   
    
    %Y5 Reporter RNA
      z(3)      * p.ktxZF * d.Rep^0.5 * fAI(p.b2, p.m2, p.w2, p.w2, y(2), y(4)) ...
    - p.kdegR   * y(5)
    
    %Y6 Reporter protein
      p.ktl     * y(5) ...
    - p.kdegRep * y(6)
    
    ], T, IV);


end
```
