## Supplementary material for "Model-guided design of mammalian genetic programs": Source Code: model_RaZFa.m

```
function sim = model_RaZFa(dose_FKBPZF1, dose_VP16FRB, z)
%Fig. 3C,F


%Genetic program
%  Promoter  Gene
%  ********  ****
%  CMV       FKBP-ZF2
%  CMV       VP16-FRB
%  ZF2x6-C   EYFP

%Retrieve parameters
[p, T] = Params;

%Inducible promoter
d.Rep = 200 / 200; %200 ng

%Time of ligand treatment
tau = 12;

%Function for activation and inhibition
    function f = fAI(b, m, wA, wI, A, I)
        f = (b + m*wA*A) ./ (1 + wA*A + wI*I);
    end

%Initial values
IV = zeros(7, 1);


%Model
[~, sim] = ode15s(@(t, y, options)[ 
    
    %Y1 VP16-FRB RNA
      z(1)       * p.ktxCMV * dose_VP16FRB ...
    - p.kdegR    * y(1)
    
    %Y2 VP16-FRB protein
      p.ktl      * y(1)                               ...
    - p.rec      * y(2) .* y(4) .* heaviside(t - tau) ... 
    - p.kdegZFP  * y(2)
    
    %Y3 FKBP-ZF1 RNA
      z(2)       * p.ktxCMV * dose_FKBPZF1 ...
    - p.kdegR    * y(3)
    
    %Y4 FKBP-ZF1 protein
      p.ktl      * y(3)                               ...
    - p.rec      * y(2) .* y(4) .* heaviside(t - tau) ... 
    - p.kdegZFP  * y(4)
    
    %Y5 VP16-FRB/Rapamycin/FKBP-ZF1 protein (reconstituted)
      p.rec      * y(2) .* y(4) .* heaviside(t - tau) ... 
    - p.kdegZFP  * y(5)
    
    %Y6 Reporter RNA
      z(3)       * p.ktxZF * d.Rep^0.5 * fAI(p.b2, p.m2, p.w2, p.w2, y(5), y(4)) ...
    - p.kdegR    * y(6)
    
    %Y7 Reporter protein
      p.ktl      * y(6) ...
    - p.kdegRep  * y(7)
    
    ], T, IV);


end
```
