## Supplementary material for "Model-guided design of mammalian genetic programs": Source Code: model_RaZFaFB.m

```
function sim = model_RaZFaFB(dose_FKBPZF2, dose_VP16FRB, z)
%Fig. 3G,H,I


%Genetic program
%  Promoter   Gene
%  ********   ****
%  CMV        VP16-ZF1
%  ZF1/2x6-C  VP16-FRB
%  CMV        FKBP-ZF2
%  ZF2x6-C    EYFP

%Retrieve parameters
[p, T] = Params;

%Plasmid ng for constitutive gene
dose_VP16ZF1 = 50;

%Inducible promoters
d.VP16FRB = dose_VP16FRB / 200;  %varied
d.Rep     = 200 / 200;           %200 ng

%Time of ligand treatment
tau = 12;

%Function for activation and inhibition
    function f = fAI(b, m, wA, wI, A, I)
        f = (b + m*wA*A) ./ (1 + wA*A + wI*I);
    end
%Function for multiple activation and inhibition
    function f = fAAI(b, m1, m2, w1A, w2A, wI, A1, A2, I)
        f = (b + m1*w1A*A1 + m2*w2A*A2) ./ (1 + w1A*A1 + w2A*A2 + wI*I);
    end

%Initial values
IV = zeros(9, 1);


%Model
[~, sim] = ode15s(@(t, y, options)[ 
    
    %Y1 VP16-ZF1 RNA
      z(1)       * p.ktxCMV * dose_VP16ZF1 ...
    - p.kdegR    * y(1)
    
    %Y2 VP16-ZF1 protein
      p.ktl      * y(1) ...
    - p.kdegZFP  * y(2)
    
    %Y3 VP16-FRB RNA
      z(2)       * p.ktxZF * d.VP16FRB^0.5 * fAAI(p.bH, p.m1, p.m2, p.w1H, p.w2H, p.w2H, y(2), y(7), y(6)) ...
    - p.kdegR    * y(3)
    
    %Y4 VP16-FRB protein
      p.ktl      * y(3)                               ...
    - p.rec      * y(4) .* y(6) .* heaviside(t - tau) ... 
    - p.kdegZFP  * y(4)
    
    %Y5 FKBP-ZF2 RNA
      z(3)       * p.ktxCMV * dose_FKBPZF2 ...
    - p.kdegR    * y(5)
    
    %Y6 FKBP-ZF2 protein
      p.ktl      * y(5)                               ...
    - p.rec      * y(4) .* y(6) .* heaviside(t - tau) ... 
    - p.kdegZFP  * y(6)    
    
    %Y7 VP16-FRB/Rapamycin/FKBP-ZF2 protein (reconstituted)
      p.rec      * y(4) .* y(6) .* heaviside(t - tau) ... 
    - p.kdegZFP  * y(7)
    
    %Y8 Reporter RNA
      z(4)       * p.ktxZF * d.Rep^0.5 * fAI(p.b2, p.m2, p.w2, p.w2, y(7), y(6)) ...
    - p.kdegR    * y(8)
    
    %Y9 Reporter protein
      p.ktl      * y(8) ...
    - p.kdegRep  * y(9)

    ], T, IV);


end
```
