## Supplementary material for "Model-guided design of mammalian genetic programs": Source Code: model_TwoLayer.m

```
function [sim, T] = model_TwoLayer(dose_ZF1a_1, dose_ZF1a_2, z)


%Retrieve parameters
[p, T] = Params;

%Inducible promoters
d.ZF1a_2 = dose_ZF1a_2 / 200;  %varied
d.Rep1   = 21.1 / 200;         %~4e9 gene copies

%Initial values
IV = zeros(6, 1);


%Model
[~, sim] = ode15s(@(t, y, options)[ 
    
    % Y1 ZF1a RNA (INPUT)
      z(1)       * p.ktxEF1a * dose_ZF1a_1 ...
    - p.kdegR    * y(1)
    
    % Y2 ZF1a protein
      p.ktl      * y(1) ...
    - p.kdegZFP  * y(2)

    % Y3 ZF1a RNA (INPUT)
      z(2)       * p.ktxZF * d.ZF1a_2^0.5 * (p.b1 + p.m1E64 * p.w1E64 * y(2)) ./ (1 + p.w1E64 * y(2)) ...
    - p.kdegR    * y(3)
    
    % Y4 ZF1a protein
      p.ktl      * y(3) ...
    - p.kdegZFP  * y(4)
    
    % Y5 Reporter RNA
      z(3)       * p.ktxZF * d.Rep1^0.5 * (p.b1 + p.m1E64 * p.w1E64 * y(4)) ./ (1 + p.w1E64 * y(4)) ...
    - p.kdegR    * y(5)
    
    % Y6 Reporter protein
      p.ktl      * y(5) ...
    - p.kdegRep  * y(6)
    
    ], T, IV);


end
```
