## Supplementary material for "Model-guided design of mammalian genetic programs": Source Code: LICENSE.rtf

Copyright (c) 2020 Joseph J. Muldoon

Permission is hereby granted, free of charge, to any person obtaining a copy of this software and associated documentation
files (the "Software"), to deal in the Software without restriction, including without limitation the rights to use, copy,
modify, merge, publish, distribute, sublicense, and/or sell copies of the Software, and to permit persons to whom the Software
is furnished to do so, subject to the following conditions:

The above copyright notice and this permission notice shall be included in all copies or substantial portions of the Software.

THE SOFTWARE IS PROVIDED "AS IS", WITHOUT WARRANTY OF ANY KIND, EXPRESS OR IMPLIED, INCLUDING BUT NOT LIMITED TO THE WARRANTIES
OF MERCHANTABILITY, FITNESS FOR A PARTICULAR PURPOSE AND NONINFRINGEMENT. IN NO EVENT SHALL THE AUTHORS OR COPYRIGHT HOLDERS BE
LIABLE FOR ANY CLAIM, DAMAGES OR OTHER LIABILITY, WHETHER IN AN ACTION OF CONTRACT, TORT OR OTHERWISE, ARISING FROM, OUT OF OR
IN CONNECTION WITH THE SOFTWARE OR THE USE OR OTHER DEALINGS IN THE SOFTWARE.
