## Supplementary material for "Model-guided design of mammalian genetic programs": Source Code: Params.m

```
function [p, T] = Params
%This function is called by model files.


%Timecourse (start and end time in hours)
T = [0, 42];

%Split inteins
p.rec      = 0.34;
p.kdegintC = 1.3;

%CMV VP16-based TFs
p.b1  = 0.08; 
p.m1  = 33;
p.w1  = 0.036;
p.b2  = 0.25;
p.m2  = 54;
p.b10 = 0.01;
p.w10 = p.w1;

%EF1a VP64-based TFs
p.m1E64  = 52;
p.w1E64  = 0.192;
p.m10E64 = p.m1E64;
p.w10E64 = p.w1E64;

%TF at a ZF2x6-C promoter
p.w2 = 0.082;

%TFs at a ZF1/2x6-C promoter
p.w1H  = 0.072;
p.w2H  = 0.170;
p.bH   = 0.08;
p.wr1H = 4 * p.w1H;
p.wr2H = 4 * p.w2H;

%CMV DsDed-ZFs
p.wr1  = 4 * p.w1;
p.wr10 = 4 * p.w10;
p.l    = 4 * 0;
p.u    = 4 * 1.5;

%EF1a VP64-based TFs
p.wr1E64  = 4 * p.w1E64;
p.wr10E64 = 4 * p.w10E64;

%Other
p.ktl     = 1;
p.kdegR   = 2.7;
p.kdegZFP = 0.35;
p.kdegZFP_PEST = p.kdegZFP * 2;
p.kdegRep = 0.029;
p.ktxCMV  = 1;
p.ktxEF1a = p.ktxCMV;
p.ktxZF   = 1;


end
```
