## Supplementary material for "Model-guided design of mammalian genetic programs": Source Code: README.rtf


1. System requirements

The code can be run using Matlab (https://www.mathworks.com/products/matlab.html). Code was developed and tested on macOS High Sierra.

2. Installation guide

No specific installation is required other than for Matlab.

3. Demo

Files produce an output argument containing the simulated outcomes. The expected run time using the population model is on the order of seconds on a standard desktop computer.

4. Instructions for use

An example run file is provided.
