## Supplementary Material for "Model-guided design of mammalian genetic programs"

**This file includes:**

Materials and Methods
Figs. S1 to S4
Tables S1 to S2
References cited in Supplementary Material

**Other supplementary material includes:**

Source_code.zip
Source_data.xlsx
Plasmid_maps.zip

**Abbreviations**

| | |
|---|---|
| ABA | Abscisic acid |
| ABA-ZFa | Abscisic acid-activated zinc finger activator |
| ABI1 | Abl interactor 1 |
| AD | Activation domain |
| CMV | Cytomegalovirus promoter |
| COMET | COmposable Mammalian Elements of Transcription |
| DMSO | Dimethyl sulfoxide vehicle for rapamycin treatment |
| DsDed | R95K mutant for DsRed-Express2 |
| DsRed | Abbreviation used for DsRed-Express2 |
| ECD | Ectodomain |
| EF1α | Elongation factor-1 alpha promoter |
| EtOH | Ethanol vehicle for rapalog treatment and abscisic acid treatment |
| FKBP | FK506 binding protein |
| FRB | FKBP-rapamycin binding |
| intC | Split intein C-terminal fragment |
| intN | Split intein N-terminal fragment |
| MEFL | Molecules of Equivalent Fluorescein |
| MEPTR | Molecules of Equivalent PE-Texas Red |
| MESA | Modular Extracellular Sensor Architecture |
| MFI | Mean fluorescence intensity |
| MIMO | Multi-input multi-output |
| PC | Protease chain |
| PRS | Protease recognition sequence |
| PYL1 | Pyrabactin resistance 1-like |
| RaZFa | Rapamycin-activated zinc finger activator |
| TC | Target chain |
| TF | Transcription factor |
| TMD | Transmembrane domain |
| VP16 | VP16 activation domain |
| VP64 | VP64 activation domain |
| ZF | Zinc finger |
| ZFa | Zinc finger activator |

### Materials and Methods

#### Experimental method details

##### *Split inteins*

We expanded the COMET toolkit by incorporating gp41-1 (**Fig. 1**, **Fig. S1**): a split intein that was identified putatively in a bioinformatic analysis (*1*), characterized *in vitro* and in *E. coli* (*2*), and later utilized in mammalian cells (*3*). We maintained the TRSGY motif from the native sequence upstream of intN (at the end of the intN-adjoining extein), as done by (*2, 4-6*) to retain high splicing activity; however, gp41-1 splicing has also been reported without this motif (*3*). We note that it is important to use cysteine as the first amino acid of intN ("1" site) and serine as the first amino acid downstream of intC ("+1" site) (*7*).

The protein sequence for intN was:
CLDLKTQVQTPQGMKEISNIQVGDLVLSNTGYNEVLNVFPKSKKKSYKITLEDGKEIICSEEHLFPTQTGE MNISGGLKEGMCLYVKE, where the first amino acid is the "1" site, and TRSGY precedes this site.

The protein sequence for intC was:
MMLKKILKIEELDERELIDIEVSGNHLFYANDILTHNS, where the last amino acid is the "+1" site.

The mutagenesis investigation (**Fig. S4**) was informed by a crystal structure of the gp41-1 C1A catalytically inactive mutant (*8*). Electrostatic interactions between the β3 strand at the end of intN and β6 strand at the start of intC were previously identified to form a charge zipper and proposed to be important in the capture and collapse mechanism that precedes splicing. In this mechanism, which was previously elucidated using the *Npu* DnaE split intein (*9*), capture involves electrostatic interactions between extended regions of the two fragments and collapse involves compaction and stabilization of their initially disordered regions.

##### *Plasmid cloning and purification*

Genetic components that were used in this study are listed in **Table S1**. Plasmids were designed in SnapGene (GSL Biotech LLC), and primers were ordered from Integrated DNA Technologies. Several domains were sourced from Donahue, et al. (*10*), but prior to the COMET study: VP16 and ZF domains are from Khalil, et al. (*11*), VP64 is from Chavez, et al. (Addgene #63798) (*12*), FRB and FKBP are from Daringer, et al. (Addgene #58876, # 58877) (*13*), and DsRed-Express2 is a gift from David Schaffer.

Split inteins are from Hermann, et al. (Addgene #51267, #51268) (*3*). The ABA-binding domains PYL1 and ABI1 (*14-18*) from Gao, et al. (*19*) were utilized to make ABA-ZFa. The PEST tag is from the mouse ornithine decarboxylase gene (*20*). Two types of plasmid backbones are used: pcDNA (pPD005, Addgene #138749), which was modified from Thermo Fisher Scientific #V87020 as described by Donahue, et al. (*10*); and a series of transcription unit positioning vectors (TUPVs), which are derived from the modified pcDNA and previously published by Donahue, et al. (*10*), and based upon the mMoClo system from Duportet, et al. (*21*). Insulator sequences in TUPVs are from Bintu, et al. (Addgene #78099) (*22*).

Cloning was performed primarily using standard PCR, restriction, and ligation methods (reagents from New England Biolabs and Thermo Fisher Scientific), and in some cases through Golden Gate assembly, followed by transformation into chemically competent TOP10 *E. coli* (Thermo Fisher Scientific). Transformed *E. coli* were grown on LB/Ampicillin agar plates at 37°C, colonies were picked and grown in liquid LB/Ampicillin cultures, plasmid DNA was isolated (E.Z.N.A. plasmid mini kit, Omega Bio-tek), and DNA inserts were sequence-verified (ACGT, Inc.). Plasmids were prepared using polyethylene glycol-based extraction as described previously (*10*). DNA purity and concentration were measured using a Nanodrop 2000 (Thermo Fisher Scientific).

##### *Mammalian cell culture*

HEK293FT cells were cultured in complete DMEM medium containing 1% DMEM powder (Gibco #31600091), 0.35% w/v D-glucose (Sigma #50-99-7), 0.37% w/v sodium bicarbonate (Fisher #S233-500), 10% heat-inactivated FBS (Gibco #16140071), 4 mM L-glutamine (Gibco #25030081), and 100 U $ml^{-1}$ penicillin and 100 µg $ml^{-1}$ streptomycin (Gibco #15140122) in tissue culture-treated 10 cm dishes (Corning # 500001672) at 37°C in 5% $CO_2$. To passage, medium was aspirated, and cells were washed in PBS, incubated in trypsin-EDTA (Gibco #25300054; 37°C, 5 min), detached by tapping the dish, and resuspended in fresh medium and plated. This cell line tested negative for *Mycoplasma* using the MycoAlert Mycoplasma detection kit (Lonza #LT07-318).

***Transfection***

Cells were plated in 24-well plates (Corning #3524; $3\times10^5$ cells ml$^{-1}$, 0.5 ml per well) and transfected after adhering to the plates, typically between 8–14 h after plating. Transfections were carried out using the calcium phosphate protocol (*10*): plasmids are mixed together in defined amounts, $CaCl_2$ (2 M, 15% v/v) is added, and this solution is pipetted dropwise into an equal volume of 2x HEPES-buffered saline (500 mM HEPES, 280 mM NaCl, 1.5 mM $Na_2HPO_4$); the solution is gently pipetted four times, and three minutes later it is vigorously pipetted 20 times and added dropwise onto plated cells. In this study, DNA doses are reported in plasmid mass (ng) per well of cells or gene copies per well of cells. In each transfection experiment, "empty vector" (pPD005) was included in the transfection mix to maintain a consistent total mass of DNA per well. At one day after plating, medium was aspirated and replaced with fresh medium. In some experiments, the fresh medium contained vehicle or ligand. In **Fig. 3**, the vehicle was 0.1% DMSO (v/v in cell culture) and the ligand was 100 nM rapamycin in 0.1% DMSO. In **Fig. 4B**, the vehicle was 0.1% EtOH and the ligand was either 100 nM rapalog (Takara #AP21697) or 100 µM abscisic acid (ABA; Goldbio #21293-29-8) in 0.1% EtOH. In **Fig. 4D,E**, the vehicle was 0.2% EtOH, and the ligand conditions included 100 nM rapalog, 100 µM ABA, or both ligands in 0.2% EtOH. For **Fig. S4G**, treatment was applied both prior to transfection and at the time of medium change to promote more immediate inhibitory signaling.

***Flow cytometry***

Samples were prepared for flow cytometry generally at 40–48 h post-transfection. For each well, medium was aspirated, five drops of PBS were added, PBS was aspirated, and two drops of trypsin-EDTA were added. Cells were incubated (37°C, 5 min), plates were tapped to detach cells, and four drops of cold (4°C) DMEM were added. The contents of each well were pipetted up and down several times to detach cells and pipetted into FACS tubes containing FACS buffer (FB; 2 ml; PBS pH 7.4, 5 mM EDTA, 0.1% w/v BSA). Tubes were centrifuged (150×g, 5 min), liquid was decanted, and two drops of FB were added. Samples were kept on ice and wrapped in foil, and then run on a BD LSR Fortessa special order research product using the following configuration: Pacific Blue channel with 405 nm excitation laser and 450/50 nm filter for EBFP2; FITC channel with 488 nm excitation laser and 505LP 530/30 nm filter for EYFP; and PE-Texas Red channel with 552 nm excitation laser and 600LP 610/20 nm filter for mKate2. Approximately $10^4$ live single-cell events were collected per sample.

***Flow cytometry data analysis***

Flow cytometry data were analyzed using FlowJo software (FlowJo, LLC) to gate on single-cell (FSC-A vs. FSC-H) and live (FSC-A vs. SSC-A) bases, compensated using compensation control samples, and gated as transfection-positive (**Fig. S1A**). The mean reporter signal in MFI was obtained for each sample. UltraRainbow Calibration Particles (Spherotech #URCP-100-2H) were run in each flow cytometry experiment. Beads were gated on an FSC-A vs. FSC-H basis, the nine bead subpopulations of varying intensities were identified, and the mean MFI for each subpopulation in the FITC channel and PE-Texas Red channel was obtained. These values in combination with manufacturer-supplied MEFL and MEPTR values for each subpopulation were used to fit a regression line with *y*-intercept equal to zero. The mean and S.E.M. for the three biological replicates were calculated. Autofluorescence background signal was subtracted using samples transfected with the transfection control marker, and error was propagated. MFI values were converted to MEFL or MEPTR using the slope of the regression line, and error was propagated. Histograms in supplementary figure panels represent reporter signal in MFI.

***Nomenclature***

Genes are named by their protein domains in order from N-terminus to C-terminus. Domains are generally connected by flexible linkers comprising glycine and serine. Several abbreviations are used: ZFa is an AD-ZF for any choice of AD and ZF; similarly, RaZFa is an AD-FRB and FKBP-ZF, and ABA-ZFa is an AD-PYL1 and ABI1-ZF. DsRed refers to wild type DsRed-Express2, and DsDed is an DsRed-Express2 R95K mutant. We use a streamlined nomenclature that differs from that used in the original COMET report (*10*), in that inhibitors do not use ZFi notation: ZFi is now termed ZF, and DsRed-ZFi is now termed DsRed-ZF.

The constitutive promoters used are CMV and EF1α. The inducible promoters used are COMET promoters, which are named as "[ZF domain]x[number of binding sites]-[binding site arrangement]". For example, ZF1x6-C has six compact sites for ZF1. There are two non-standard cases: ZF1/2x6-C has six compact overlapping sites for ZF1 and ZF2 (up to six sites occupied, and up to six per ZF); (ZF2/ZF6)x3 has six compact sites alternating between ZF2 and ZF6 (up to six sites occupied, and up to three per ZF).

***Statistical analysis***
Each sensor in **Fig. 4A,B** was assessed using a one-tailed Welch's unpaired *t*-test, with the null hypothesis that reporter signal was equal with and without ligand treatment. Genetic programs in **Fig. 4D,E** were assessed using a three-factor ANOVA and Tukey's honest significant difference (HSD) test, with the null hypothesis that reporter signal was equal across the two input types, four topologies, and four input combinations. Effects were considered significant if $p < 0.05$, and additionally for the HSD test if the comparisons had an adjusted $p < 0.05$.

### Computational method details

***Overview***
This section describes the extension of our original explanatory model for COMET TFs (*10*) to a predictive model incorporating split intein-mediated splicing and other attributes. Rules for formulating systems of ODEs are provided to support the formulation of models for new genetic circuits based on the genetic parts from this study.

***A statistical model for gene expression heterogeneity***
Our modeling approach accounts for variation in gene expression—including differences in the expression of a gene between cells and differences in the expression of genes within a cell—in a cell population. We generate a population matrix using the constrained sampling method (*10, 23*), which is used here to describe the distribution of gene expression observed when cells are harvested via trypsin digest (as noted in **Fig. S1G**, this distribution differs somewhat from the distribution observed when cells are harvested by another common method, suspension in FACS buffer).

The *i*th row (cell) and *p*th column (plasmid) of the population matrix **Z** is a scalar for the relative expression of a gene. The $z$ value is used as a multiplier in the production term for each RNA species.

$$\text{Production rate} = z_{i,p} \cdot k_{\text{txEF1a}} \cdot \text{dose} \tag{1}$$

A dynamical model is run separately for each cell in the simulated population, and the mean end-point simulated reporter protein level for the population is calculated. Layering the statistical model on the dynamical models at the level of RNA production enables the simulations to account for cell-to-cell variation (e.g., this method incorporates potential outlier effects that could skew a population mean) and therefore should generally enable better predictions (*23*).

Some supplementary figures employ a standard single-cell (homogeneous) model, as indicated in the figure captions. These cases forgo the incorporation of heterogeneity and instead simulate the mean-transfected cell, which represents a scenario for average gene expression from each plasmid.

***Dynamical models***
Genetic programs are represented by systems of ODEs, which are included in Source_code.zip. State variables include RNA and protein species in arbitrary concentration units. Processes include transcription (constitutive, inducible, inhibitable), RNA degradation, protein translation, split intein-mediated splicing, small molecule-based reconstitution, and protein degradation. Parameter values are in **Table S2**.

Constitutive transcription from the EF1α promoter or CMV promoter is proportional to plasmid dose (ng).

$$\text{RNA production rate} = k_{\text{txEF1a}} \cdot \text{dose} \tag{2}$$

$$\text{RNA production rate} \ = \ k_{\text{txCMV}} \cdot \text{dose} \tag{3}$$

Functions for regulated transcription are broadly represented by $f$. The dose term $\mathrm{d}$ for a regulated gene is empirically defined (i.e., based on a heuristic) and calculated by dividing the plasmid dose (ng) by 200 ng; then, the square root of this fraction is used, e.g., for 200 ng, $\mathrm{d}^{1/2} = 1$, and for 50 ng, $\mathrm{d}^{1/2} = 0.5$. The 200 ng dose was defined as a reference point because this was the dose of reporter plasmid used in the original characterization (*10*).

$$\text{RNA production rate} \ = \ k_{\text{txZF}} \cdot \mathrm{d}^{1/2} \cdot f \tag{4}$$

ZFa-inducible transcription uses the COMET model formulation, in which $b$ is TF-independent (background) transcription, $m$ is the maximal activation, and $w$ is a steepness parameter. We model the activation mediated by AD-ZF-containing proteins that also contain intC, intN, or additional ZF domains equivalently to that by a base case ZFa. The variable refers to the simulated amount of TF protein, not to plasmid dose.

$$f = \frac{b + m \cdot w \cdot \mathbf{ZFa_{Protein}}}{1 + w \cdot \mathbf{ZFa_{Protein}}} \tag{5}$$

ZFa-inducible transcription can be inhibited by a ZF, which sterically blocks the activator from binding to sites in a promoter. We model the inhibition mediated by ZF proteins that also contain intC, intN, FKBP, or additional ZF domains equivalently to that by a base case ZF. The subscripts $\mathrm{A}$ and $\mathrm{I}$ denote parameters for an activator and inhibitor, respectively.

$$f = \frac{b + m_{\mathrm{A}} \cdot w_{\mathrm{A}} \cdot \mathbf{ZFa_{Protein}}}{1 + w_{\mathrm{A}} \cdot \mathbf{ZFa_{Protein}} + w_{\mathrm{I}} \cdot \mathbf{ZF_{Protein}}} \tag{6}$$

ZFa-inducible transcription can also be inhibited by a DsDed-ZF, which acts through a dual mechanism of steric inhibition and reduction of effective promoter cooperativity. The effect of the latter mechanism is that at increasing strength or dose of inhibitor compared to activator, the cooperativity represented by $m$ ramps down to an effective value of 1. We model the inhibition mediated by DsDed-ZF-containing proteins that also contain intC, intN, or additional ZF domains equivalently to that by a base case DsDed-ZF.

$$f = \frac{b + \max\left(\min\left(\frac{\left(\frac{4 \cdot w_{\mathrm{I}} \cdot \mathbf{DsDed\text{-}ZF_{Protein}}}{w_{\mathrm{A}} \cdot \mathbf{ZFa_{Protein}}} - 4 \cdot l\right)(1-m)}{4 \cdot u - 4 \cdot l} + m, m\right), 1\right) \cdot w_{\mathrm{A}} \cdot \mathbf{ZFa_{Protein}}}{1 + w_{\mathrm{A}} \cdot \mathbf{ZFa_{Protein}} + 4 \cdot w_{\mathrm{I}} \cdot \mathbf{DsDed\text{-}ZF_{Protein}}} \tag{7}$$

RNA degradation is represented as a first-order process.

$$\text{RNA degradation rate} \ = \ -k_{\text{degRNA}} \cdot \mathbf{Species_{RNA}} \tag{8}$$

Protein translation is also first order.

$$\text{Translation rate} \ = \ k_{\text{tl}} \cdot \mathbf{Species_{RNA}} \tag{9}$$

Splicing is a second-order reaction between an intN-containing protein and an intC-containing protein with a fitted rate constant.

$$\text{Splicing rate} \ = \ k_{\text{rec}} \cdot \mathbf{Species1_{Protein}} \cdot \mathbf{Species2_{Protein}} \tag{10}$$

For example, the following the terms represent the splicing of $\mathrm{A\text{-}intN\text{-}B}$ and $\mathrm{X\text{-}intC\text{-}Y}$ to $\mathrm{A\text{-}Y}$ and $\mathrm{X\text{-}intC/intN\text{-}B}$, where $\mathrm{A}$, $\mathrm{B}$, $\mathrm{X}$, and $\mathrm{Y}$ can be DNA-binding, activating, or inhibitory domains or no domain.

$$\textbf{A-intN-B}_{\textbf{Protein}} \text{ splicing rate } = -k_{\text{rec}} \cdot \textbf{A-intN-B}_{\textbf{Protein}} \cdot \textbf{X-intC-Y}_{\textbf{Protein}} \tag{11}$$

$$\textbf{X-intC-Y}_{\textbf{Protein}} \text{ splicing rate } = -k_{\text{rec}} \cdot \textbf{A-intN-B}_{\textbf{Protein}} \cdot \textbf{X-intC-Y}_{\textbf{Protein}} \tag{12}$$

$$\textbf{A-Y}_{\textbf{Protein}} \text{ splicing rate } = k_{\text{rec}} \cdot \textbf{A-intN-B}_{\textbf{Protein}} \cdot \textbf{X-intC-Y}_{\textbf{Protein}} \tag{13}$$

$$\textbf{X-intC/intN-B}_{\textbf{Protein}} \text{ splicing rate } = k_{\text{rec}} \cdot \textbf{A-intN-B}_{\textbf{Protein}} \cdot \textbf{X-intC-Y}_{\textbf{Protein}} \tag{14}$$

Small molecule-based reconstitution to form RaZFa uses the Heaviside function *H* with ligand treatment at time $\tau$ (hours) post-transfection. For simplicity, the $k_{\text{rec}}$ parameter is also used to describe reconstitution.

$$\text{RaZFa reconstitution rate } = k_{\text{rec}} \cdot \textbf{AD-FRB}_{\textbf{Protein}} \cdot \textbf{FKBP-ZF}_{\textbf{Protein}} \cdot H(t-\tau) \tag{15}$$

Prior to reconstitution, FKBP-ZF can act as a ZF-like inhibitor against RaZFa or ZFa at a target promoter.

$$\text{RNA production rate} = k_{\text{txZF}} \cdot \frac{b + m \cdot w \cdot \textbf{RaZFa}_{\textbf{Protein}}}{1 + w \cdot \textbf{RaZFa}_{\textbf{Protein}} + w \cdot \textbf{FKBP-ZF}_{\textbf{Protein}}} \tag{16}$$

Protein degradation is first order. Rate constants vary for non-intC-containing TFs, intC-containing TFs, and reporter protein, respectively.

$$\text{Protein degradation rate } = -k_{\text{degZFP}} \cdot \textbf{TF}_{\textbf{Protein}} \tag{17}$$

$$\text{Protein degradation rate } = -k_{\text{degintC}} \cdot \textbf{TF}_{\textbf{Protein}} \tag{18}$$

$$\text{Protein degradation rate } = -k_{\text{degRep}} \cdot \textbf{Reporter}_{\textbf{Protein}} \tag{19}$$

As an example, the following system of equations represents the reconstitution of a ZFa and induction of a reporter. This system produces an AND gate for: if AD-intN and intC-ZF are present, then induce reporter.

$$\frac{d\textbf{AD-intN}_{\textbf{RNA}}}{dt} = \begin{aligned}&z_{i,1} \cdot k_{\text{txEF1a}} \cdot \text{dose}_{\text{AD-intN}}\\ &-k_{\text{degRNA}} \cdot \textbf{AD-intN}_{\textbf{RNA}}\end{aligned} \tag{20}$$

$$\frac{d\textbf{AD-intN}_{\textbf{Protein}}}{dt} = \begin{aligned}&k_{\text{tl}} \cdot \textbf{AD-intN}_{\textbf{RNA}}\\ &-k_{\text{rec}} \cdot \textbf{AD-intN}_{\textbf{Protein}} \cdot \textbf{intC-ZF}_{\textbf{Protein}}\\ &-k_{\text{degZFP}} \cdot \textbf{AD-intN}_{\textbf{Protein}}\end{aligned} \tag{21}$$

$$\frac{d\textbf{intC-ZF}_{\textbf{RNA}}}{dt} = \begin{aligned}&z_{i,2} \cdot k_{\text{txEF1a}} \cdot \text{dose}_{\text{intC-ZF}}\\ &-k_{\text{degRNA}} \cdot \textbf{intC-ZF}_{\textbf{RNA}}\end{aligned} \tag{22}$$

$$\frac{d\textbf{intC-ZF}_{\textbf{Protein}}}{dt} = \begin{aligned}&k_{\text{tl}} \cdot \textbf{intC-ZF}_{\textbf{RNA}}\\ &-k_{\text{rec}} \cdot \textbf{AD-intN}_{\textbf{Protein}} \cdot \textbf{intC-ZF}_{\textbf{Protein}}\\ &-k_{\text{degintC}} \cdot \textbf{intC-ZF}_{\textbf{Protein}}\end{aligned} \tag{23}$$

$$\frac{d\textbf{AD-ZF}_{\textbf{Protein}}}{dt} = \begin{aligned}&k_{\text{rec}} \cdot \textbf{AD-intN}_{\textbf{Protein}} \cdot \textbf{intC-ZF}_{\textbf{Protein}}\\ &-k_{\text{degZFP}} \cdot \textbf{AD-ZF}_{\textbf{Protein}}\end{aligned} \tag{24}$$

$$\frac{d\textbf{intC/intN}_{\textbf{Protein}}}{dt} = \begin{aligned}&k_{\text{rec}} \cdot \textbf{AD-intN}_{\textbf{Protein}} \cdot \textbf{intC-ZF}_{\textbf{Protein}}\\ &-k_{\text{degintC}} \cdot \textbf{intC/intN}_{\textbf{Protein}}\end{aligned} \tag{25}$$

$$\frac{d\mathbf{Reporter_{RNA}}}{dt} = z_{i,3} \cdot k_{\mathrm{txZF}} \cdot \mathrm{d}_{\mathrm{Reporter}}^{1/2} \cdot \frac{b + m \cdot w \cdot \mathbf{AD\text{-}ZF_{Protein}}}{1 + w \cdot \mathbf{AD\text{-}ZF_{Protein}} + w \cdot \mathbf{intC\text{-}ZF_{Protein}}} - k_{\mathrm{degRNA}} \cdot \mathbf{Reporter_{RNA}} \tag{26}$$

$$\frac{d\mathbf{Peporter_{Protein}}}{dt} = k_{\mathrm{tl}} \cdot \mathbf{Reporter_{RNA}} - k_{\mathrm{degRep}} \cdot \mathbf{Reporter_{Protein}} \tag{27}$$

Since the genetic parts employed for activation, inhibition, splicing, and dimerization exhibit functional modularity, one can utilize the formalisms described above to generate systems of equations to represent a variety of circuits. ODEs for the circuits in this study are provided in MATLAB files in Source_code.zip.

***Parameterization***
Some parameter values are from the COMET study (*10*) and others are newly estimated or fitted here (**Table S2**).

***Ultrasensitivity***
Ultrasensitivity is a type of nonlinear signal processing in which a small change in an input produces a large change in an output. We demonstrate how this property can be achieved with engineered motifs such as a double inhibition cascade (**Fig. 1K**), activation thresholded by an inhibitor (**Fig. 3B**), and reconstitutable activation (**Fig. 3C**). The ultrasensitivity of experimental and simulated dose responses is quantified using the Hill coefficient $n$ from a modified Hill equation, in which $x$ is input plasmid dose (ng), $y$ is reporter signal (MEPTRs or MEFLs), $y_0$ is reporter signal for zero input, and $a$ and $b$ are other fitted parameters. Standard ZFa dose responses are characterized by $n$ ~ 1. Ultrasensitive responses are those for which $n$ > 1.

$$y = y_0 + \frac{a \cdot x^n}{\left(\frac{1}{b}\right)^n + x^n} \tag{28}$$

***Diagrams***
Genetic programs for digital functions are depicted using genetic diagrams and electronic diagrams. The former represents each promoter, protein, and regulatory interaction, and the latter represents the logic underlying these interactions.

### A Flow cytometry gating

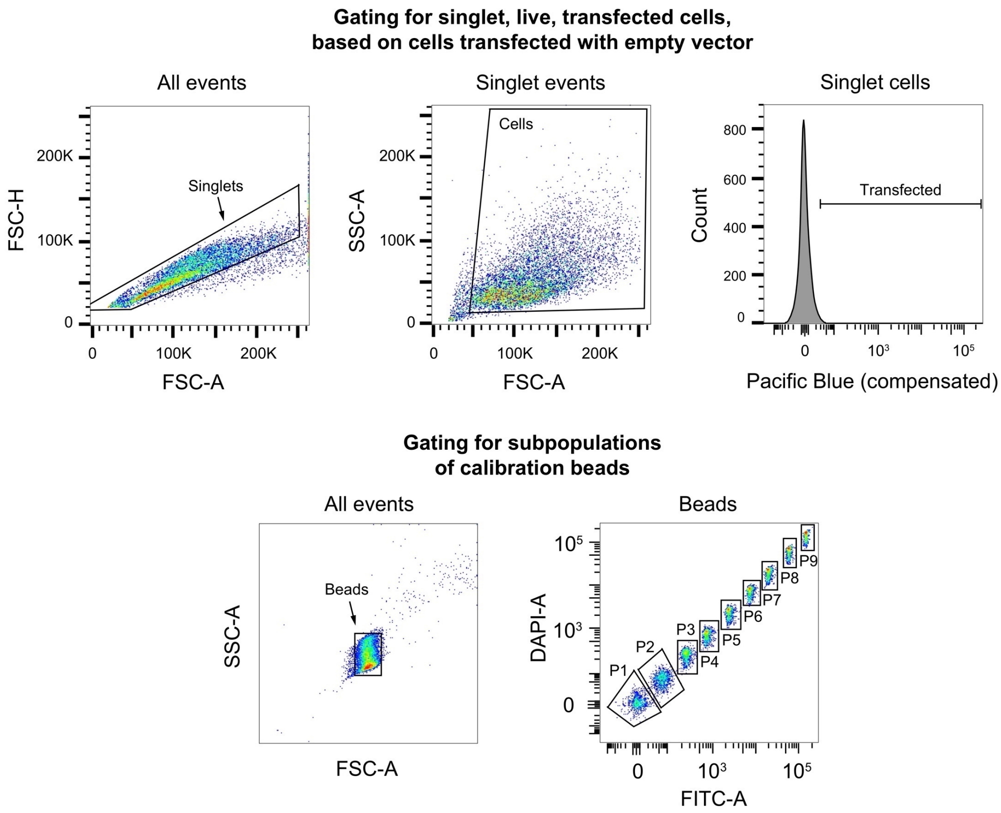

**Fig. S1. Development of the genetic components and models.** (**A**) Flow cytometry gating was used to identify live cells (FSC-A vs. SSC-A), singlet events (FSC-A vs. FSC-H), and EBFP2 (transfection control) expression (Pacific Blue channel). The EBFP2+ gate was defined by setting a threshold to include the top ~1% of live single-cell events for cells transfected with empty vector only.

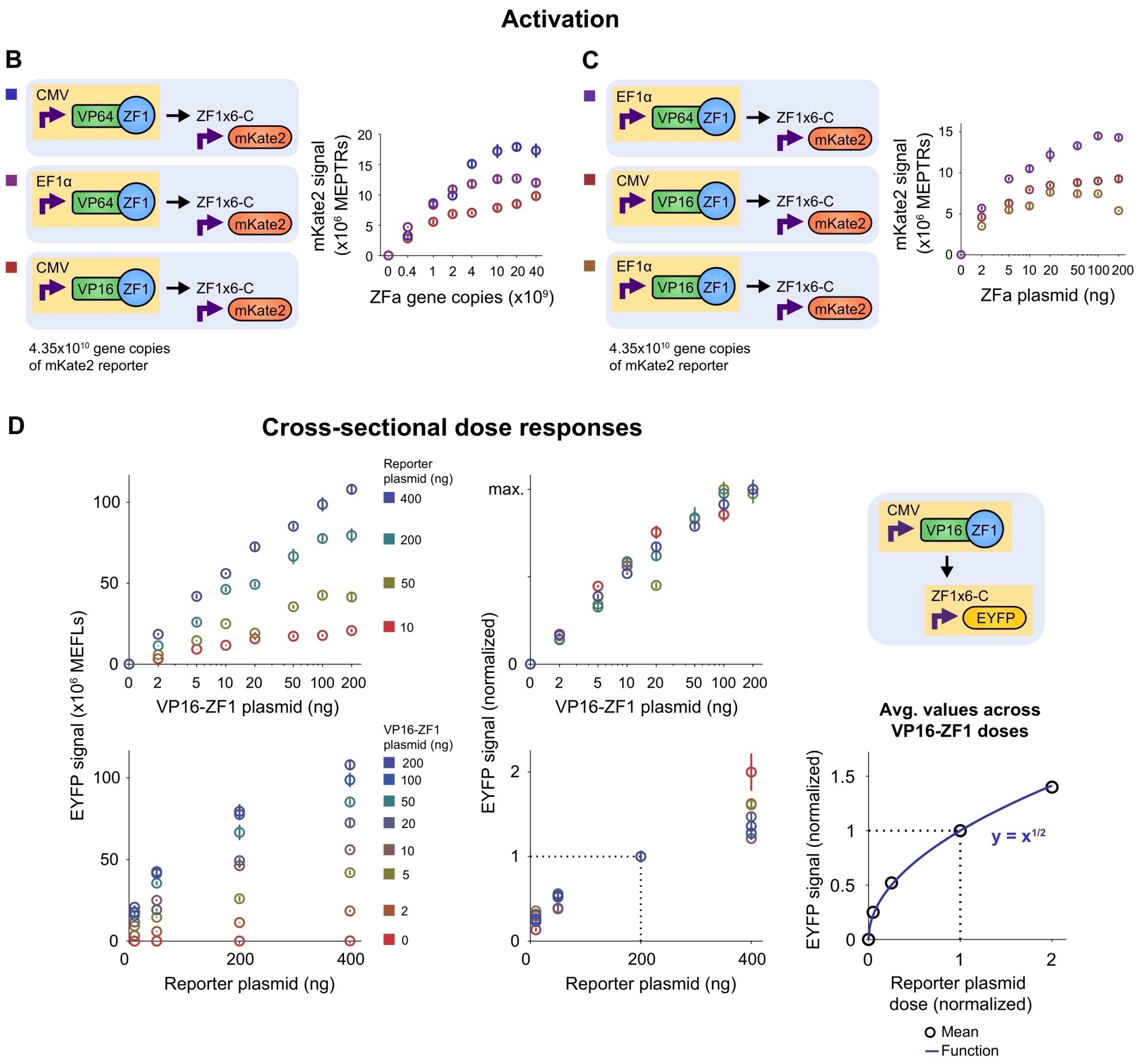

**Fig. S1** (**B–C**) Trends in mKate2 reporter signal for ZF1-based activator dose responses vary depending on choice of constitutive promoter (CMV, EF1α) and activation domain (VP16, VP64); the combinations are color-coded. These are the types of dose responses from which transcriptional activation parameters can be estimated (**Materials and Methods**). Component doses indicate the amount transfected per well in units of either gene copies or plasmid mass. Diagrams use these conventions: purple right-angle arrows are promoters; black arrows are transitions (splicing, ligand binding, proteolytic cleavage) or transcriptional regulation (activation, inhibition); highlighting is for components for which dose is varied. In these diagrams, RNA species and the transfection control marker EBFP2 are not depicted. (**D**) The plots display cross-sections of the data for EYFP reporter signal across VP16-ZF1 activator plasmid doses and EYFP reporter plasmid doses, which are color-coded. Reporter signal follows an approximately square root relationship with reporter plasmid dose, and this trend holds across activator plasmid doses; this observation was used to arrive at the heuristic in Eqn. 4.

**E** **Split intein-mediated reconstitution of activation**

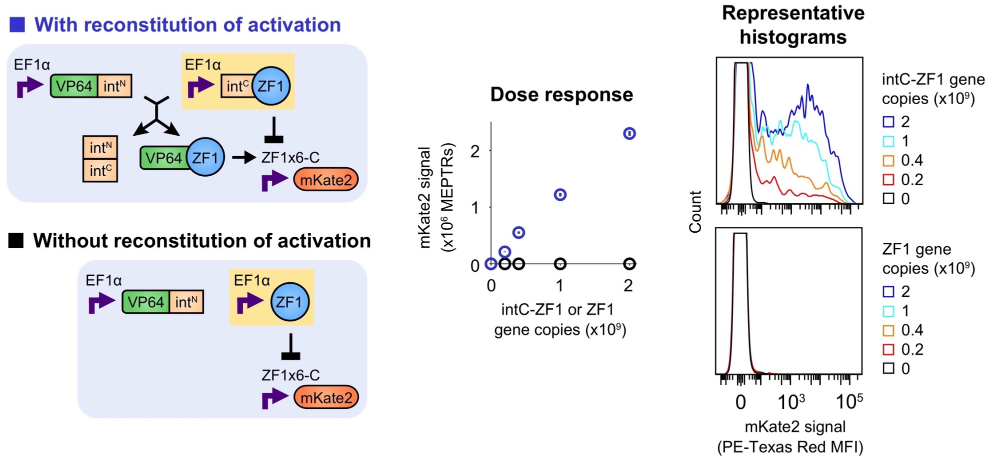

**F** **Fitted parameters for split intein-mediated reconstitution and intC-ZF1 degradation**

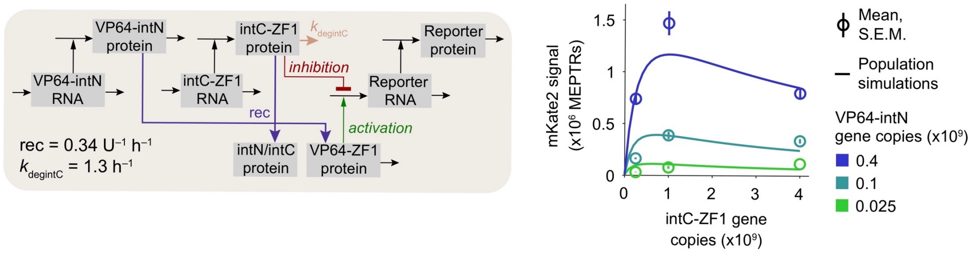

**Fig. S1** (**E**) COMET TF function can be reconstituted using gp41-1 split inteins. The split inteins comprise an N-terminal fragment (intN) and a C-terminal fragment (intC) that can be used to ligate proteins. The splicing reaction proceeds as A-intN-B + X-intC-Y → A-Y + X-intC/intN-B, where A, B, X, and Y denote adjoining protein sequences (exteins) or a lack thereof, "-" denotes a peptide bond, and "/" denotes a non-covalent interaction. Here, activation is reconstituted by ligating ZF1 and VP64. The results indicate that in the absence of intC on ZF1, co-expression of VP64-intN and ZF1 does not induce reporter expression. For the histograms, y-axes are scaled to distinguish each condition, such that the near-zero-signal region of each histogram is truncated. Color-coding indicates intC-ZF1 and ZF1 gene copies. (**F**) Model parameters for split intein-mediated reconstitution (rec) and degradation of intC-containing proteins ($k_{\mathrm{degintC}}$) were fit using the AND gate data in **Fig. 1C**. The diagram depicts the state variables including RNAs and proteins. Color-coding indicates VP64-intN gene copies.

### G Distribution of gene expression in a cell population

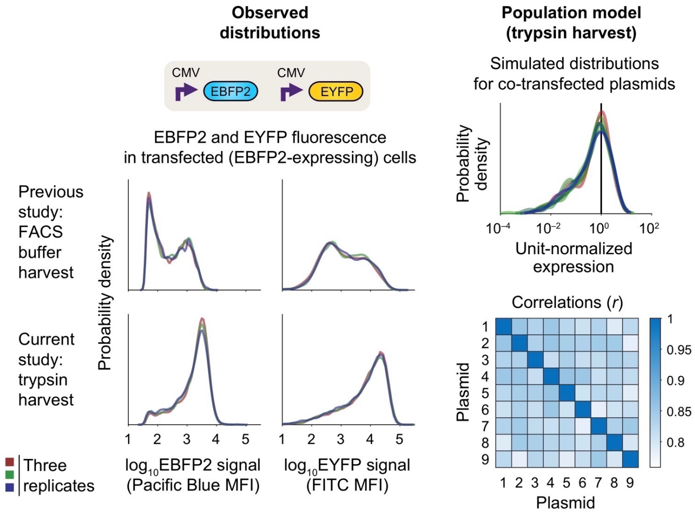

**Fig. S1** (**G**) A statistical model for observed variation in gene expression was used to simulate heterogeneous cell populations (**Materials and Methods**). This approach was first developed in (*23*) and later used in (*10*). The flow cytometric distributions (collected in the same experiment) are EBFP2 signal and EYFP signal after gating on EBFP2-expressing (transfected) cells. The current study uses a statistical model based on the distribution observed when cells are harvested for flow cytometry using a trypsin digest; this model differs somewhat from that which we previously utilized in a study in which cells were harvested by a slightly different but common method (FACS buffer harvest) (*10*). A key lesson is that the method used to prepare cells for flow cytometry affects the observed fluorescence distribution, and thus the mathematical description should be matched to the choice of experimental method. We expect that an experiment like the one in this figure could be employed to generate a statistical distribution for any given method of gene delivery and cell harvest as needed. The right panels are unit-normalized simulated distributions of gene expression and simulated correlations for co-expressed genes on transfected plasmids, all based on the generated population **Z** matrix. This matrix is used to introduce heterogeneity into dynamical models by applying unitless multipliers to the rates of transcription from each plasmid (columns) from each cell (rows). The algorithm for generating **Z** as described in (*10*) ensures that relative expression from each plasmid species is similarly distributed across cells (upper right) and that expression across plasmid species is similarly correlated ($r \sim 0.8$) within cells (lower right).

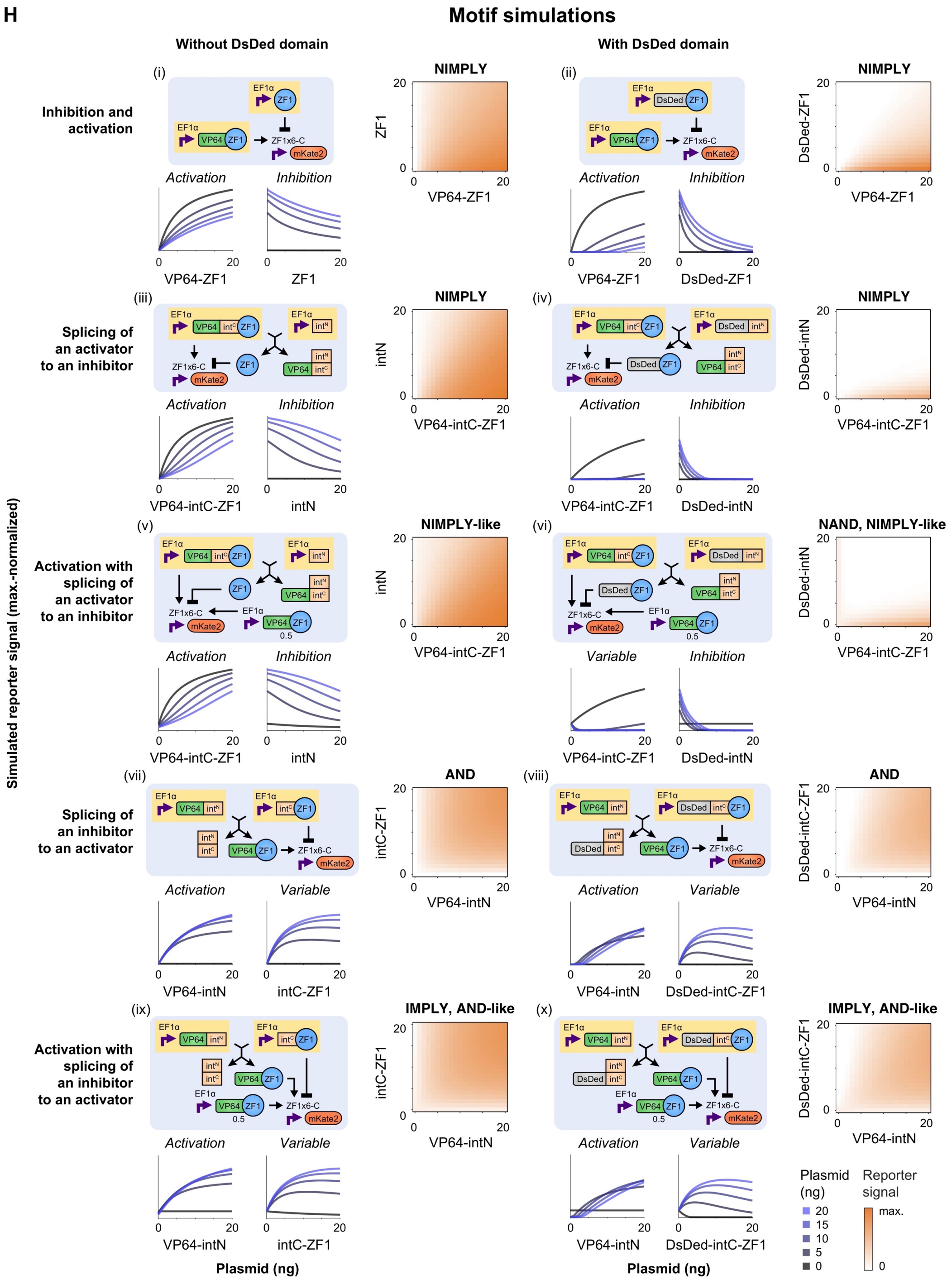

**Fig. S1** (**H**) The incorporation of split inteins provides new ways to utilize TFs. Homogeneous (single-cell) simulations were conducted for a panel of proposed motifs to survey the responses that could be generated beyond COMET base cases (i, ii). Plots below each diagram display cross-sections of plasmid dose responses: one component is varied along the x-axis, and the other component is varied as specified by the blue color-coding. The descriptor above each plot indicates whether the dose response outcome is activating or inhibitory, or if these outcomes vary depending on the dose of the other component. The heatmaps provide more continuous depictions of outcomes (red color-coding scaled to the maximum value across the ten cases), and the descriptor above each heatmap indicates the type of gate that each outcome resembles. The new motifs (iii–x) broaden the types of responses that can be generated.

**I** **Characterization of a non-fluorescent bulky inhibitor**

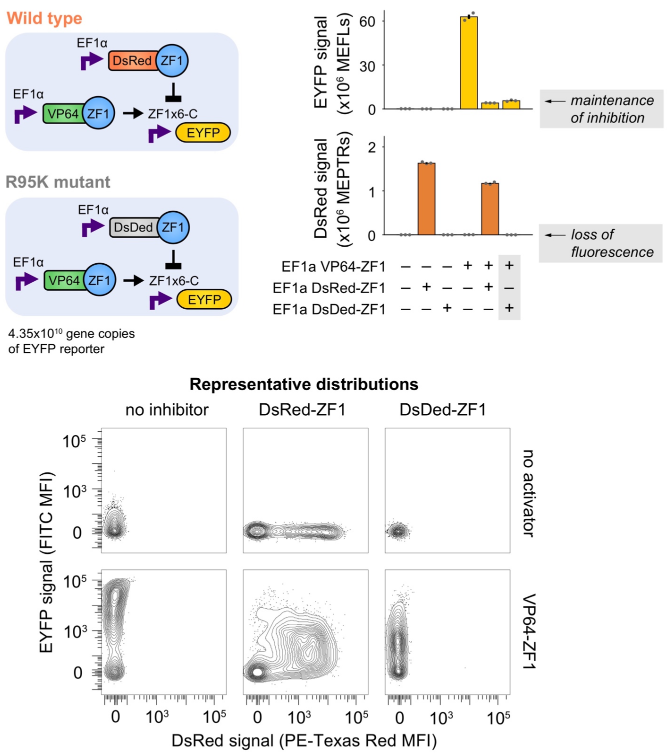

**Fig. S1** (**I**) The emission spectrum of mKate2 overlaps with that of DsRed-Express2 (abbreviated as DsRed), so we sought to remove this fluorescent signal from the DsRed-ZF inhibitor. An R95K mutation (*24*) was introduced to the catalytic triad of the chromophore to generate a variant that we termed DsDed-ZF. This TF produced no detectable DsRed signal, and it retained inhibitory potency against a VP64-ZF1 activator as evident from the low EYFP reporter signal.

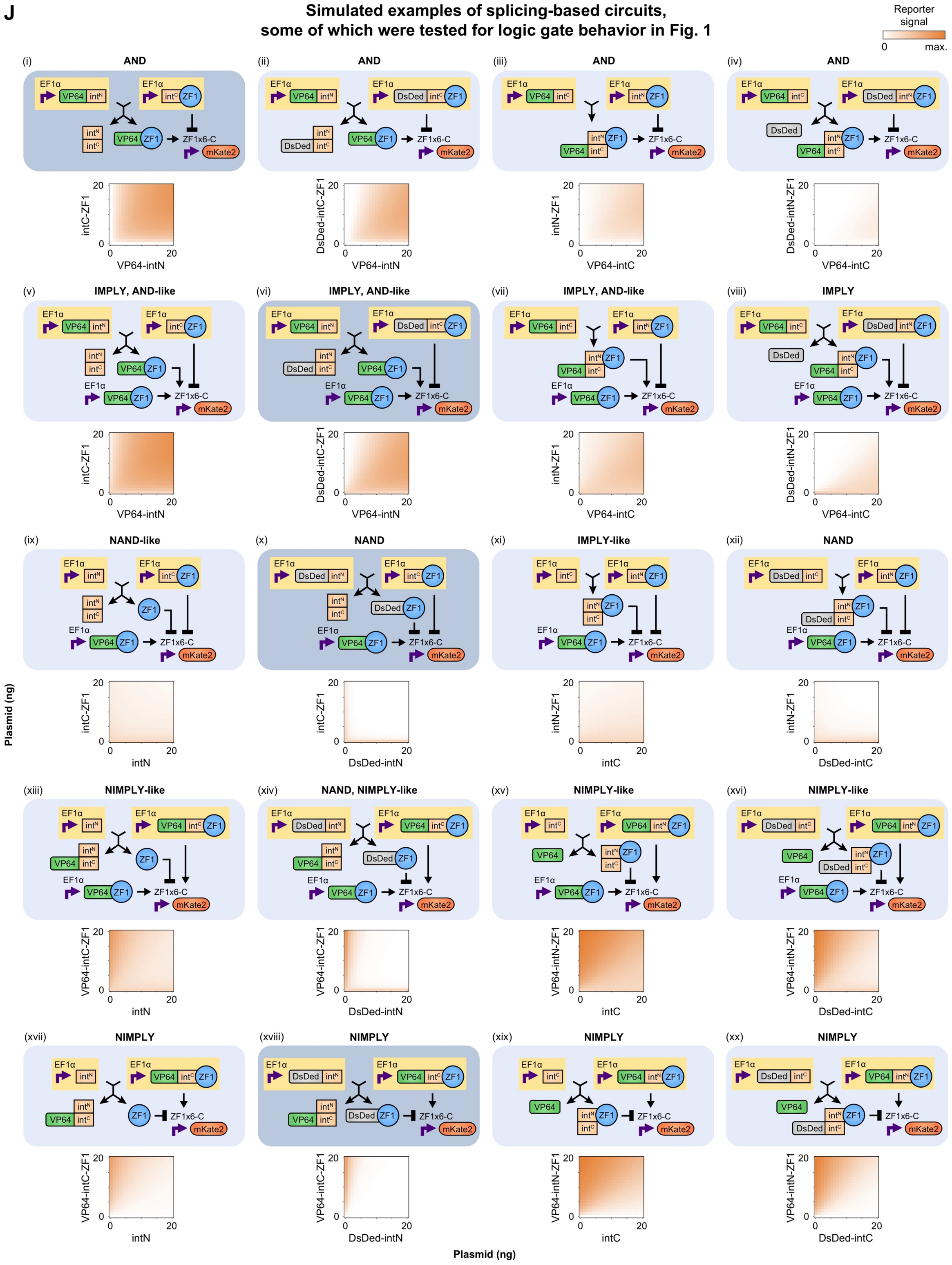

**Fig. S1** (**J**) A panel of splicing-based circuits was proposed, and population simulations were conducted to identify designs for experimental implementation and testing. The descriptor above each diagram indicates the gate(s) that each outcome matches and/or resembles (where “-like” indicates imperfect performance for the stated gate outcome). In (v–xvi), the plasmid dose of constitutive VP64-ZF1 was set to 1 ng. Circuits diagrammed with a dark background (i, vi, x, xviii) were among those tested in **Fig. 1**.

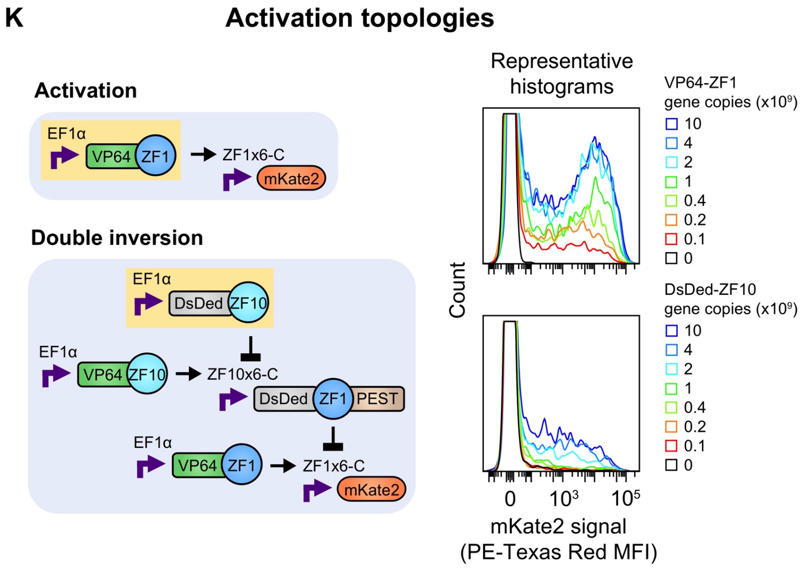

**Fig. S1** (**K**) Comparison of reporter signal in standard activation and in a double inversion cascade. The cascade produces an ultrasensitive response; *n* is the exponent from the Hill equation (Eqn. 28) fitted to data. For the histograms, *y*-axes are scaled to distinguish each condition, such that the region near zero signal of each histogram is truncated. Color-coding indicates component doses.

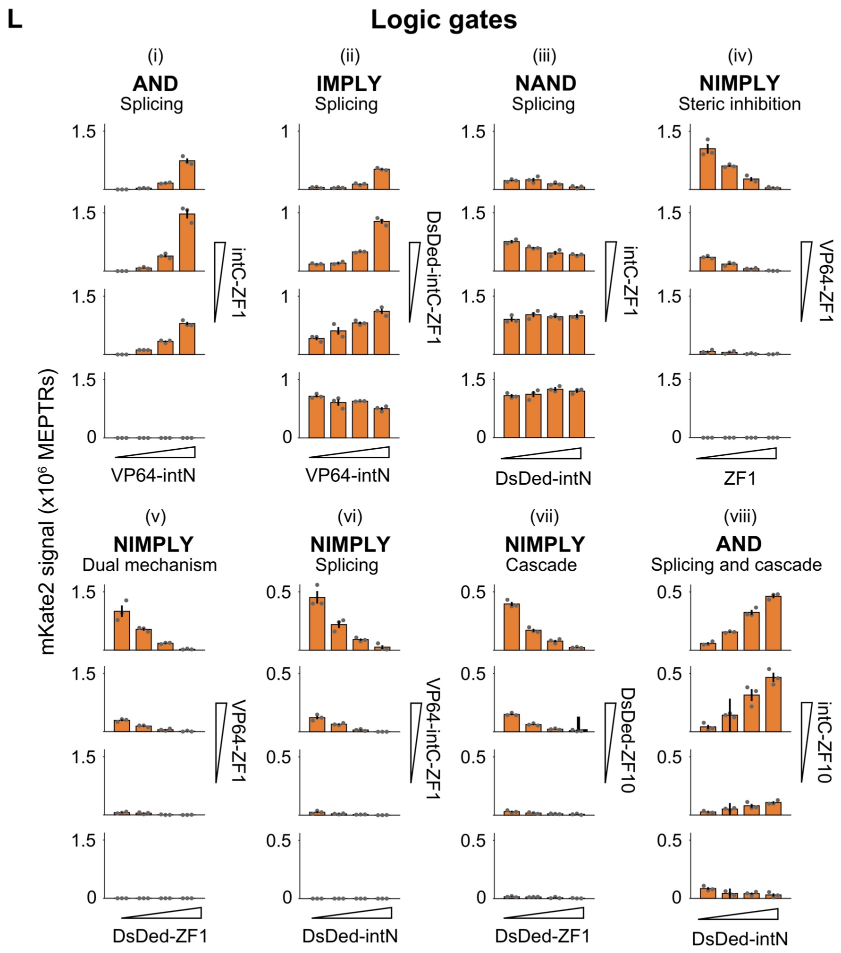

**Fig. S1** (**L**) Bar graphs represent the mean and S.E.M. of reporter signal from three biological replicates. Data correspond to **Fig. 1C–J**. Data for (i, ii, vii), (iii, iv, v, viii), and (vi) were collected separately.

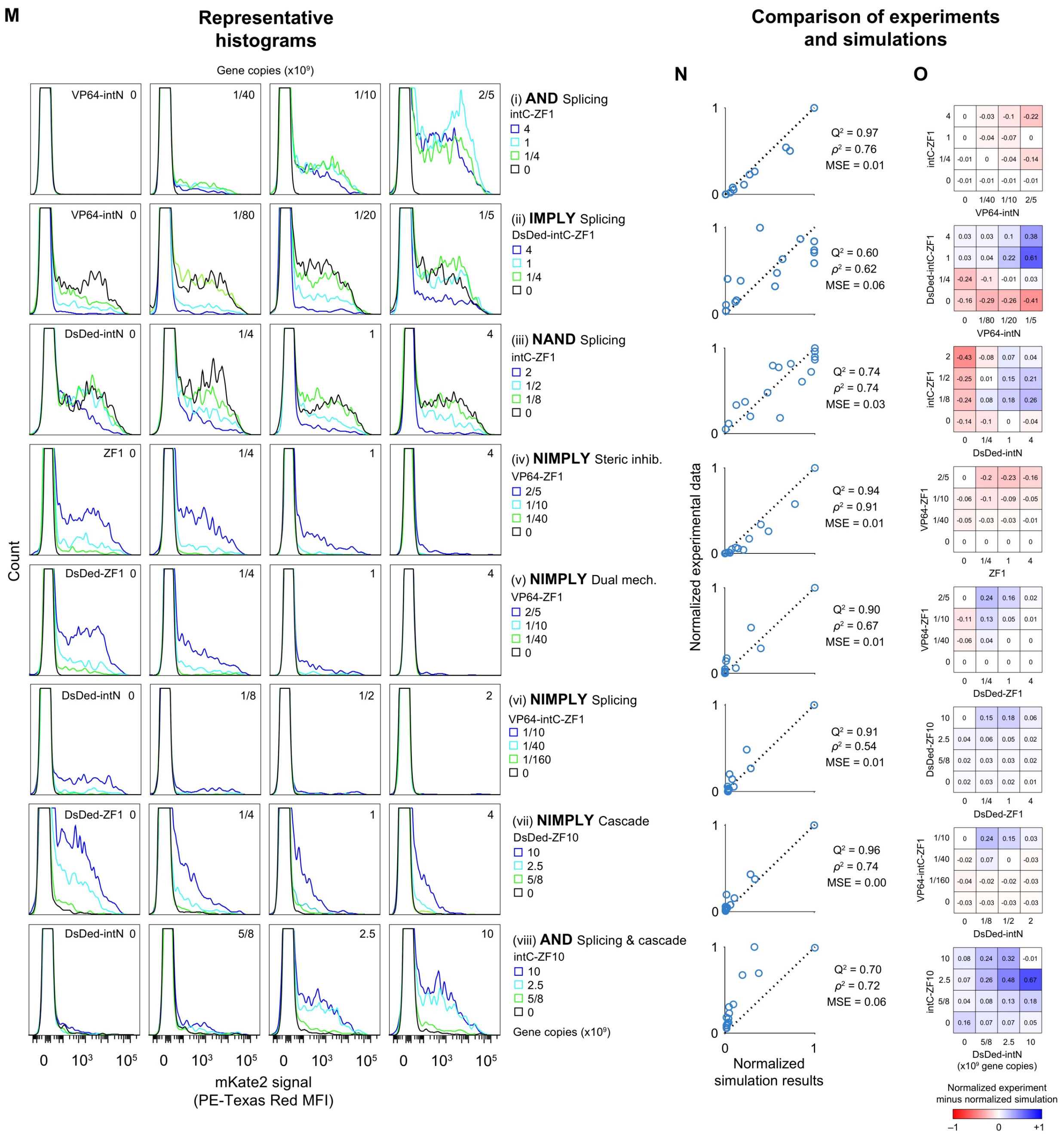

**Fig. S1** Data and simulations correspond to **Fig. 1C–J**: (**M**) Representative flow cytometry histograms; (**N**) goodness of prediction ($Q^2$), Spearman's rank correlation coefficient squared ($\rho^2$), and mean squared error (MSE) comparing max.-normalized simulated and observed mean signal. (**O**) Difference heatmaps for max.-normalized simulated and observed mean signal. The $Q^2$ values indicate that for all eight gates, the simulations explain the majority of the variance in the data; five gates provide very close agreement (>90% $Q^2$).

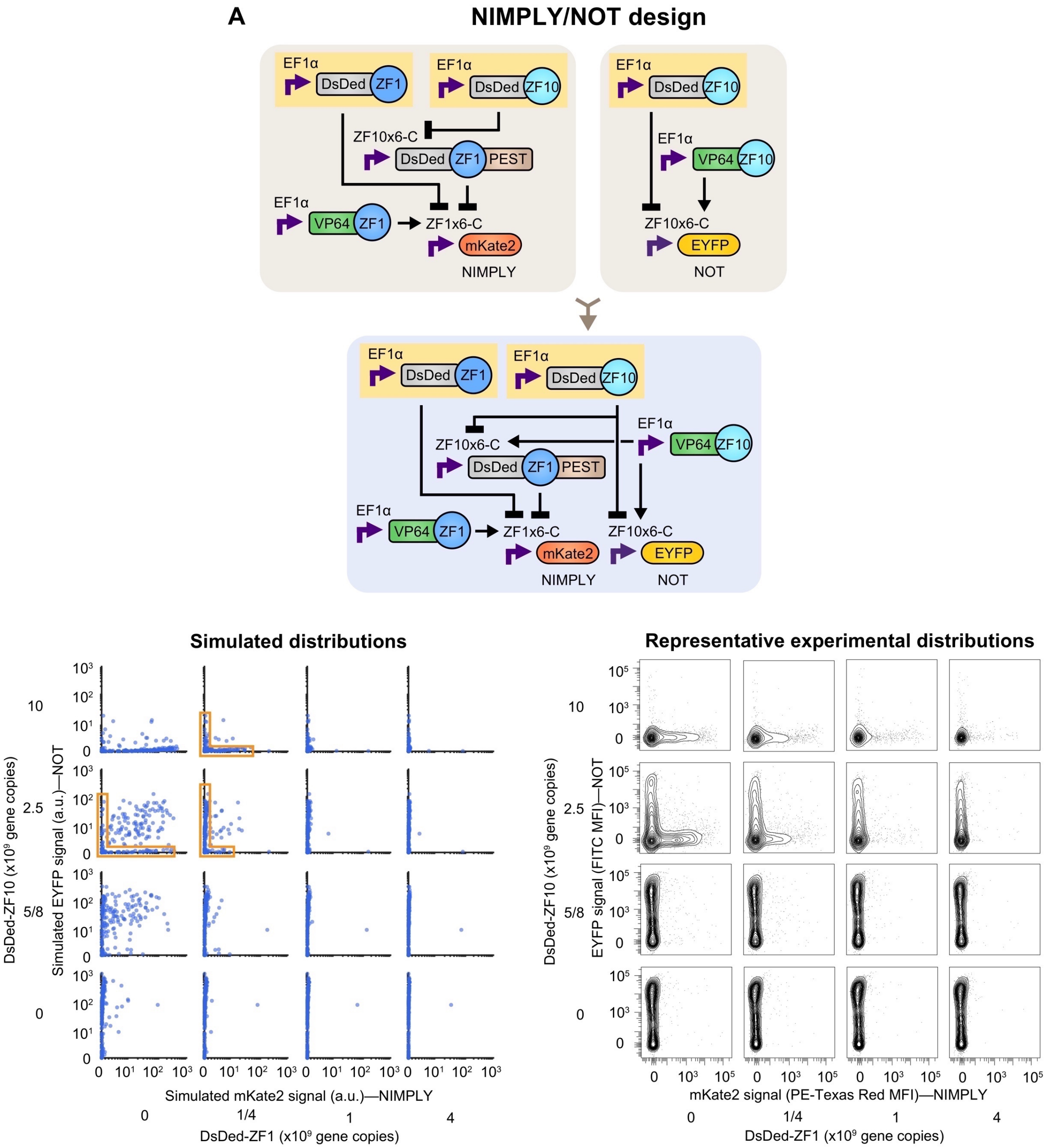

**Fig. S2. MIMO gate designs and single-cell outcomes.** (**A–G**) Data correspond to **Fig. 2B–F** and were collected in the same experiment. (**A–E**) compression of genetic programs to arrive at MIMO designs, population simulations (200 cells; signals in model-specific a.u.), and corresponding representative flow cytometric distributions (in MFI) for mKate2 and EYFP reporter signals. Orange outlines for three of the gates (**Fig. S2A,D,E**) denote cases in which some cells are predicted to substantially express at most one reporter or the other but not both. This task distribution is also evident in the experimental data. This result highlights how the approach used here for representing cell heterogeneity can capture different types of population outcomes. All of the distribution plots use logicle axis scaling. (**A**) NIMPLY/AND corresponding to **Fig. 2B**.

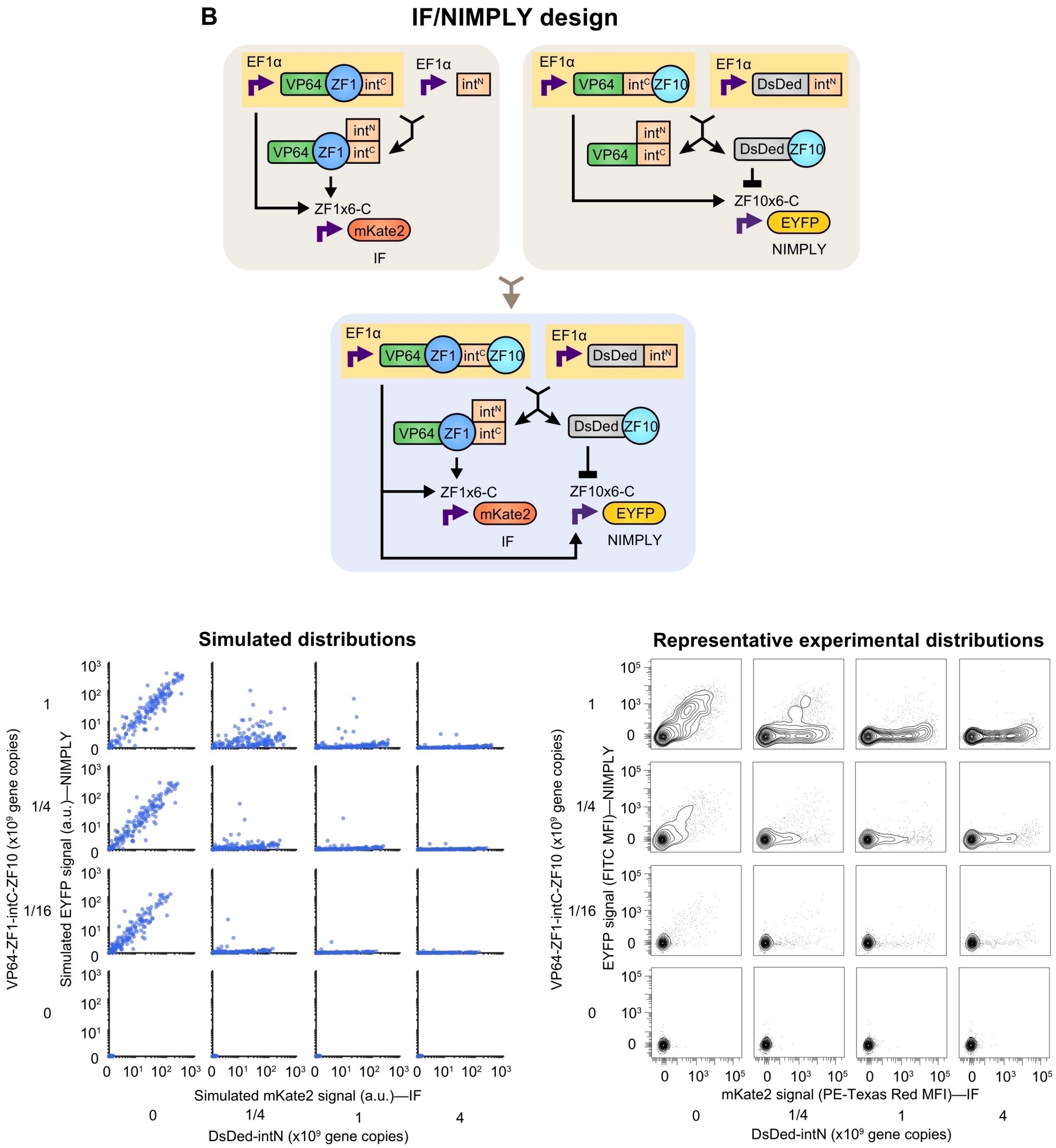

**Fig. S2** (**B**) IF/NIMPLY corresponding to **Fig. 2C**.

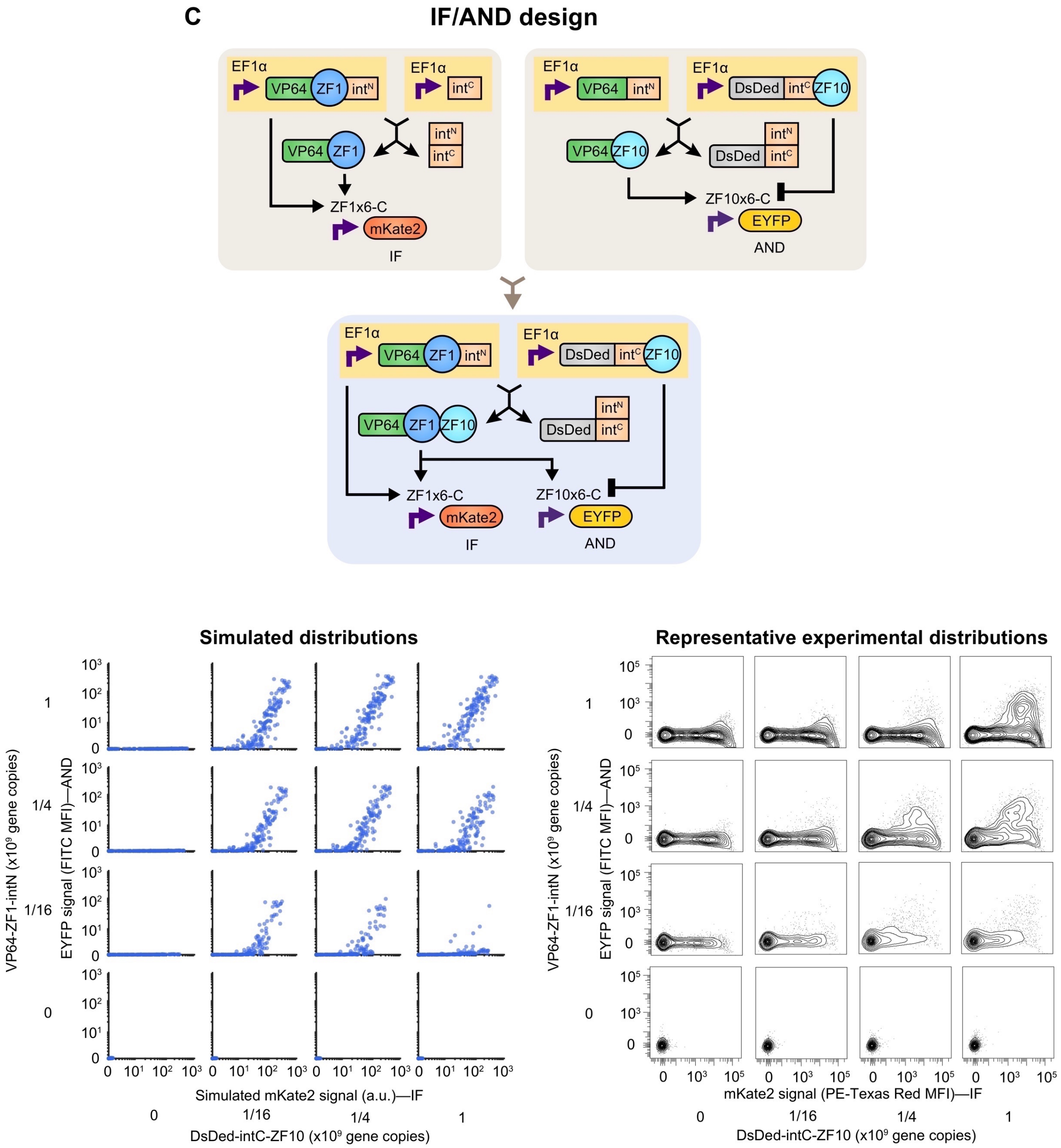

**Fig. S2** (**C**) IF/AND corresponding to **Fig. 2D**.

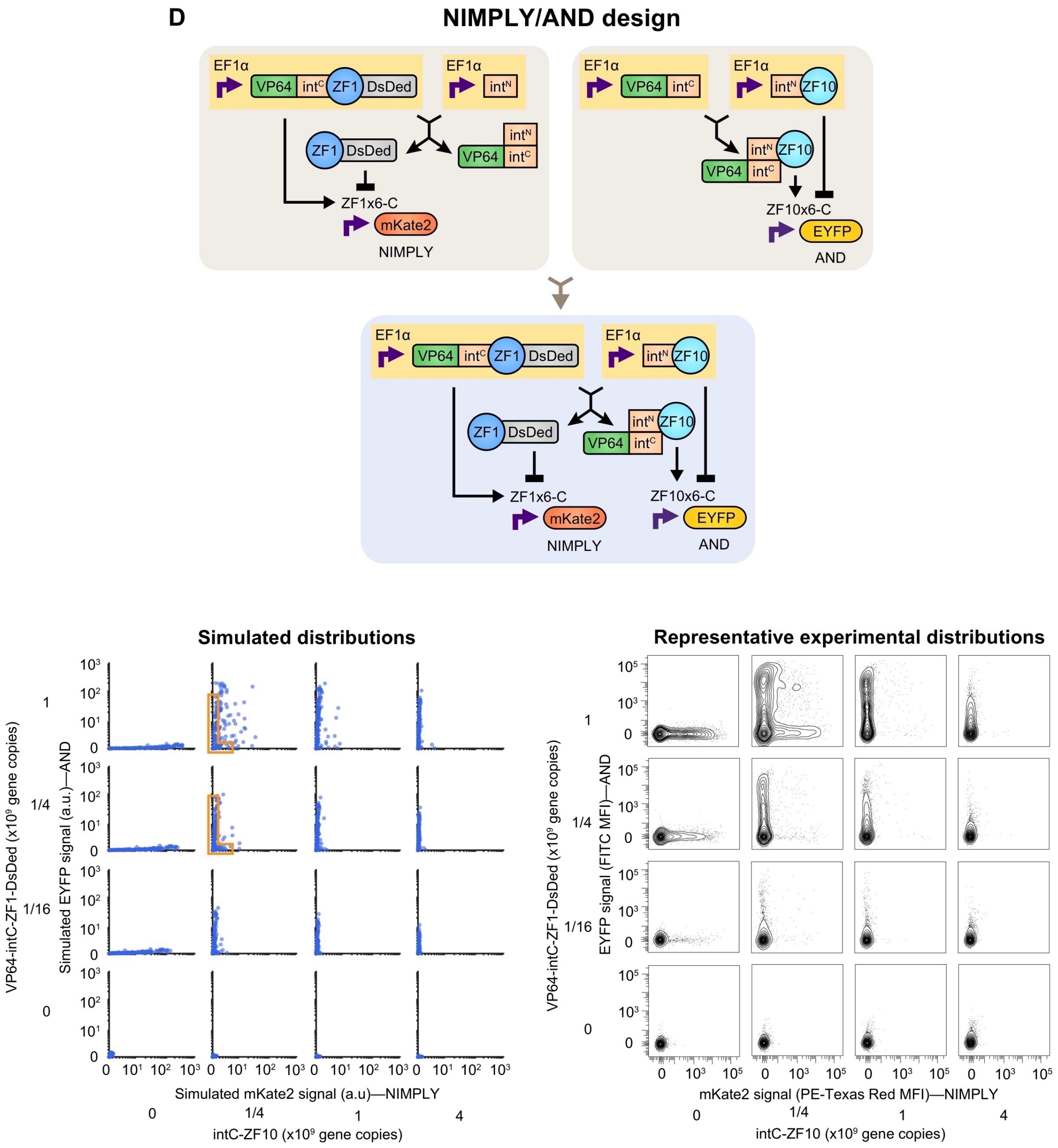

**Fig. S2** (**D**) NIMPLY/AND corresponding to **Fig. 2E**.

E

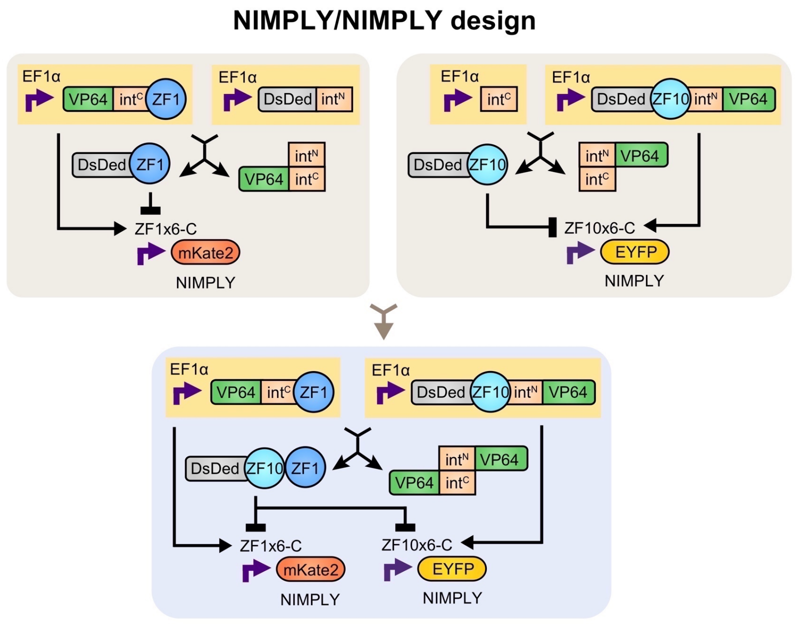

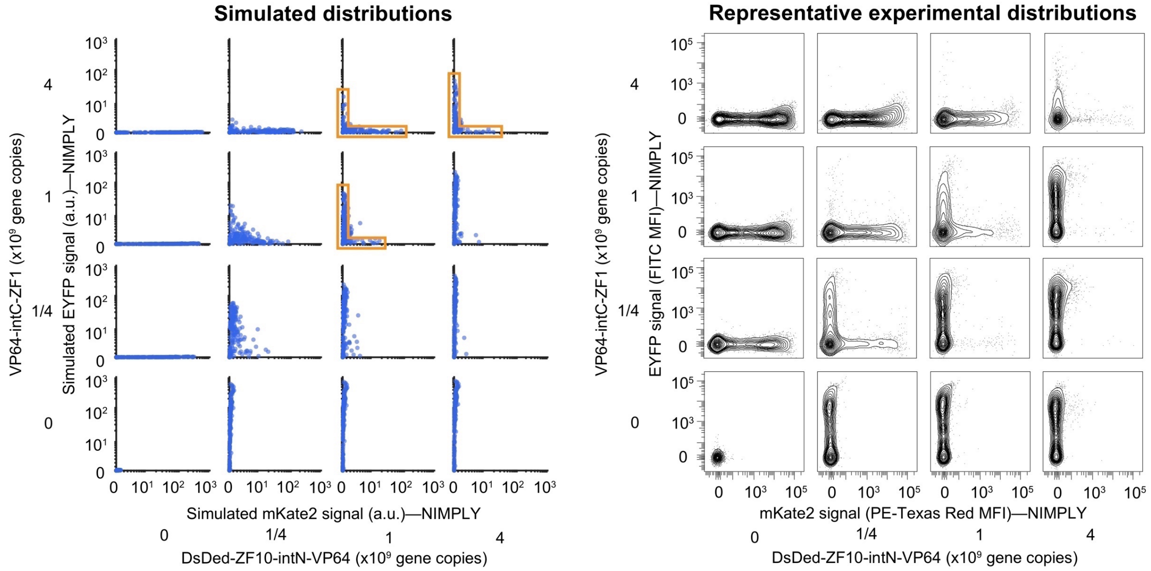

**Fig. S2** (**E**) NIMPLY/NIMPLY corresponding to **Fig. 2F**.

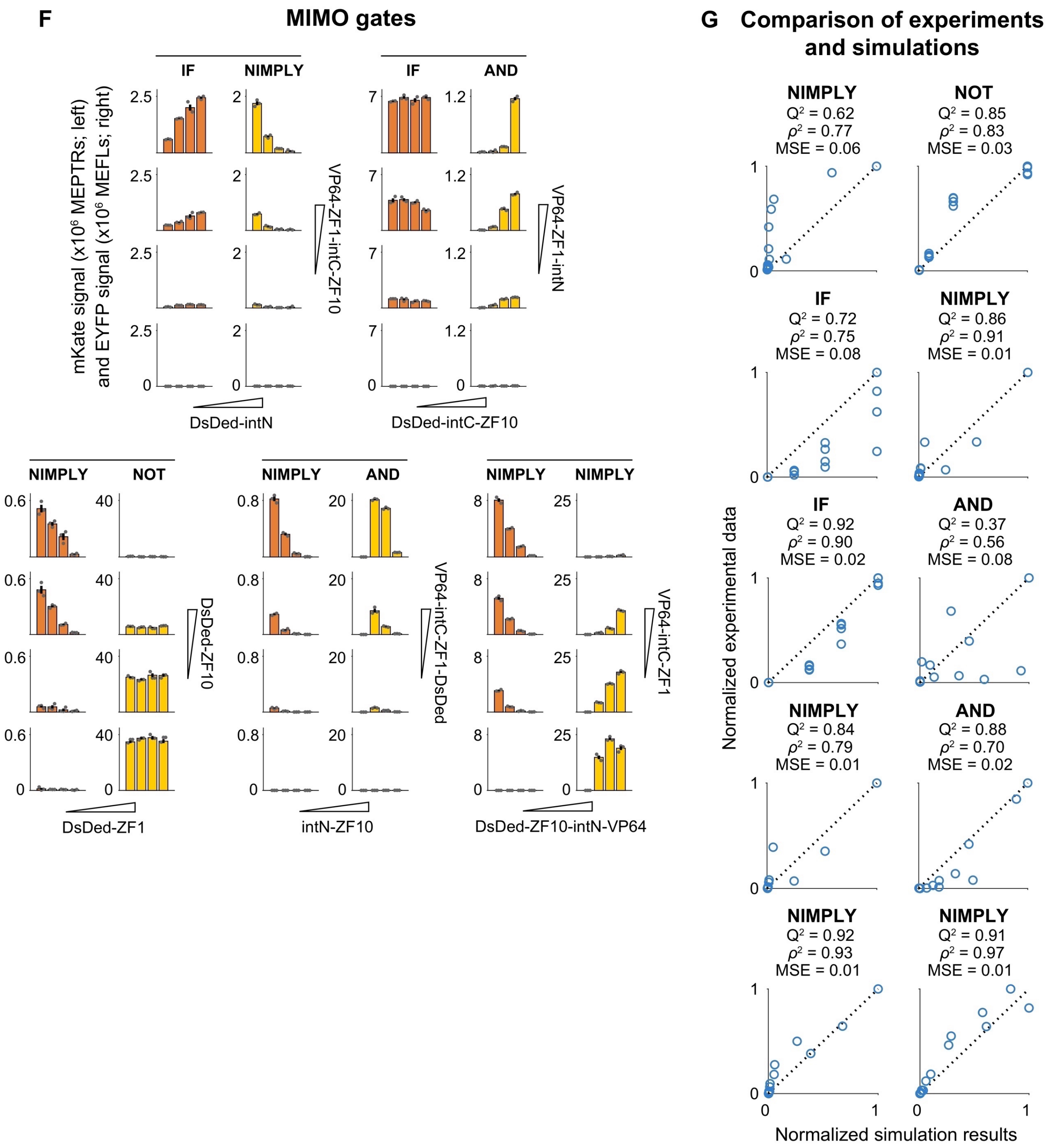

**Fig. S2** (**F**) Bar graphs represent the mean and S.E.M. of reporter signal corresponding to **Fig. 2B–F** for three biological replicates. (**G**) $Q^2$, $\rho^2$, and MSE comparing max-normalized simulated and observed signals. The comparisons indicate that in most cases, model simulations can explain the majority of the variance in the experimental data, and several cases have very close agreement (≥90% $Q^2$).

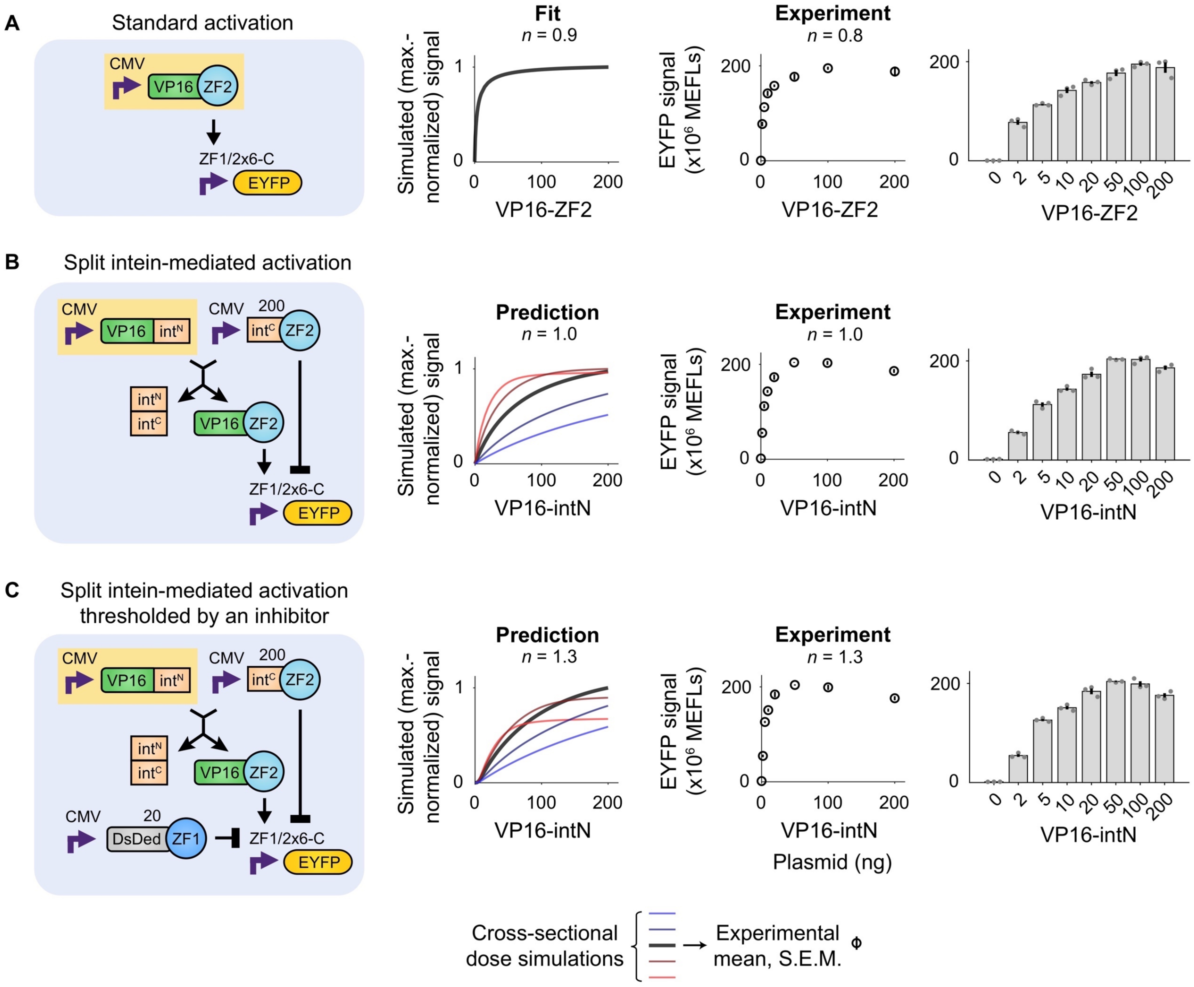

**Fig. S3. Analog signal processing** (**A–C**) Split intein-based programs were not substantially more ultrasensitive than a standard activation dose response; *n* is the exponent from the Hill equation fitted to the monotonically increasing portion of the data. We note that despite an apparent similarity of the split intein design in **Fig. S3B** and the RaZFa design in **Fig. 3C**, these systems have different ultrasensitivity outcomes. We interpret that this difference is due at least in part to decreased protein stability conferred by the intC domain, in that there would be less unspliced intC-ZF available than there would be non-dimerized FKBP-ZF for the same DNA dose, and thus less protein to act as an inhibitor in the former case. Data were collected in the same experiment. The mean and S.E.M. are plotted both as dot plots and bar graphs.

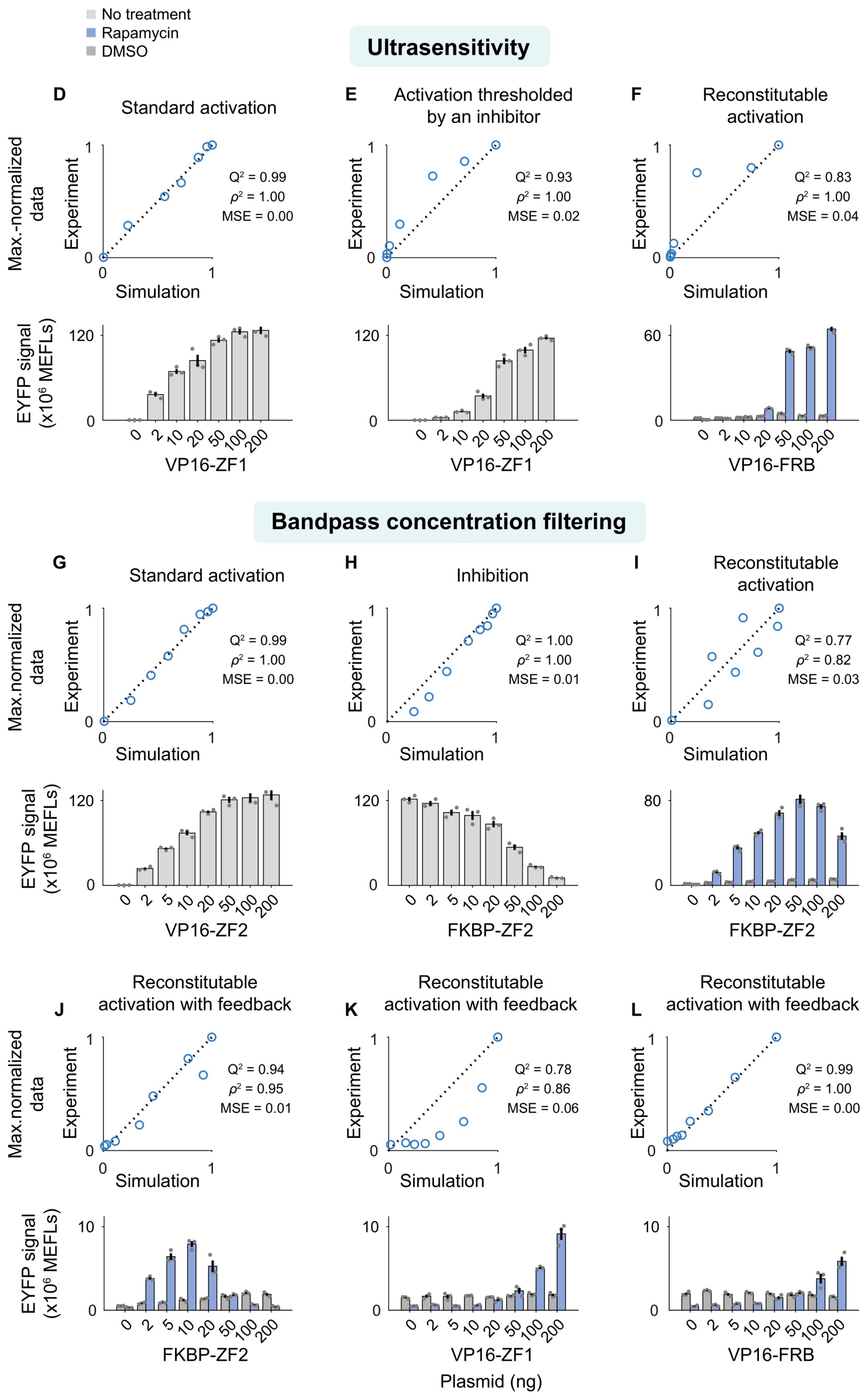

**Fig. S3** (**D–L**) Data correspond to the main figure: (**D–F**) **Fig. 3A–C**, (**G–H**) **Fig. 3D–E**, (**I**) **Fig. 3F**, and (**J–L**) **Fig. 3G–I**, and dose responses within each of these groupings were measured in the same experiment. Comparisons are of the max.-normalized simulated and observed mean reporter signal. The comparisons indicate that in each case, model simulations can explain the majority of the variance in the experimental data, and some cases have very close agreement (>90% $Q^2$).

**M**

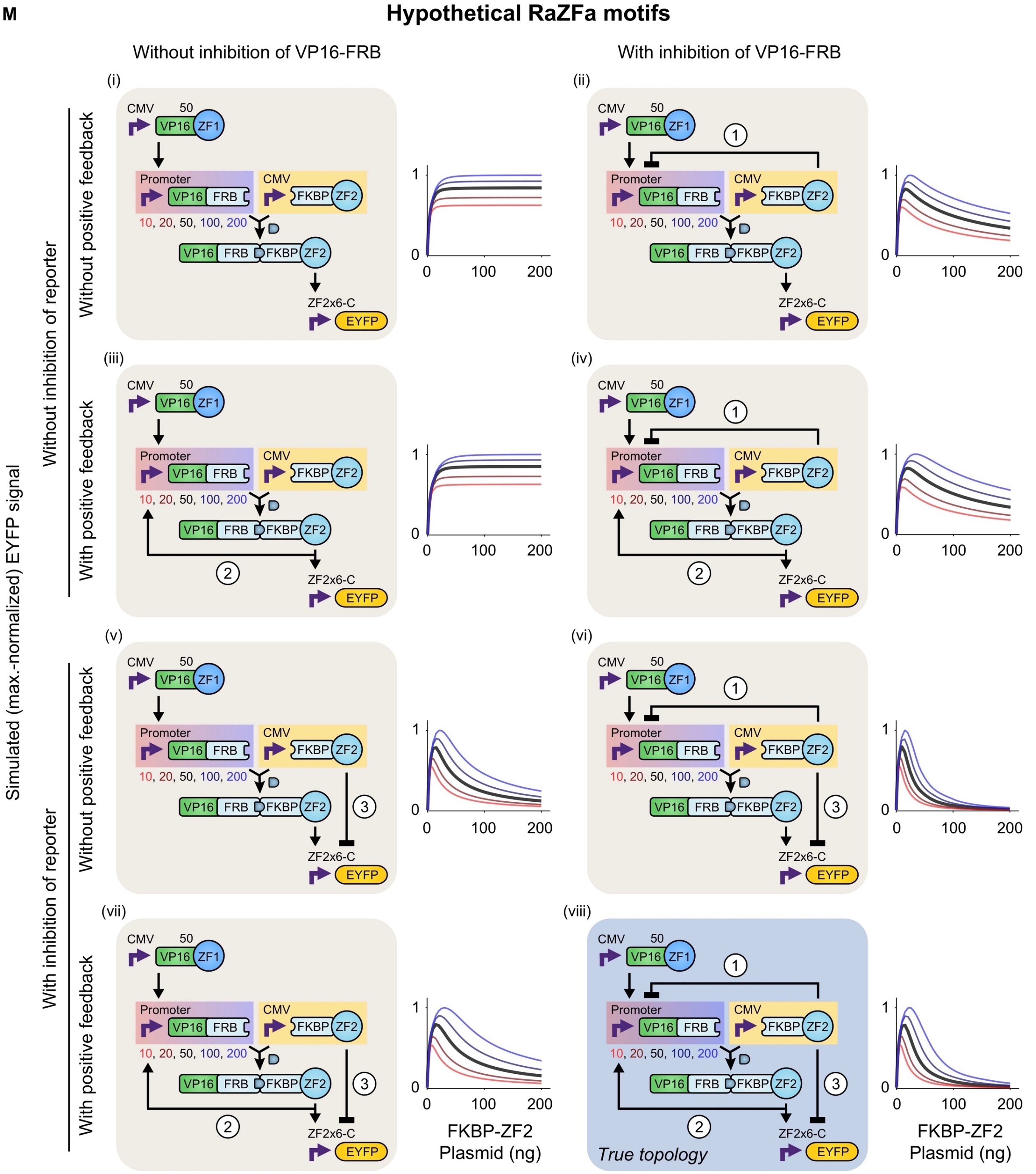

**Fig. S3** (**M**) Population simulations for hypothetical variations on the topology investigated in **Fig. 3G**. Three regulatory connections—(1) FKBP-ZF-mediated inhibition of VP16-FRB expression, (2) RaZFa-mediated induction of VP16-FRB expression, and (3) FKBP-ZF-mediated inhibition of reporter expression—are included or withheld in eight combinations. The lower-right scenario is the true scenario (the case which was experimentally evaluated in **Fig. 3G**) with all three connections included. The simulations indicate scenarios that are expected to produce bandpass filtering. Tight filtering is most evident in scenarios (vi) and (viii), which suggests that connections 1 and 3 contribute more than does 2 towards this design goal.

**A** **Simulated activation cascades**

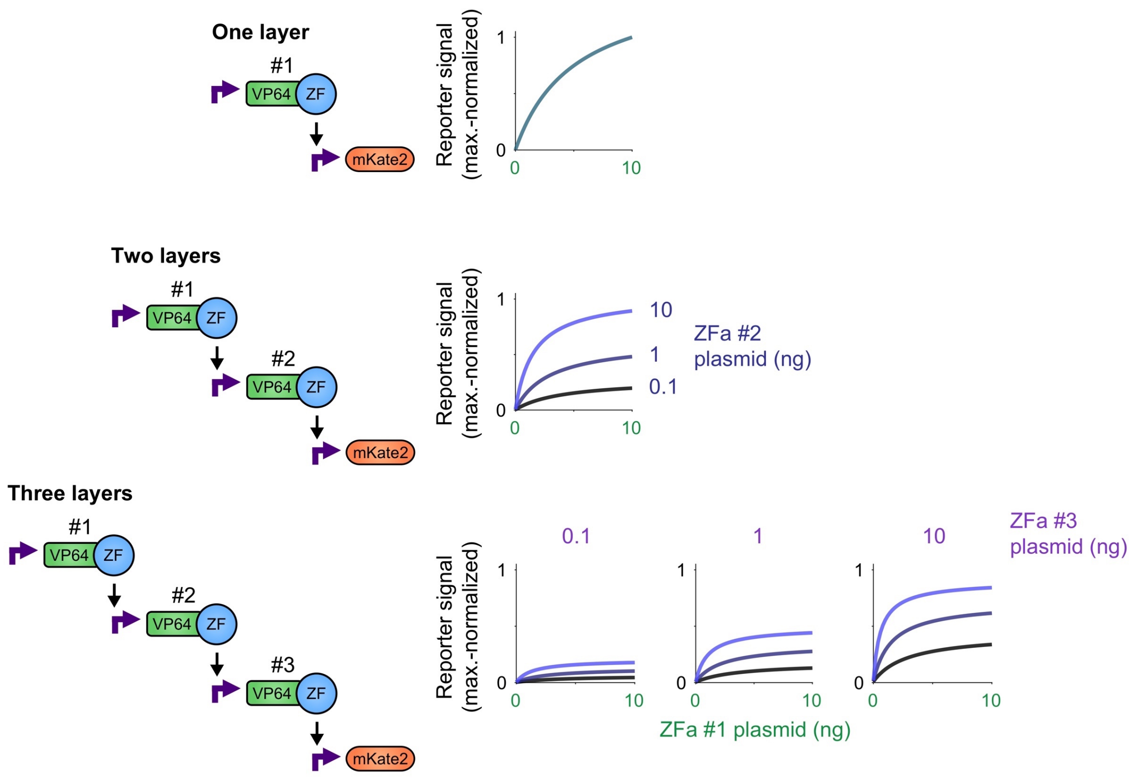

**Fig. S4. Receptor engineering and integration of ligand-responsive components.** (**A**) Homogeneous (single-cell) simulations suggest that different numbers of COMET transcriptional layers lead to altered dose responses (varied steepness and plateau of reporter expression) but do not produce a substantial leak in reporter signal. This absence of background signal propagation would support the implementation of high-performance multi-layer cascades. ZFa activities at target promoters are treated as orthogonal, and for simplicity all layers are assigned the same parameters using the base case of VP64-ZF1 at a ZF1x6-C promoter. Plasmid doses: ZFa #1 varied along the *x*-axis, ZFa #2 varied by color-coding, and ZFa #3 varied across plots.

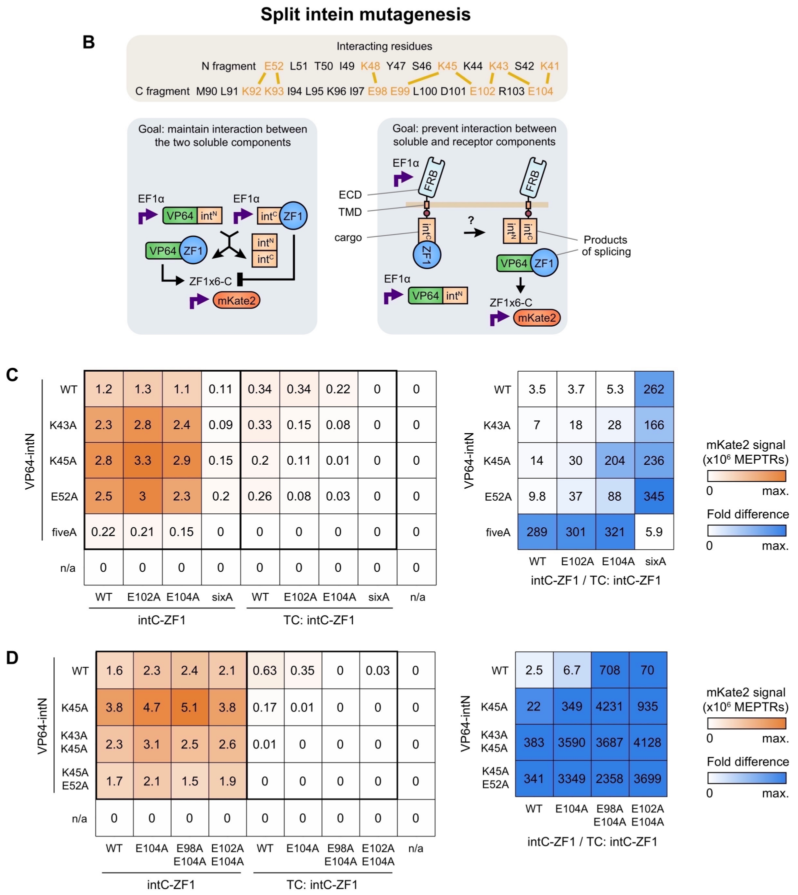

**Fig. S4** (**B–D**) A strategy was investigated for modulating split intein splicing efficiency. 11 glutamine and lysine residues across intN and intC were selected based on crystallographic evidence (*8*) for electrostatic involvement in the initial capture step in the mechanism for gp41-1 folding and splicing. We hypothesized that mutating residues to alanine (either individually or in combination, to reduce the number of electrostatic interactions) would decrease the likelihood of intN and intC folding upon coming into contact with each other—and by extension decrease TF reconstitution efficiency—and that this effect would be greater for intC as membrane-proximal cargo on a MESA target chain (TC) than as an intracellular protein. We reasoned that a sufficient differential effect between these two contexts would enable effective fusion of intC-ZF1 onto a TC, such that interactions with intracellular intN-containing components would occur only after proteolytic cargo release in ligand-induced receptor signaling. Abbreviations: ectodomain (ECD), transmembrane domain (TMD). (**B**) The cartoon illustrates the evaluation of reconstitution between two intracellular components (VP64-intN and intC-ZF1) and between an intracellular component and a receptor (VP64-intN and Rapa-MESA TC:intC-ZF1). Mutations were considered ideal if reporter signal was retained in the former scenario and not produced in the latter. (**C–D**) Several mutants were generated and tested (**C**), and based on these results, double mutants with K45A for intN and with E104A for intC were generated and tested (**D**). Heatmaps denote the mean reporter signal from three biological replicates (left heatmaps) and the fold difference in mean signal with intC-ZF1 vs. TC:intC-ZF1 (right heatmaps). The results indicate that the new pairings disrupted interactions with intN more for TC-fused intC than for intracellular intC. In some cases, pairings also produced up to several fold greater reporter signal in the intracellular context than did the WT-WT case, and in tandem with the reduction in signal in the receptor context there was a several thousand-fold context-dependent difference. High-performing variants were carried forward for further investigation (**Fig. S4E**). For the mutation of all 11 residues using the pairing of intN fiveA (K41A, K43A, K45A, K48A, E52A) and intC sixA (K92A, K93A, E98A, E99A, E102A, E104A) (**C**), reporter expression was not induced in either context. Thus, mutations can be used tune reconstitution efficiency from WT level to effectively none, and there exists an intermediate regime with a differential effect based on whether intC is TC cargo or intracellular.

**E** **Protease chain and ligand treatment**

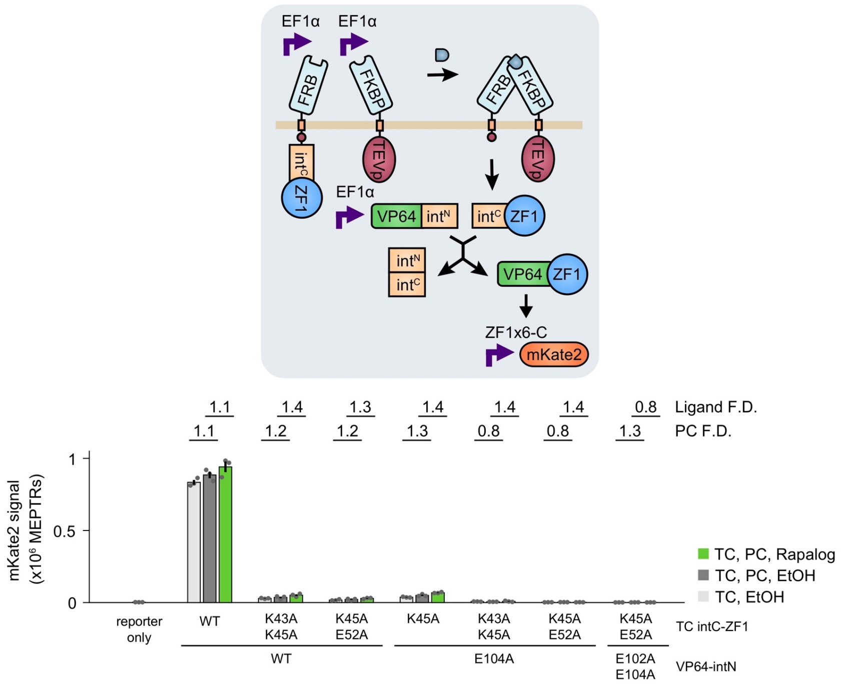

**Fig. S4** (**E**) A functional assay was conducted for ligand-inducible receptor signaling incorporating split inteins. A panel of intC and intN pairings from the mutagenesis assay was evaluated for compatibility with the MESA signaling mechanism by measuring reporter signal in three scenarios: (1) VP64-intN and TC:intC-ZF1 with vehicle (EtOH), (2) VP64-intN, TC:intC-ZF1, and MESA protease chain (PC) with vehicle, and (3) VP64-intN, TC:intC-ZF1, and PC with receptor ligand (rapalog). Outcomes are considered ideal if reporter signal is low in the first two scenarios and high in the third. However, we observed that for each pairing, reporter signal was similar regardless of PC co-expression or ligand treatment. This result does not support the ability of PC to cleave intC-containing cargo from the TC, and instead indicates that the conditions in which reporter signal was observed were due to residual interactions between intracellular intN and TC-bound intC partial-mutant variants. These designs were not carried forward, but variants with different intracellular linker lengths were evaluated in the next panel.

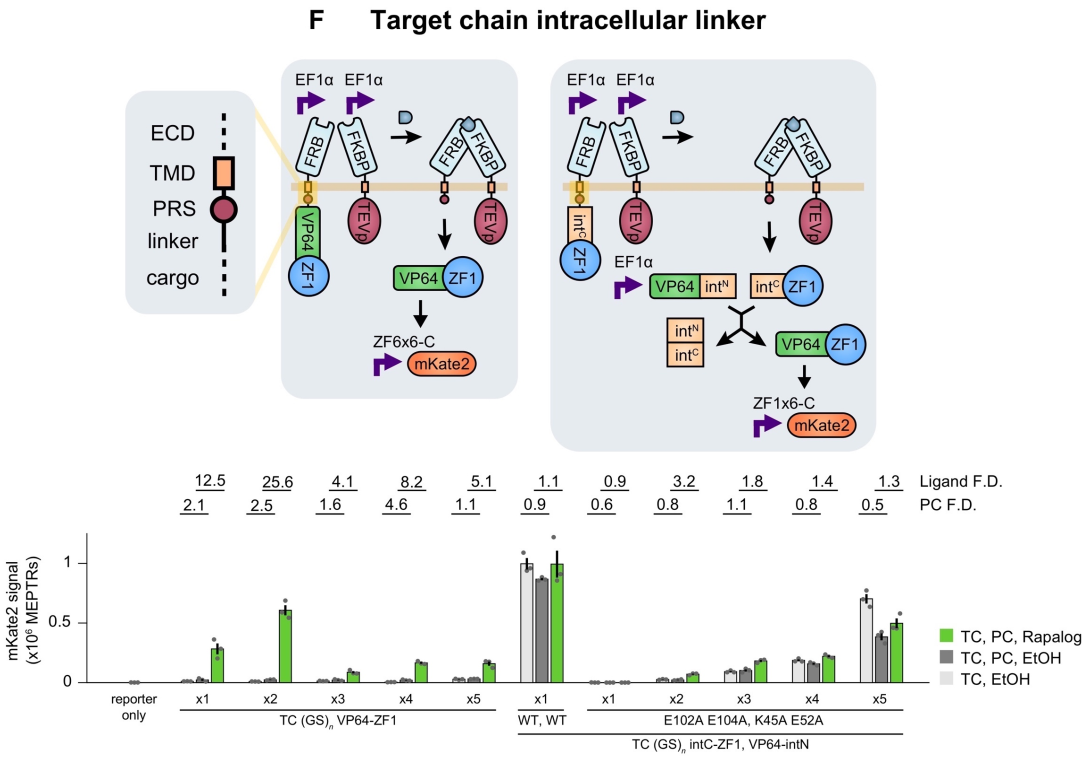

**Fig. S4** (**F**) Functional assay to assess the effect of TC intracellular linker length. We hypothesized that introducing more physical distance or geometric flexibility between the protease recognition sequence (PRS) and cargo would enable PC-mediated cleavage of TC:intC-ZF1. The base case TC linker had one glycine-serine (GS) repeat, and between one and five repeats were tested for mutant intC-ZF1 cargo co-expressed with mutant VP64-intN and for VP64-ZF1 cargo. For the case with intC-ZF1 cargo, reporter signal increased with increasing linker length; however, a signal increase occurred regardless of PC co-expression and ligand treatment. Therefore, modifications to the TC membrane-proximal region did not alleviate the inability of the PC to cleave these TCs. It is possible that the effect of increasing linker length is to make the membrane-proximal intC more accessible (resembling intracellular intC) to intracellular intN. For the case with VP64-ZF1 cargo, there was low reporter signal with PC and no ligand, and there was high reporter signal with PC and ligand, demonstrating for the first time that a ZFa can be effectively used on MESA. For increasingly long linkers, the reporter signal decreased. Based on these findings, we chose to use full transcription factors such as ZFa as cargo. Linkers for subsequent TCs (**Fig. 4**, **Fig. S4G,H**) contained one GS repeat.

### G Adapting MESA for inhibitory signaling

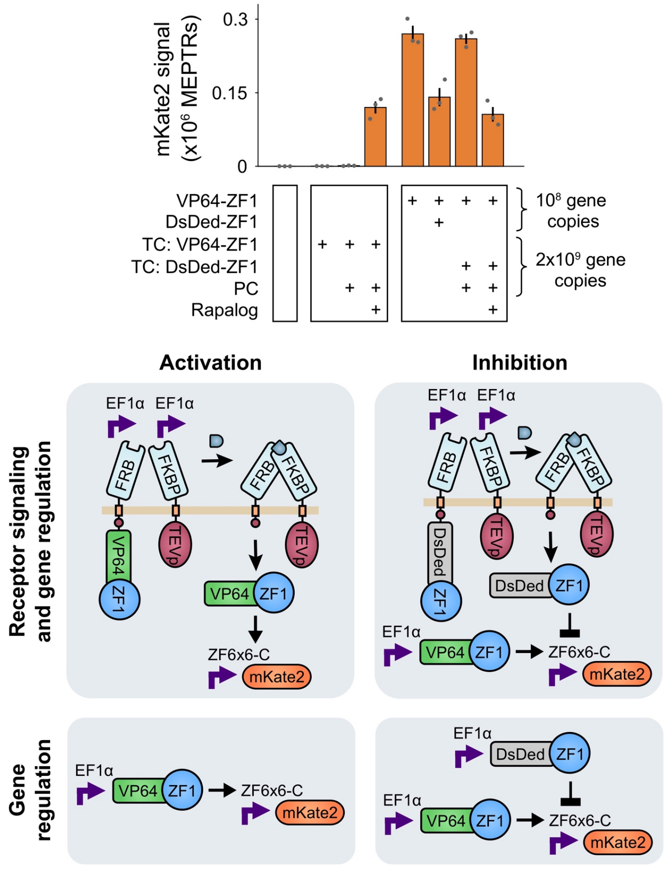

**Fig. S4** (**G**) MESA with DsDed-ZF1 cargo can ligand-inducibly signal to inhibit target gene expression. In this assay, treatment with EtOH or rapalog was applied both at the time of transfection and at the time of media change (rather than only at the latter) to promote inhibitory signaling upon expression of the receptor, analogously to inhibition that could take place upon expression of the intracellular inhibitor. In summary, for **Fig. S4B–G,** we pursued a strategy of mutating charged residues at the intN and intC fragment interface to counter their association. Functional assays identified mutants that prevented intC-ZF1 cargo from reconstituting with soluble VP64-intN and that did not prevent soluble intC-ZF1 from doing so, suggesting that cargo could be isolated from downstream activity without precluding activity. With certain mutations, it was also possible to tune soluble intC-ZF1 and VP64-intN splicing down to zero (**Fig. S4B–D**). However, full MESA signaling was not observed with intC cargo (**Fig. S4E,F**), but since inducible signaling did prove to be effective with ZFa cargo, we selected this configuration.

### H Fig. 4 data with linear y-axis scaling

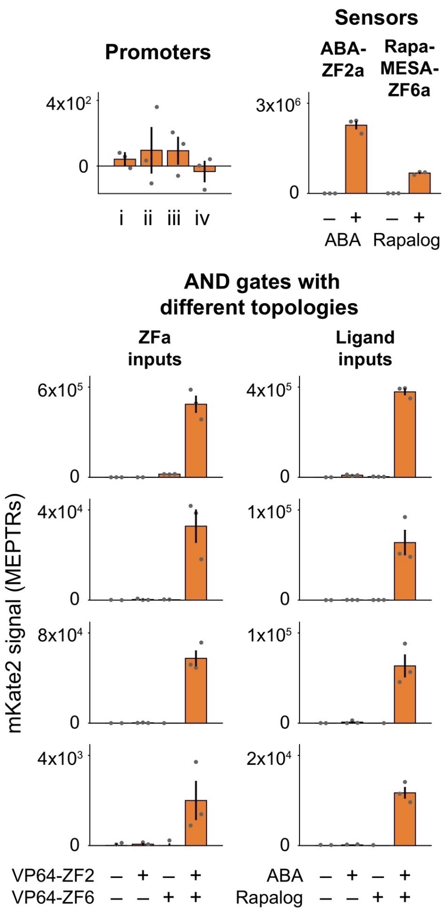

**Fig. S4** (**H**) Data correspond to **Fig. 4** with linear *y*-axis scaling. Split intein-containing components were placed under the control of COMET inducible promoters and incorporated downstream of either constitutive TFs or a ligand-responsive TF and receptor. Data were collected in the same experiment.

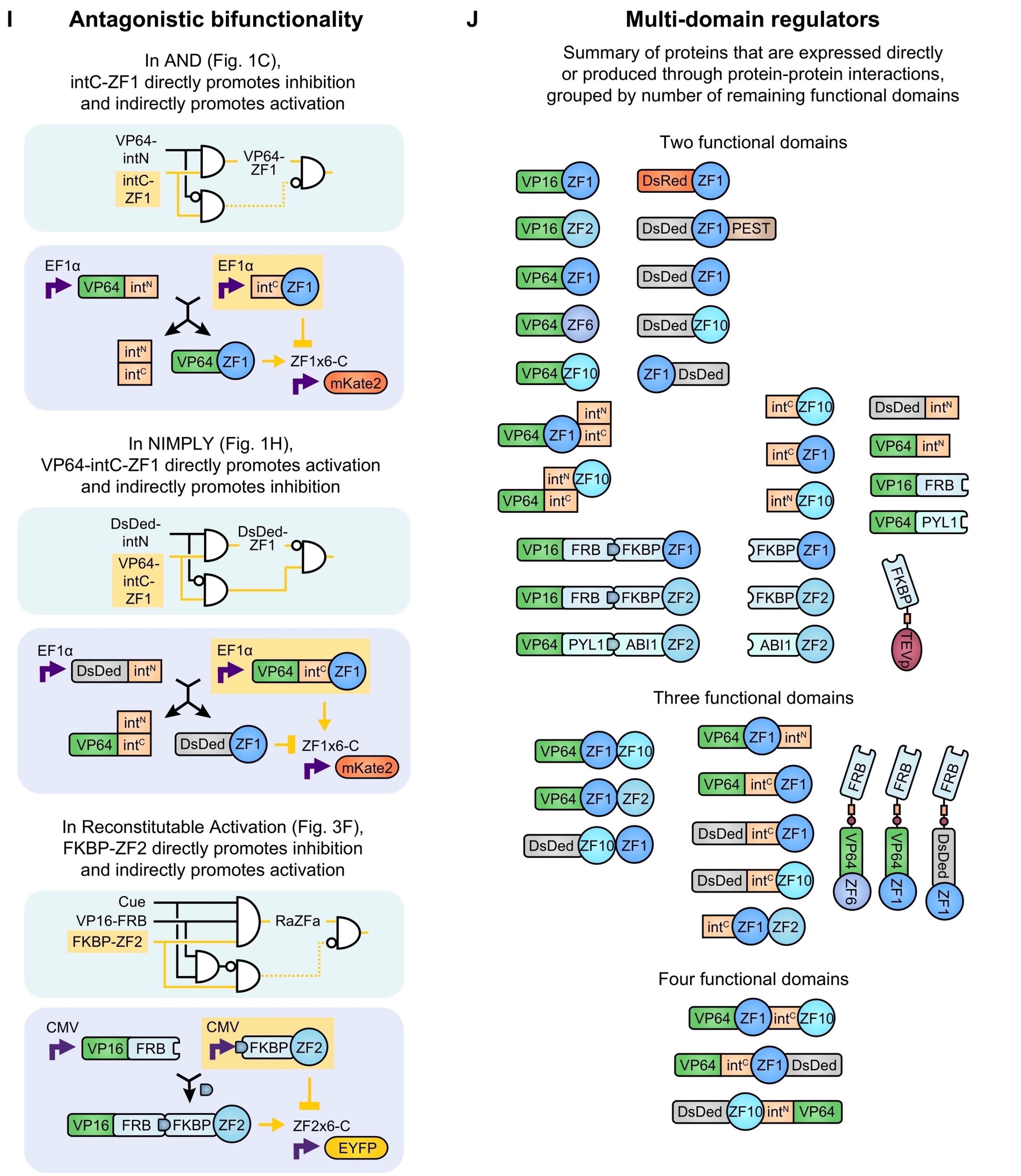

**Fig. S4** (**I**) Selected examples of antagonistic bifunctionality, in which a component exerts opposing effects. (**J**) Depiction of validated multi-domain regulators used (or generated through reactions) in this study. Even a relatively small set of domains can be combined in numerous ways, such that multiple activities can be concisely encoded in TFs. An additional note: after small molecule-reconstituted TFs dimerize, the small molecule-binding domains are not involved in subsequent interactions, and because the remaining functional domains (those that can carry out subsequent activities) are the AD and ZF, the small molecule-reconstituted TFs are depicted in the category for two functional domains.

### Table S1. Plasmids used in this study

| ID | Promoter | Gene | Figs. |
|---|---|---|---|
| Constitutive FPs | | | |
| pPD193 [i] | CMV | EBFP2 | S1, 3, S3, 4, S4 |
| pPD133 [i] | CMV | EYFP | S1, 3, S3 |
| pJM409 | CMV | mKate2 | S1 |
| pJM413 | EF1α | mKate2 | 1, S1, 2, S2, 4, S4 |
| pJM451 | EF1α | EBFP2-P2A-BlastR | S1 |
| pJM500 | EF1α | EYFP | S1, 2, S2 |
| pJM501 | EF1α | EBFP2 | 1, S1, 2, S2, S4 |
| Reporter FPs | | | |
| pPD270 [i] | ZF1/2x6-C | EYFP | 3, S3 |
| pPD290 [i] | ZF1x6-C | EYFP | S1 |
| pPD545 [i] | ZF2x6-C | EYFP | 3, S3 |
| pJM410 | ZF1x6-C | mKate2 | 1, S1, 2, S2, 4, S4 |
| pJM502 | ZF1x6-C | EYFP | S1 |
| pJM584 | ZF2x6-C | mKate2 | 4, S4 |
| pJM585 | ZF6x6-C | mKate2 | 4, S4 |
| pJM587 | ZF10x6-C | EYFP | 2, S2 |
| pJM597 | ZF(2/6)x3 | mKate2 | 4, S4 |
| TFs (main group) | | | |
| pJB002 | CMV | 3xFLAG-NLS-DsDed-ZF1 | 3, S3 |
| pPD100 [i] | CMV | 3xFLAG-NLS-VP16-ZF1 | S1, 3, S3 |
| pPD189 [i] | CMV | 3xFLAG-NLS-VP16-ZF2 | 3, S3 |
| pPD303 [i] | CMV | 3xFLAG-NLS-VP64-ZF1 | S1 |
| pJM465 | EF1α | 3xFLAG-NLS-VP64-ZF1 | S1 |
| pJM466 | EF1α | 3xFLAG-NLS-VP16-ZF1 | S1 |
| pJM503 | EF1α | 3xFLAG-NLS-ZF1 | 1, S1 |
| pJM505 | EF1α | 3xFLAG-NLS-DsRed-ZF1 | S1 |
| pJM506 | EF1α | 3xFLAG-NLS-DsDed-ZF1 | 1, S1, 2, S2 |
| pJM507 | EF1α | 3xFLAG-NLS-DsDed-ZF10 | 1, S1, 2, S2 |
| pJM512 | EF1α | 3xFLAG-NLS-VP64-ZF1 | 1, S1, 2, S2 |
| pJM520 | EF1α | 3xFLAG-NLS-VP64-ZF10 | 1, S1, 2, S2 |
| pJM553 | ZF10x6-C | 3xFLAG-NLS-DsDed-ZF1-PEST | 1, S1, 2, S2 |
| pJM574 | ZF2x6-C | 3xFLAG-NLS-DsDed-ZF1 | 4, S4 |
| pJM580 | ZF6x6-C | 3xFLAG-NLS-VP64-ZF1 | 4, S4 |
| pJM582 | EF1α | 3xFLAG-NLS-VP64-ZF2 | 4, S4 |
| pJM583 | EF1α | 3xFLAG-NLS-VP64-ZF6 | 4, S4 |
| Small molecule-responsive TFs | | | |
| pJB001 | CMV | 3xFLAG-NLS-FKBP-ZF2 | 3, S3 |
| pJB003 | ZF1/2x6-C | 3xFLAG-NLS-VP16-FRB | 3, S3 |
| pPD341 [i] | CMV | 3xFLAG-NLS-FKBP-ZF1 | 3, S3 |
| pPD353 [i] | CMV | 3xFLAG-NLS-VP16-FRB | 3, S3 |
| pPD1122 | CMV | 3xFLAG-NES-PYL1-VP64 | 4, S4 |
| pKD011 | CMV | 3xFLAG-NLS-ZF2-ABI | 4, S4 |
| TFs with WT split inteins | | | |
| pJB004 | CMV | 3xFLAG-NLS-VP16-intN | S3 |
| pJB005 | CMV | 3xFLAG-NLS-intC-ZF2 | S3 |

| | | | |
|---|---|---|---|
| pJM516 | EF1α | intC-ZF1-NLS-HA | 1, S1, S4 |
| pJM522 | EF1α | intC-ZF10-NLS-HA | 1, S1 |
| pJM524 | EF1α | 3xFLAG-NLS-VP64-intC-ZF1-NLS-HA | 1, 2, S2 |
| pJM525 | EF1α | 3xFLAG-NLS-DsDed-intC-ZF1-NLS-HA | 1, S1 |
| pJM529 | EF1α | 3xFLAG-NLS-DsDed-intC-ZF10-NLS-HA | 2, S2 |
| pJM554 | EF1α | 3xFLAG-NLS-VP64-intN | 1, S1, S4 |
| pJM555 | EF1α | 3xFLAG-NLS-DsDed-intN | 1, S1, 2, S2 |
| pJM571 | ZF2x6-C | 3xFLAG-NLS-VP64-intN | 4, S4 |
| pJM572 | ZF6x6-C | intC-ZF1-NLS-HA | 4, S4 |
| pJM573 | ZF6x6-C | 3xFLAG-NLS-DsDed-intC-ZF1-NLS-HA | 4, S4 |
| pJM575 | ZF6x6-C | 3xFLAG-NLS-DsDed-ZF10 | 4, S4 |
| pJM576 | ZF2x6-C | 3xFLAG-NLS-VP64-intC-ZF1-NLS-HA | 4, S4 |
| pJM577 | ZF6x6-C | 3xFLAG-NLS-DsDed-intN | 4, S4 |
| pJM588 | EF1α | 3xFLAG-NLS-VP64-intC-ZF1-DsDed-NLS-HA | 2, S2 |
| pJM589 | EF1α | intN-ZF10-NLS-HA | 2, S2 |
| pJM590 | EF1α | 3xFLAG-NLS-VP64-ZF1-intC-ZF10-NLS-HA | 2, S2 |
| pJM591 | EF1α | 3xFLAG-NLS-VP64-ZF1-intN | 2, S2 |
| pJM592 | EF1α | 3xFLAG-NLS-DsDed-ZF10-intN-VP64 | 2, S2 |
| pJM595 | ZF6x6-C | intC-ZF1-ZF2-NLS-HA | 4, S4 |
| TFs with mutant split inteins | | | |
| pJM556 | EF1α | 3xFLAG-NLS-VP64-intN_K43A | S4 |
| pJM557 | EF1α | 3xFLAG-NLS-VP64-intN_K45A | S4 |
| pJM558 | EF1α | 3xFLAG-NLS-VP64-intN_E52A | S4 |
| pJM559 [ii] | EF1α | 3xFLAG-NLS-VP64-intN_fiveA | S4 |
| pJM560 | EF1α | intC-ZF1-NLS-HA_E102A | S4 |
| pJM561 | EF1α | intC-ZF1-NLS-HA_E104A | S4 |
| pJM562 [iii] | EF1α | intC-ZF1-NLS-HA_sixA | S4 |
| pJM563 | EF1α | 3xFLAG-NLS-VP64-intN_K43A_K45A | S4 |
| pJM564 | EF1α | 3xFLAG-NLS-VP64-intN_K45A_E52A | S4 |
| pJM565 | EF1α | intC-ZF1-NLS-HA_E98A_E104A | S4 |
| pJM566 | EF1α | intC-ZF1-NLS-HA_E102A_E104A | S4 |
| Receptors (main group) | | | |
| pJM600 | EF1α | PC 3xFLAG-FKBP-FGFR4-TEVp | 4, S4 |
| pJM612 | EF1α | TC 3xFLAG-FRB-FGFR4-PRS(M)-(GS)x1-NLS-VP64-ZF1 | S4 |
| pJM621 | EF1α | TC 3xFLAG-FRB-FGFR4-PRS(M)-(GS)x1-NLS-DsDed-ZF1 | S4 |
| pJM671 | EF1α | TC 3xFLAG-FRB-FGFR4-PRS(M)-(GS)x1-NLS-VP64-ZF6 | 4, S4 |
| Receptor with WT split intein and (GS)x1 linker | | | |
| pJM616 | EF1α | TC 3xFLAG-FRB-FGFR4-PRS(M)-(GS)x1-intC-ZF1-NLS | S4 |
| Receptors with mutant split inteins and (GS)x1 linker | | | |
| pJM626 | EF1α | TC 3xFLAG-FRB-FGFR4-PRS(M)-(GS)x1-intC-ZF1-NLS_E102A | S4 |
| pJM627 | EF1α | TC 3xFLAG-FRB-FGFR4-PRS(M)-(GS)x1-intC-ZF1-NLS_E104A | S4 |
| pJM628 [iii] | EF1α | TC 3xFLAG-FRB-FGFR4-PRS(M)-(GS)x1-intC-ZF1-NLS_sixA | S4 |
| pJM633 | EF1α | TC 3xFLAG-FRB-FGFR4-PRS(M)-(GS)x1-intC-ZF1-NLS_E98A_E104A | S4 |
| pJM634 | EF1α | TC 3xFLAG-FRB-FGFR4-PRS(M)-(GS)x1-intC-ZF1-NLS_E102A_E104A | S4 |
| Receptors with extended (GS) linkers | | | |
| pJM614 | EF1α | TC 3xFLAG-FRB-FGFR4-PRS(M)-(GS)x2-NLS-VP64-ZF1 | S4 |
| pJM657 | EF1α | TC 3xFLAG-FRB-FGFR4-PRS(M)-(GS)x3-NLS-VP64-ZF1 | S4 |
| pJM658 | EF1α | TC 3xFLAG-FRB-FGFR4-PRS(M)-(GS)x4-NLS-VP64-ZF1 | S4 |
| pJM659 | EF1α | TC 3xFLAG-FRB-FGFR4-PRS(M)-(GS)x5-NLS-VP64-ZF1 | S4 |

| | | | |
|---|---|---|---|
| pJM661 | EF1α | TC 3xFLAG-FRB-FGFR4-PRS(M)-(GS)x2-intC-ZF1-NLS_E102A_E104A | S4 |
| pJM662 | EF1α | TC 3xFLAG-FRB-FGFR4-PRS(M)-(GS)x3-intC-ZF1-NLS_E102A_E104A | S4 |
| pJM663 | EF1α | TC 3xFLAG-FRB-FGFR4-PRS(M)-(GS)x4-intC-ZF1-NLS_E102A_E104A | S4 |
| pJM664 | EF1α | TC 3xFLAG-FRB-FGFR4-PRS(M)-(GS)x5-intC-ZF1-NLS_E102A_E104A | S4 |
| Other | | | |
| pPD005 [i] | CMV | n/a | 1, S1, 2, S2, 3, S3, 4, S4 |

[i] Generated and used in the COMET study (*10*)
[ii] Mutations in intN fiveA: K41A, K43A, K45A, K48A, E52A
[iii] Mutations in intC sixA: K92A, K93A, E98A, E99A, E102A, E104A

### Table S2. Model parameters

| Symbol | Description | Value [i,ii] | Source |
|---|---|---|---|
| $b_1$ | Basal transcription at ZF1x6-C promoter | 0.08 U | COMET |
| $m_1$ | Max. induction for CMV-driven VP16-ZF1 at ZF1x6-C promoter | 33 | COMET |
| $w_1$ | Steepness for CMV-driven VP16-ZF1 at ZF1x6-C promoter | 0.036 | Fitted here |
| $m_{1E64}$ | Max. induction by EF1α-driven VP64-ZF1 at ZF1x6-C promoter | 52 | Fitted here |
| $w_{1E64}$ | Steepness for EF1α-driven VP64-ZF1 at ZF1x6-C promoter | 0.192 | Fitted here |
| $b_2$ | Basal transcription at ZF2x6-C promoter | 0.25 U | COMET |
| $m_2$ | Max. induction by CMV-driven VP16-ZF2 at ZF2x6-C promoter | 54 | COMET |
| $w_2$ | Steepness for CMV-driven VP16-ZF2 at ZF2x6-C promoter | 0.082 | Fitted here |
| $b_H$ | Basal transcription at ZF1/2x6-C promoter | $b_1$ | Assumed |
| $m_{1H}$ | Max. induction by CMV-driven VP16-ZF1 at ZF1/2x6-C promoter | $m_1$ | Assumed |
| $m_{2H}$ | Max. induction by CMV-driven VP16-ZF2 at ZF1/2x6-C promoter | $m_2$ | Assumed |
| $w_{1H}$ | Steepness for CMV-driven VP16-ZF1 at ZF1/2x6-C promoter | 0.072 | Fitted here |
| $w_{2H}$ | Steepness for CMV-driven VP16-ZF2 at ZF1/2x6-C promoter | 0.170 | Fitted here |
| $b_{10}$ | Basal transcription at ZF10x6-C promoter | 0.01 U | COMET |
| $m_{10E64}$ | Max. induction by EF1α-driven VP64-ZF10 at ZF10x6-C promoter | $m_{1E64}$ | Assumed |
| $w_{10E64}$ | Steepness for EF1α-driven VP64-ZF10 at ZF10x6-C promoter | $w_{1E64}$ | Assumed |
| $l$ | Start of loss of cooperativity (0.5 in COMET) | 0 | Adjusted here |
| $u$ | End of loss of cooperativity (2 in COMET) | 1.5 | Adjusted here |
| $w_{r1E64}$ | Steepness for DsDed-ZF1 inhibition of EF1α-driven VP64-ZF1 at ZF1x6-C | 4 * $w_{r1E64}$ | Definition |
| $w_{r10E64}$ | Steepness for DsDed-ZF10 inhibition of EF1α-driven VP64-ZF10 at ZF10x6-C | 4 * $w_{r10E64}$ | Definition |
| $w_{r1H}$ | Steepness for DsDed-ZF1 inhibition of CMV-driven VP64-ZF1 at ZF1/2x6-C | 4 * $w_{r1H}$ | Definition |
| rec | Reconstitution of split TFs (fitted based on split inteins; also applied to RaZFa) | 0.34 $U^{-1}$ $h^{-1}$ | Fitted here |
| $k_{txCMV}$ | Transcription at CMV promoter | 1 U $h^{-1}$ | Default |
| $k_{txEF1a}$ | Transcription at EF1α promoter | 1 U $h^{-1}$ | Default |
| $k_{txZF}$ | Transcription at COMET promoters | 1 U $h^{-1}$ | Assumed |
| $k_{tl}$ | Translation | 1 $h^{-1}$ | Default |
| $k_{degR}$ | Degradation of RNA | 2.7 $h^{-1}$ | COMET |
| $k_{degZFP}$ | Degradation of TF protein (default) | 0.35 $h^{-1}$ | COMET |
| $k_{degZFP\_PEST}$ | Degradation of PEST-tagged TF protein | 0.7 $h^{-1}$ | Assumed |
| $k_{degintC}$ | Degradation of intC-containing TF protein | 1.3 $h^{-1}$ | Fitted here |
| $k_{degRep}$ | Degradation of reporter protein | 0.029 $h^{-1}$ | COMET |

[i] The symbol U indicates arbitrary units.
[ii] Parameters for which no units are indicated are unitless.
